# Design of a genetically programmable and customizable protein scaffolding system for the hierarchical assembly of robust, functional macroscale materials

**DOI:** 10.1101/2024.05.02.592261

**Authors:** Ruijie Zhang, Sun-Young Kang, Francois Gaascht, Eliana L. Pena, Claudia Schmidt- Dannert

**Affiliations:** Department of Biochemistry, Molecular Biology & Biochemistry, University of Minnesota, Minneapolis, MN 55455, USA; BioTechnology Institute, University of Minnesota, St. Paul, MN 55108, USA

**Keywords:** Self-assembly, protein nanomaterials, biomanufacturing, biomaterials, biocatalysis, synthetic biology

## Abstract

Inspired by the properties of natural protein-based biomaterials, protein nanomaterials are increasingly designed with natural or engineered peptides, or with protein building blocks. Very few examples describe the design of functional protein-based materials for biotechnological applications that can be readily manufactured, are amenable to functionalization, and exhibit robust assembly properties for macroscale material formation. Here, we designed a protein-scaffolding system that self-assembles into robust, macroscale materials suitable for cell-free applications. By controlling the co-expression in *E. coli* of self-assembling scaffold building blocks with and without modifications for covalent attachment of cross-linking cargo proteins, hybrid scaffolds with spatially organized conjugation sites are overproduced that can be readily isolated. Cargo proteins, including enzymes, are rapidly cross-linked onto scaffolds for the formation of functional materials. We show that these materials can be used for the cell-free operation of a co-immobilized two-enzyme reaction and that the protein material can be recovered and reused. We believe that this work will provide a versatile platform for the design and scalable production of functional materials with customizable properties and the robustness required for biotechnological applications.

## INTRODUCTION

The chemical and structural variability of proteins make them ideal building blocks for the design of functional materials that can be genetically programmed and recombinantly manufactured. Numerous protein-based materials are known in nature; many with functions and multi-scale assembly properties that are unmatched by human-made, synthetic materials. Materials like silk, elastin or collagen therefore have inspired the design of synthetic peptide-based materials that have long been studied for biomedical applications^1–4^. Increasingly, self-assembling protein nanomaterials are created from natural or designed peptides and protein domains by leveraging a fast-growing knowledgebase on such building blocks as advances in de novo protein design^5–9^. Significantly fewer examples describe the design of functional protein-based materials for biotechnological applications that can be readily manufactured, are amenable to functionalization and exhibit robust assembly properties for macroscale material formation. Towards the fabrication of such materials, we previously showed that the bacterial microcompartment shell protein EutM^10^ self-assembles into highly robust, hexameric two-dimensional scaffolds that can be readily produced and isolated from recombinant *E. coli*^11^. Scaffold building blocks can be genetically modified with a SpyCatcher domain to allow for rapid covalent attachment of SpyTag modified cargo proteins to scaffolds via isopeptide bond formation^12^. We then created a toolbox of EutM homologs for the assembly of scaffolds, including hybrid scaffolds from two homologs, with different morphologies and surface properties^13^. Finally, we used our scaffolds for enzyme immobilization and demonstrated that scaffold attachment increased enzyme stability and efficiency of an enzyme cascade reaction^12,14^.

In this work, we sought to further develop our versatile scaffolding system for the programmable, hierarchical assembly of scaffolds into macroscale, functional materials as illustrated in **Fig. 1**. To achieve higher-order assembly, we expanded our previously used SpyTag/SpyCatcher (SpyT/SpyC hereafter) approach covalent scaffold attachment of cargo proteins with the orthogonal SnoopTag/SnoopCatcher (SnoopT/SnoopC hereafter) system for isopeptide bond formation^15^. Fusion of these orthogonal Tag/Catcher sequences to the N– and C-termini of EutM scaffold building blocks and cargo-proteins of choice will then allow cross-linking of scaffolds between cognate Tag and Catcher moieties. Considering our goal of creating functional materials, we envision that bioconjugated cargo proteins can be structural, cross-linking and/or functional (e.g. enzymes for biocatalysis) components of the formed material where their spatial arrangement is controllable by the density of Tag/Catcher attachment points on the scaffolds. To proto-type our envisioned platform, we first designed and characterized scaffold building blocks that upon bioconjugation with double-tagged GFP as model cargo protein drive the *in vitro* self-organization of higher ordered assemblies. A similar approach has recently been used for the 3D-assembly of S-layer sheets^16^. For ease of future scaffold fabrication and to control the density of attachment point, we then demonstrated that by controlling the co-expression of Tag/Catcher fused and unmodified scaffold building blocks in *E. coli* we can produce “hybrid” scaffolds with different building block ratios. A one-step purification approach allowed the isolation of these pre-configured hybrid scaffolds for subsequent attachment of cargo proteins for cross-linking into higher order assemblies and for functionalization. We observed that hybrid scaffolds assemble as arrays of nanotubes under a range of conditions. Importantly, double-tagged GFP could readily be conjugated to these preformed scaffolds and yielded macroscale materials of cross-linked and stacked microtubes that organize into clusters or radial scaffold particles.

**Fig. 1.**
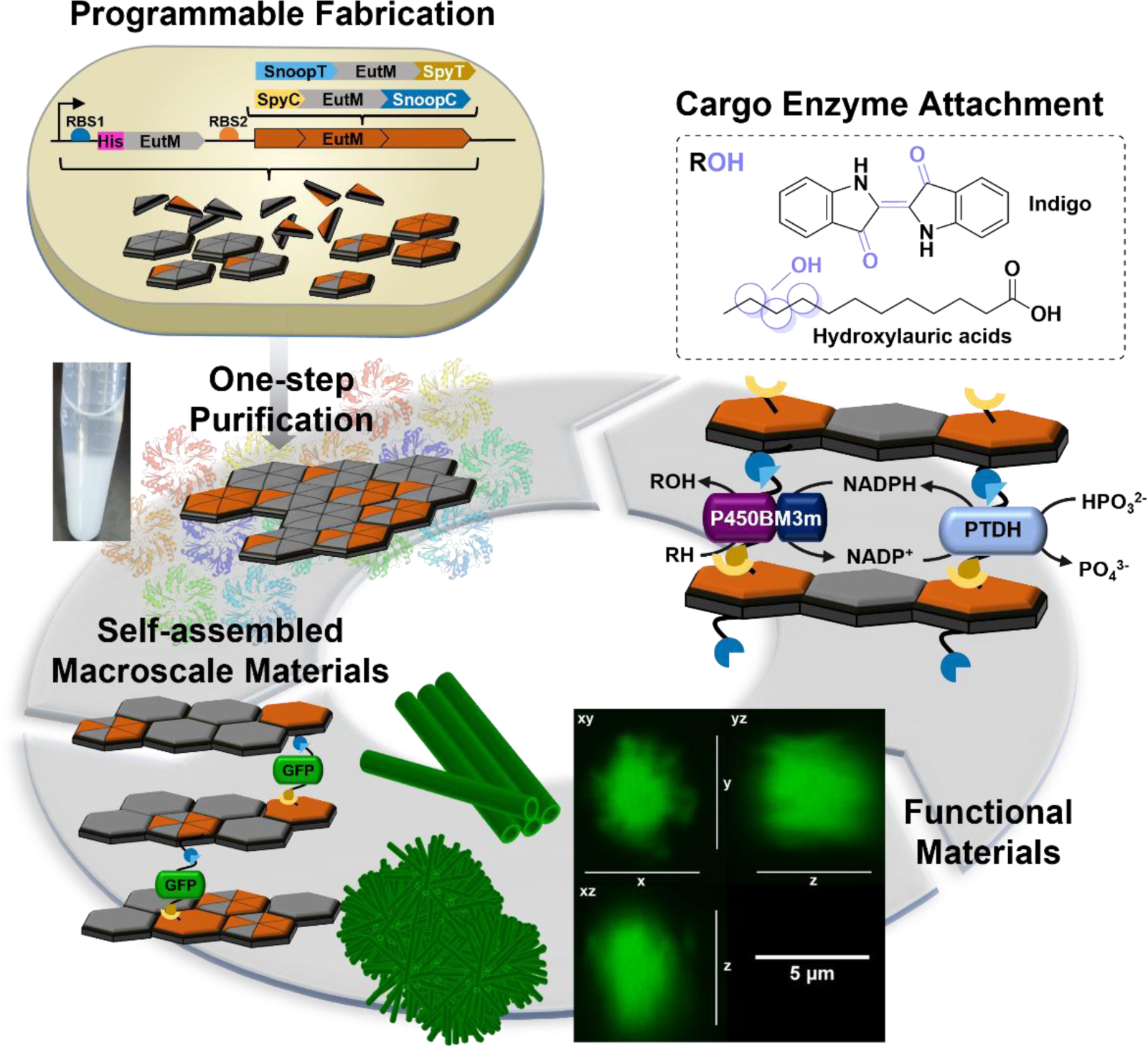
Fabrication of programmable self-assembling functional materials. *E. coli* is engineered for the programmable co-expression of unmodified and dual-modified EutM scaffold building blocks that self-assemble into hybrid scaffolds. A one-step purification yields macroscale scaffolds with controllable building block ratios. Scaffolds can then be cross-linked via isopeptide formation between Spy/SnoopTag and Catcher moieties displayed by the dual-modified EutM building blocks of the scaffolds and fused to cargo proteins of choice such as shown for GFP. Finally, the utility of the designed system for the assembly of functional materials is demonstrated with the co-immobilization of a dual-enzyme system composed of a complex, multi-domain P450 enzyme (P450BM3) and a phosphite dehydrogenase (PTDH) for co-factor recycling.

To show the applied utility of our system, we then confirmed that GFP can be replaced by cargo enzymes without compromising scaffold cross-linking and assembly as well as activity of the bioconjugated enzymes. We chose to attach a challenging yet for biocatalysis widely investigated, multi-domain cytochrome P450 enzyme (P450BM3) and validated its function in concert with a second, co-immobilized enzyme, phosphite dehydrogenase (PTDH) for co-factor recycling^17,18^. Despite decades of research dedicated to enzyme immobilization for biocatalytic process development, significant challenges remain in developing strategies that are cost-effective and especially, allow for the immobilization of multi-enzyme systems without loss of activities due to incompatible conjugation methods and supports^19^. Bioinspired protein-based bioconjugation and compartmentalization strategies are therefore increasingly explored to address these issues^2,3,8,12,16,20–26^. But in most studies, model enzyme systems or materials that lack robustness, modularity, and versatility as well as macroscale assembly properties are used for cell-free operation. We believe this work will add to the current landscape of genetically programmable materials by providing a platform for the design of robust and engineerable functional materials for a multitude of applications.

## RESULTS AND DISCUSSION

### Design and characterization of building blocks for scaffold assembly

Our first goal was to modify our EutM scaffolds to allow not only for the immobilization of biocatalysts as we have done before^12,14^, but also facilitate the formation of larger and more complex hierarchical structures without impeding scaffold assembly. We chose to use the efficient SpyCatcher/Tag system together with the similar SnoopCatcher/Tag system^15^ for isopeptide-mediated cross-linking of double-tagged EutM scaffolds and cargo proteins with cognate tags into higher-order assemblies (**Fig. 2a**). From previous work, we knew that the fusion of the larger SpyC domain is tolerated by EutM scaffolds, while fusion of this domain can negatively affect cargo enzyme activity^12,14^. We therefore designed two opposite configurations with either the smaller Tags (SpyT = 1.4 kDa, SnoopT = 1.5 kDa) or larger Catcher (SpyC = 9.1 kDa, SnoopC = 12.6 kDa) moieties fused to the N– and C-termini of EutM. As a model cargo protein for characterization of the effectiveness of scaffold building block cross-linking and for subsequent visualization, we engineered GFP with corresponding fusions. For purification of the recombinant proteins by one-step nickel affinity chromatography, an N-terminal His-tag was added to the two EutM scaffold building blocks (His-SnoopT-EutM-SpyT, His-SpyC-EutM-SnoopC) and GFP cargo proteins (His-SpyT-GFP-SnoopT, His-SnoopC-GFP-SpyC).

**Fig. 2.**
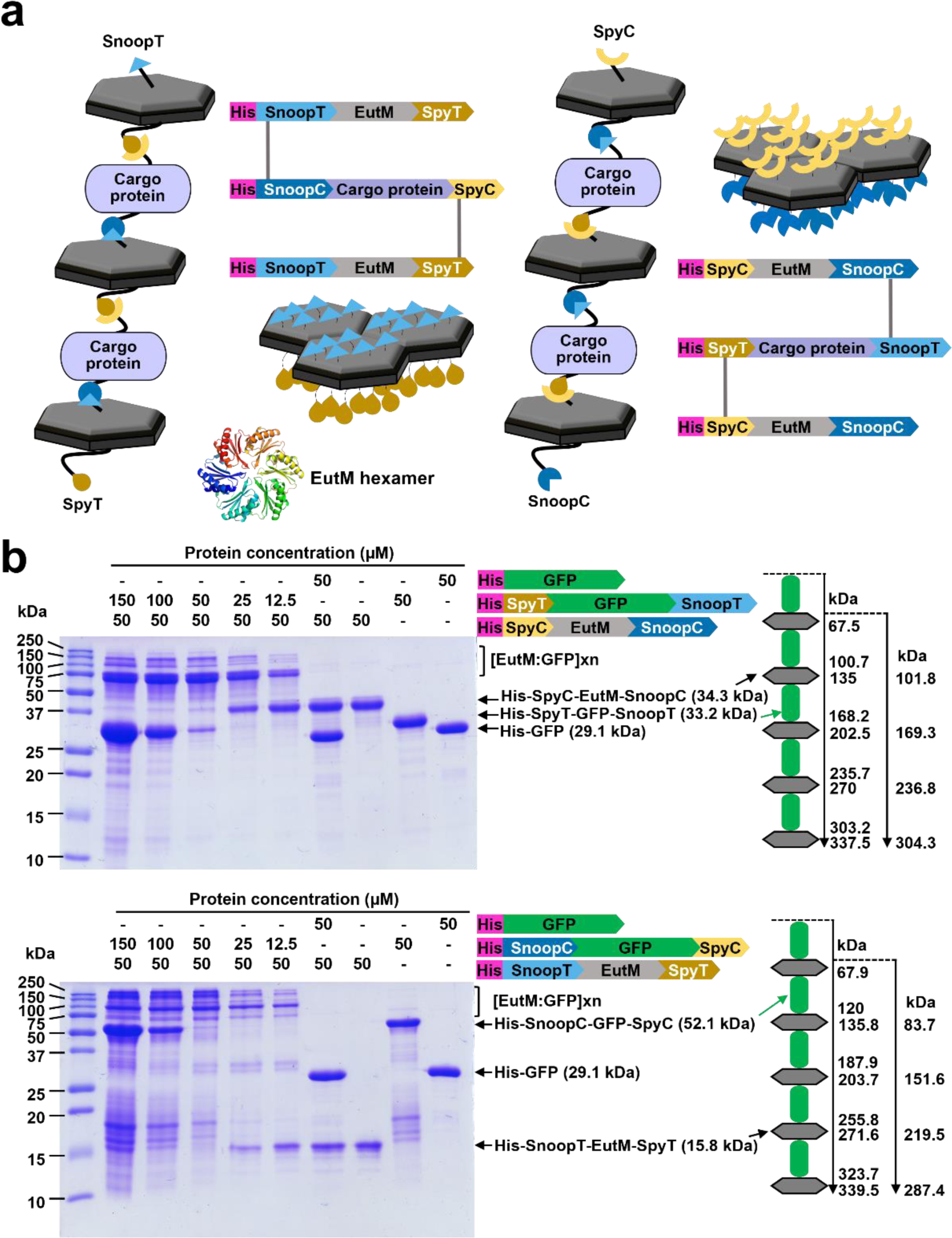
Design and testing of scaffold building block cross-linking. **a** EutM scaffold-building blocks self-assemble into hexamers that then assemble into larger scaffolds. Cross-linking of scaffold building blocks is achieved with N– and C-terminal fusion of SpyCatcher/Tag (SpyC, SpyT) and SnoopCatcher/Tag (SnoopC, SnoopT) moieties to EutM and cargo-proteins. Isopeptide bond formation (gray line) between cognate Spy/Snoop Catcher/Tag pairs allows for directional linkages shown for the two types of EutM building blocks designed in this work. The illustration shows one out of six conjugation sites per EutM hexamer. **b** Cross-linking of EutM building blocks with GFP was analyzed by SDS-PAGE. Purified His-SpyC-EutM-SnoopC or His-SnoopT-EutM-SpyT was mixed with GFP with or without (control) cognate Catcher/Tag fusions at the molar concentrations shown. Proteins were incubated in 0.1 M sodium phosphate buffer (pH 7.0) at 25 °C for 1 h prior to analysis. Expected molecular weights for one and multiple cross-linked GFP-EutM units are shown to the right. Higher molecular weight bands > 67 kDa corresponding to EutM-GFP conjugates form only with dual-modified GFP; with the majority of the GFP conjugated to EutM at a 1:1 molar ratio. Shown data are representative for one set of purified proteins.

Both dual-modified EutM building blocks could be readily overproduced and isolated from *E. coli*. We noticed though that the two purified proteins were much more soluble than our previously characterized His-EutM-SpyC and His-EutM^11–14^. The Catcher modified His-SpyC-EutM-SnoopC remained soluble at >20 mg/mL, while the Snoop/SpyT modified His-SnoopT-EutM-SpyT precipitated out of solution at ∼5 mg/mL. No large structures were observed for either protein by transmission electron microscopy (TEM). In contrast, as we have previously shown^11–14^, His-EutM assembles into hexameric rolled up or flat scaffolds that precipitate out at 2 mg/mL, while His-EutM-SpyC is more soluble and begins to scaffold out of solution at >5 mg/mL assemblies of long fibril-like structures as we observed previously^11,13^. Because of the formation of insoluble scaffolds, all EutM building block purifications were performed in the presence of 4 M urea for scaffold solubilization to maximize protein yield as described previously^11,13^. Removal of urea (and imidazole after nickel affinity purification) then causes the reformation of insoluble scaffolds (visible as a white pellet, see **Fig. 3a)** except for the highly soluble His-SpyC-EutM-SnoopC scaffold building block. Native PAGE analysis of scaffolds solubilized in urea (**Supplementary Fig. 1a**) and after removal of urea (**Supplementary Fig. 1b**) show similar higher molecular weight assemblies of ∼400-800 kDa in size for each dual-modified building block, suggesting that scaffolds are able to migrate into the gel form the same stable assemblies under both conditions. The assemblies appear to be composed of two (His-SpyC-EutM-SnoopC) or eight (His-SnoopT-EutM-SpyT) hexamers based on their molecular weights (**Supplementary Fig. 1c**). For comparison, His-EutM scaffolds seem to be assembled from six hexamers (**Supplementary Figs. 1a-c**). Surprisingly, we did not observe a band with a size corresponding to a single hexamer, which was the major band obtained for the bacterial microcompartment shell protein (RmmH) from *Mycobacterium smegmatis*^27^. From these results, it appears that the N– and C-terminal fusions added to EutM do not affect monomer assembly into hexamers but rather the assembly into larger scaffolds as previously observed for His-EutM building blocks without an N-terminal fusion^11^.

**Fig. 3.**
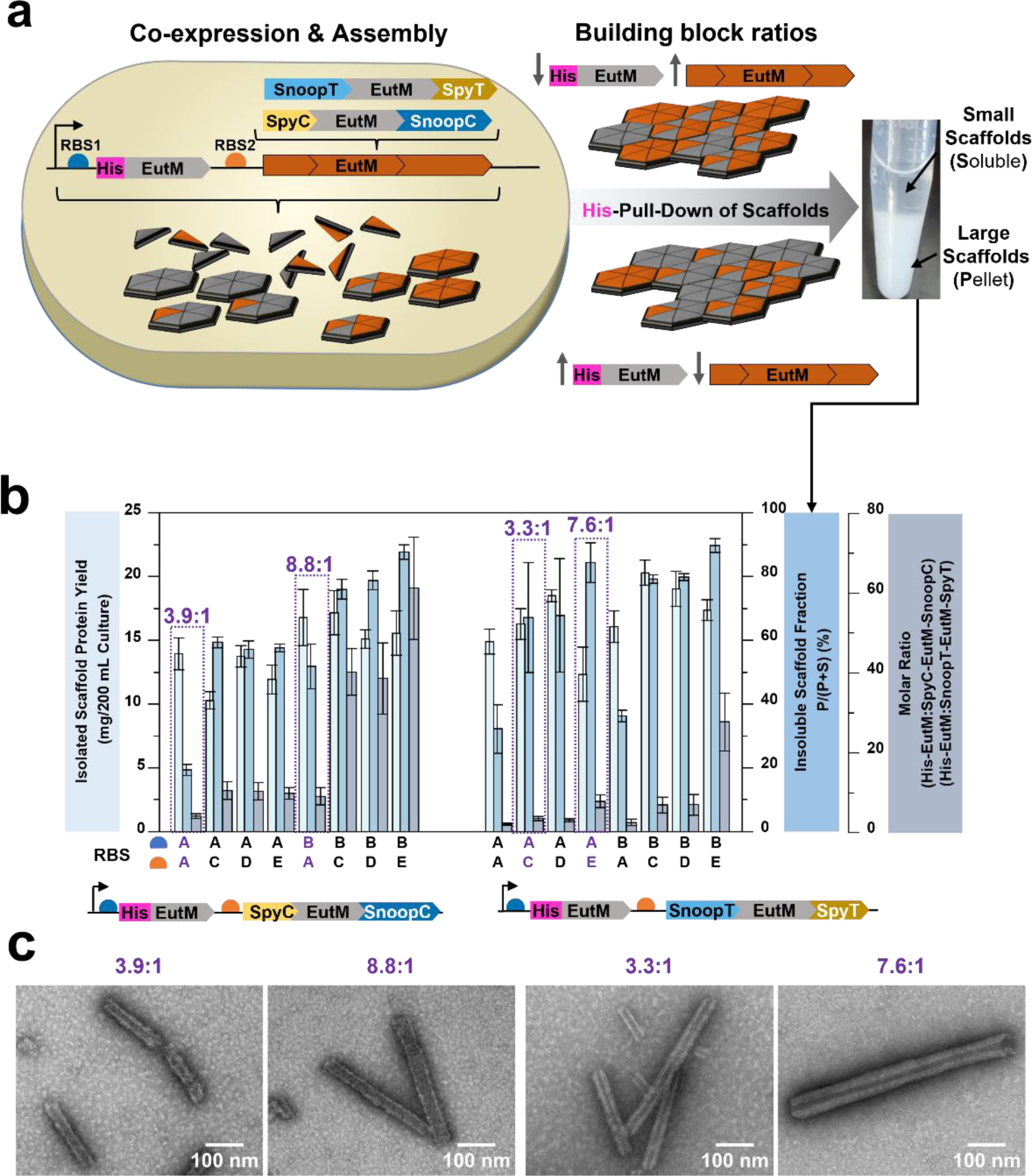
Production and characterization of macroscale hybrid scaffolds. **a** Co-expression of unmodified and dual-modified EutM scaffold building blocks in *E. coli* results in the self-assembly of hybrid scaffolds. By varying the expression levels of the building blocks with ribosome binding sites of different strengths, hybrid scaffolds with different molar ratios of building blocks are produced. Hybrid scaffolds can be selectively pulled-down by His-tag affinity purification, yielding purified scaffolds that separate into a soluble fraction (S) composed of smaller scaffolds and an insoluble fraction (pellet, P). Shown are co-expression results for constructs where the His-tag is on the unmodified EutM. **Supplementary Fig. 4** shows results for the same constructs where the dual-modified EutMs contain an N-terminal His-tag. **b** Characterization of hybrid scaffolds co-assembled from His-EutM and dual-modified EutM. Eight RBS combinations (RBS A-E) were used to vary scaffold building block ratios. Scaffolds were purified from 200 mL cultures in the presence of 4 M urea and the total yield of isolated scaffolds was quantified. The percentage of the insoluble scaffold fraction of normalized scaffolds (2 mg/mL) (as an indicator of assembly strength) was determined after dialysis into 0.1 M Tris-HCl buffer (pH 7.5) to remove urea. The molar ratios between EutM building blocks of the isolated hybrid scaffolds were analyzed by SDS-PAGE densitometry. Two hybrid scaffolds (highlighted by purple boxes and fonts) for each of the two building block combinations that have comparable molar ratios were selected for further studies. Data are shown as mean values ± SD and error bars represent the standard deviations of replicates from three independent cultures for each expression construct. **Supplementary Fig. 5** shows representative SDS-PAGE gels used for analysis. **c** Selected hybrid scaffolds were analyzed by TEM after dialysis into 0.1 M sodium phosphate buffer (pH 7.0) and dilution to 1 mg/mL. Shown are representative images of scaffolds captured at 60kx magnification (see **Supplementary Fig. 6** for additional images and magnifications (10k-120kx)).

With scaffold self-assembly of the dual-modified EutM building blocks confirmed, we next investigated covalent cross-linking of these building blocks with the corresponding dual-modified GFP fusion proteins as a critical step for the design of functionalized macroscale materials (**Fig. 2b, Supplementary Fig. 2**). Mixing of the modified EutM building blocks with their cognate GFP fusion protein partners at different molar ratios results in rapid cross-linking that is complete within 1 h based on SDS-PAGE analysis. As we have observed previously for EutM-SpyC, a small amount of Catcher domain modified EutM or GFP remained unconjugated regardless of molar ratios, incubation time or temperature (**Fig. 2b, Supplementary Fig. 2**), suggesting that their Catcher domains are not a competent state for peptide bond formation. Cross-linking progressed from one EutM monomer conjugated with one GFP to four or more EutM cross-linked by GFPs to create larger assemblies, including assemblies with molecular weights too large for SDS-PAGE separation. The formation of larger assemblies proceeded over time and seemed to be faster between His-SnoopT-EutM-SpyT and His-SnoopC-GFP-SpyC compared to His-SpyC-EutM-SnoopC and His-SpyT-GFP-SnoopT as observed by changes in SDS-PAGE gel banding patterns over time (**Supplementary Fig. 2**).

### Co-assembly of scaffold building blocks into macroscale hybrid scaffolds

Because the modification of both the N– and C-terminus of EutM impacted assembly into large, insoluble scaffolds desirable for macroscale material formation, we therefore sought to co-assemble hybrid scaffolds from dual-modified and unmodified EutM building blocks to promote the assembly of larger scaffolds. Co-assembly of these two building blocks would then also allow for the spatial distribution of bioconjugation points to control cross-linking and/or cargo attachment. In previous work we found that spacing of attachment points for enzyme immobilization on EutM-SpyC scaffolds was important for the optimization of biocatalyst activities. However, our first attempt of simply mixing in urea purified His-EutM and dual-modified His-EutM proteins followed by dialysis (for scaffold formation) did not yield the desired co-assembled hybrid scaffolds (**Supplementary Figs. 1 & 3**). Instead, each building block retained similar soluble assemblies as we observed for the individual blocks (**Supplementary Fig. 1**). In addition, the dual-modified building blocks remained largely in the soluble fraction after urea removal while the unmodified EutM scaffolded out of solution (**Supplementary Fig. 3**).

From these experiments, we concluded that the EutM monomers must self-assemble quickly into stable multimeric units (e.g. as hexamers and/or multiple hexamers) that cannot be disassembled and re-configured once expressed in *E. coli* and after solubilization in urea. To overcome this problem, we designed co-expression constructs for *E. coli* where individual building block expression levels are controlled by ribosome binding sites (RBS) with different strengths (**Fig. 3)**. We chose five different RBS (**Supplementary Table 1**) for co-expression of EutM and double-tagged EutM under the control of a cumate-inducible promoter^13,28–31^. Only one of the co-expressed building blocks contains an N-terminal His-tag (**Fig. 3b** shows the His-EutM designs and **Supplementary Fig. 4** the opposite designs), allowing for the pull-down of only hybrid-scaffolds by nickel affinity purification; an approach that we have previously used to confirm the co-assembly of EutM homologs from different bacteria^13^. Depending on building block expression levels, we expected to obtain scaffolds with different molar ratios of unmodified and dual-modified EutM building blocks. A higher molar ratio of unmodified EutM was anticipated to increase the fraction of larger scaffolds that pellet out of solution as illustrated in **Fig. 3a**.

We designed EutM co-expression constructs with different RBS combinations for each of the two dual EutM building blocks and placed the His-tag either on EutM or the modified EutMs, resulting in twenty-nine different designs (**Fig. 3b**, **Supplementary Fig. 4**). We then comprehensively characterized protein yield, scaffold assembly behavior (i.e. formation of large, insoluble scaffolds) and molar ratio of EutM to dual-modified EutM building blocks in the hybrid scaffolds (**Fig. 3b, Supplementary Fig. 4**). Scaffold production yields and molar ratios of scaffold building blocks were determined directly after nickel affinity purification pull-down in urea by measuring protein concentrations and by SDS-PAGE densitometry (**Supplementary Fig. 5**). Assembly behavior was compared by normalizing the concentration of all purified scaffolds in urea first to ∼2 mg/mL prior to dialysis into 0.1 M Tris-HCl pH 7.5 buffer for assembly into larger scaffolds. Insoluble scaffolds were then separated from soluble assemblies by centrifugation. Quantification of proteins in each fraction yielded the percentage of insoluble scaffold material formed as an indicator of assembly strength.

We found that scaffold yields, as well as the molar ratios of building blocks varied significantly across designs with no strong correlations between RBS combinations (**Fig. 3b**). Notably, scaffold yields were much lower for the designs where the His-tag was on the modified EutM building blocks because of its lower expression levels (**Supplementary Fig. 4**). Consequently, we chose to proceed only with the His-EutM containing hybrid scaffolds shown in **Fig. 3b** that are readily overproduced in *E. coli*. For these hybrid scaffolds, the ratios between EutM and dual-modified EutM ranged from 1.9:1 to 61:1, therefore providing options for creating scaffolds with different densities of bioconjugation sites. As expected, higher molar ratios of unmodified EutM increased the formation of larger, insoluble scaffolds for all hybrid scaffold co-expression designs. Likewise, hybrid scaffolds with SnoopT-EutM-SpyT showed stronger assembly into larger, insoluble scaffolds compared to those containing the Catcher modified EutM building block.

We selected two designs from each hybrid scaffold type with either SpyC/SnoopC or SnoopT/SpyT conjugation sites for additional characterization and cross-linking studies (**Fig. 3b**, purple boxes). The molar ratios of the four chosen hybrid scaffolds range from approximately 3:1 to 9:1, providing a density of conjugation sites comparable to ratios that we previously confirmed to work well for enzyme immobilization on EutM-SpyC scaffolds^12^. In addition, these four hybrid scaffold constructs (from here on referred to by their molar ratios: 3.9:1, 8.8:1, 3.3:1, 7.6:1) afford the isolation of insoluble hybrid scaffolds at good yields from recombinant *E. coli* cells. Importantly, TEM confirms that all four hybrid scaffolds self-assemble into crystalline structures composed of long, hollow rolled-up nanotubes with diameters ranging from 25-60 nm (**Fig. 3c, Supplementary Fig. 6**). Nanotube walls appear to be thicker and multi-layered for the SnoopT-EutM-SpyT containing hybrid scaffolds, with more pronounced layering, striations and larger nanotube diameters for the His-EutM:SnoopT-EutM-SpyT=7.6:1 hybrid scaffold which contains a higher molar ratio of unmodified EutM (**Supplementary Fig. 6**). The single-layered nanotubes look very similar to those reported for the bacterial microcompartment shell protein RmmH from *M. smegmatis*^27^, while the thicker tubes are more similar to the structures we have previously observed for EutM homologs^13^. Note that TEM analysis required dilution of scaffolds to 1 mg/mL to prevent grid breakage and allow imaging of individual structures.

### Characterization of hybrid scaffold assembly stability

Protein self-assembly, including the assembly of microcompartment shell proteins like EutM, is dynamic and dependent on environmental conditions such as protein concentration, pH, salts and temperature^32,3,10^. The usefulness of our designed protein materials for different applications such as biocatalysis will depend on their stable assembly across a range of commonly used conditions. Prior to proceeding with cargo attachment and cross-linking of our selected hybrid scaffolds, we therefore investigated the influence of buffers (pH 5.0-9.0), temperatures (4, 25, 30, 37 °C) (**Fig. 4**), and salt (100, 250 mM NaCl) (**Supplementary Figs. 7 & 8**) on their assembly stability by quantifying the percentage of insoluble scaffolds after 24 h incubation. For comparison, we also assessed the assembly stability of the individual scaffold building blocks under the same conditions.

**Fig. 4.**
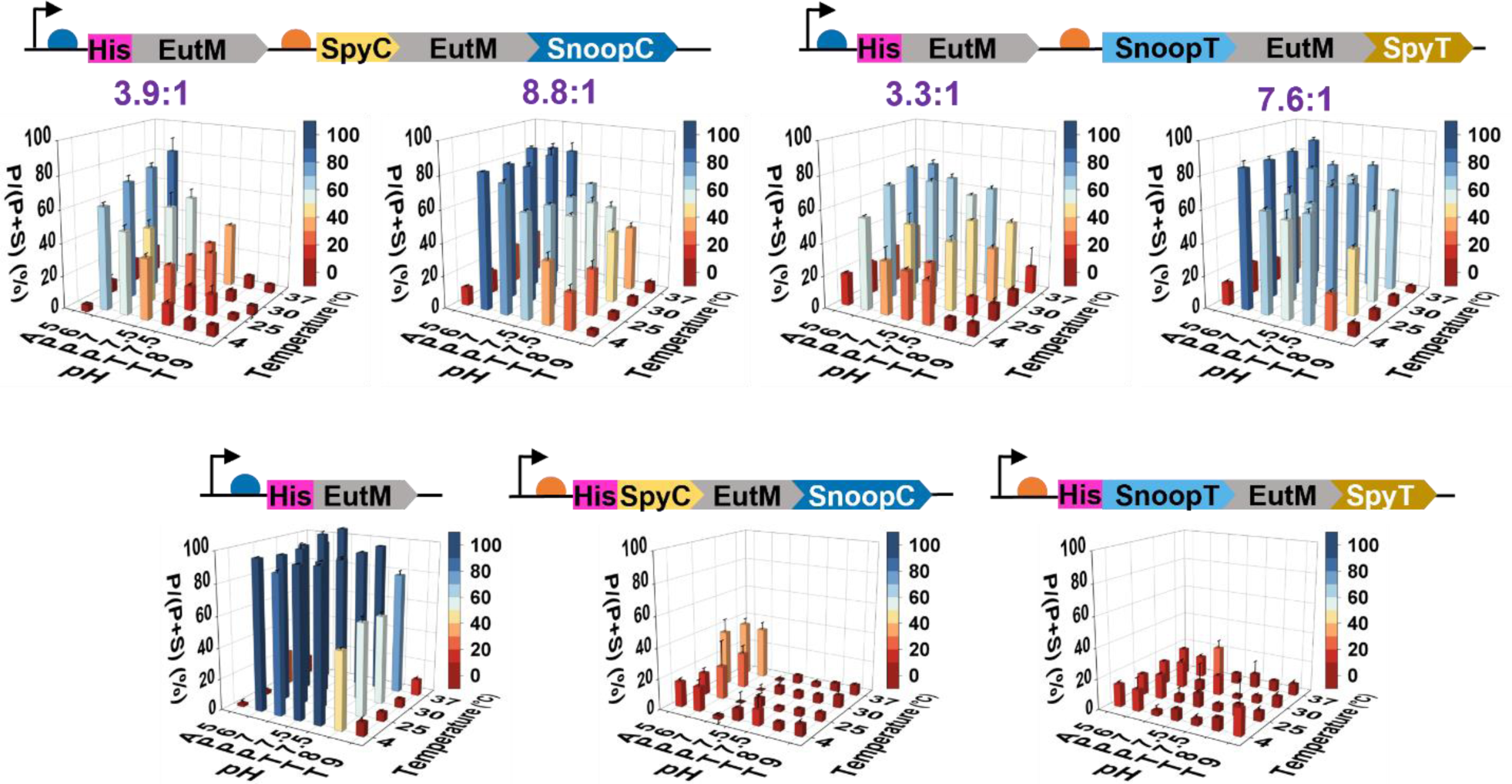
Characterization of scaffold assembly strength. Four hybrid scaffolds with different molar ratios (purple numbers) of co-expressed unmodified and dual-modified EutM building blocks were selected for characterization of assembly strength and compared to scaffolds formed by individual building blocks. Scaffolds (3 mg/mL in urea) were dialyzed into buffers with different pH values, normalized to 2 mg/mL and then incubated for 24 h at different temperatures. The percentage of the insoluble scaffold (P) fraction relative to the combined amount of soluble (S) and insoluble scaffold (P) ((P/(P + S) as in **Fig. 3**) was then calculated as indicator of assembly strength. The following buffers were used: 0.1 M sodium acetate (A, pH 5), 0.1 M sodium phosphate (P, pH 6, 7, or 7.5), and 0.1 M Tris-HCl (T, pH 7.5, 8 or 9). The same experiments were also conducted with 0.1 and 0.25 M NaCl added to buffers (**Supplementary Figs. 7 & 8**). Data are shown as mean values ± SD and error bars represent the standard deviations of three independent experiments per scaffold.

Consistent with the observations above, the dual-modified EutM building blocks showed the lowest assembly propensity, while the unmodified EutM protein formed stable, insoluble scaffolds that made up 49-100% of the total protein sample across all conditions except at pH 5.0 and 9.0. The assembly stabilities of the hybrid scaffolds fall in between their constituent building blocks and molar ratios (**Fig. 4, Supplementary Figs. 7 & 8**). Elevated temperatures increased self-assembly stability as we have observed previously for EutM homologs^12,13^. Assembly was optimal between pH 6.0-7.5, with a preference for a neutral pH 7.0 and phosphate instead of Tris-HCl buffers. Variations among scaffolds may be attributed to the different isoelectric points of the building blocks (calculated pI values are: His-EutM = 6.7, SpyC-EutM-SnoopC = 5.4, SnoopT-EutM-SpyT = 7.8). The addition of NaCl slightly decreased scaffold formation, except for the unmodified EutM (**Supplementary Figs. 7 & 8**). Together these results confirm that scaffold assembly is retained in conditions relevant for applications such as biocatalysis, and that higher EutM building block ratios increase self-assembly stability.

### GFP cargo attachment and cross-linking of hybrid scaffolds

Our next goal was to test whether our cross-linking model GFP cargo protein can still be efficiently conjugated to the dense, macroscale materials formed by the hybrid scaffolds. For these experiments, purified scaffolds (20-40 mg/mL) were first incubated with equimolar ratios of dual-modified GFP cargo (unmodified GFP as control) to dual-modified EutM building block (50 μM) in the hybrid scaffolds. After 1 h of incubation, large scaffold structures can be observed by light microscopy that are conjugated with the dual-modified GFP cargo (**Fig. 5**). Testing of different molar ratios of GFP cargo to dual-modified EutM (4:1 to 1:4, with the EutM partner fixed at 50 μM) demonstrated efficient cross-linking of EutM building blocks with 25-50 μM of GFP into higher molecular weight complexes (**Supplementary Fig. 9**). However, more unbound GFP remained after 1 h incubation with the hybrid scaffolds that have higher unmodified EutM ratios, suggesting that not all conjugation sites are readily accessible in these dense materials.

**Fig. 5.**
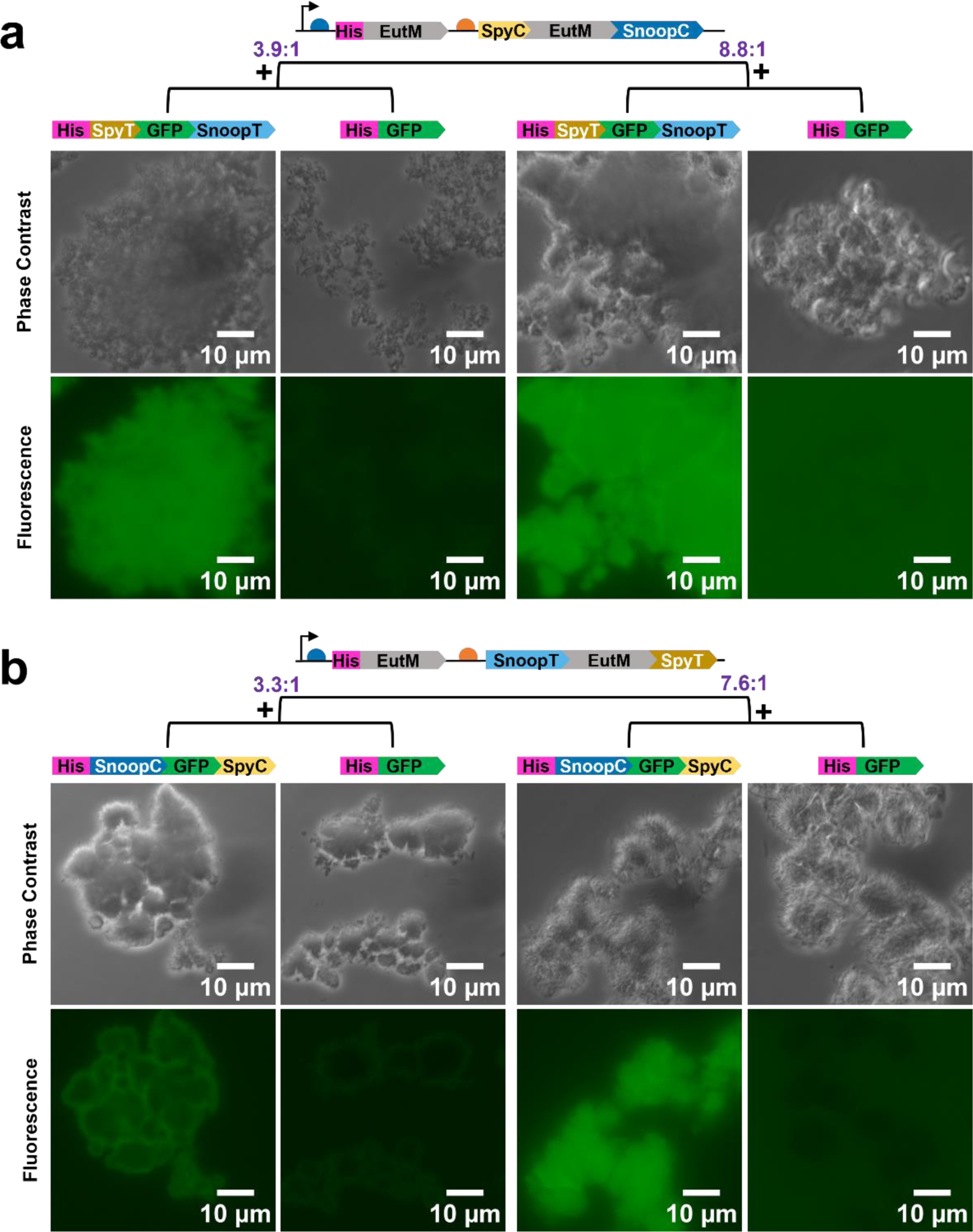
Imaging of hybrid scaffold structures and GFP attachment. **a** His-EutM:SpyC-EutM-SnoopC hybrid scaffolds (molar ratios of 3.9:1 and 8.8:1 from **Fig. 3**) were mixed with either His-SpyT-GFP-SnoopT or unmodified (control) His-GFP cargo proteins to observe scaffold structures and GFP cargo attachment. Scaffolds are visible as macroscale materials that appear more compact for His-EutM:SpyC-EutM-SnoopC=8.8:1 hybrid scaffolds that contain a higher ratio of unmodified EutM. GFP attachment to scaffolds is only observed with tagged GFP cargo. **b.** His-EutM:SnoopT-EutM-SpyT hybrid scaffolds (molar ratios of 3.3:1 and 7.6:1 from **Fig. 3**) were mixed with His-SnoopC-GFP-SpyC or unmodified (control) His-GFP cargo protein. Both hybrid scaffolds form macroscale materials visible as dense structures with radial microtube assemblies that are more pronounced for the His-EutM:SnoopT-EutM-SpyT=7.6:1 hybrid scaffolds. GFP cargo attachment is only observed with the Catcher modified GFP cargo. Experiments were performed by mixing 50 µM GFP cargo in 0.1 M sodium phosphate buffer (pH 7.0) with hybrid scaffolds such that concentration of their modified EutM building blocks are equimolar to that of their cognate GFP cargo partner (see Methods for protein concentrations). After incubation for 1 h at 25 °C and 180 rpm, samples were prepared for imaging. Representative images from one set of purified proteins are shown. Additional images, including for different GFP cargo ratios, are provided in **Supplementary Figs. 10 & 11**.

The morphologies of the hybrid scaffold materials vary, with more dense structures formed with higher ratios of unmodified EutM (His-EutM:SpyC-EutM-SnoopC=8.8:1, His-EutM:SnoopT-EutM-SpyT=7.6:1) (**Fig. 5**, see **Supplementary Figs. 10 & 11** for different GFP cargo ratios). Closer inspection of the His-EutM:SpyC-EutM-SnoopC=8.8:1 and both His-EutM:SnoopT-EutM-SpyT materials (visible at the fringes of the structures) suggest scaffolds are formed from stacked microtubes that assemble into crystalline appearing, radial particles that are particularly pronounced in the His-EutM:SnoopT-EutM-SpyT=7.6:1 scaffolds which are assembled from larger nanotubes (**Fig. 3c)**. To further characterize the structural features of GFP conjugated hybrid scaffolds, we chose cargo-loaded His-EutM:SpyC-EutM-SnoopC=8.8:1 hybrid scaffolds (**Fig. 5a**) that we would later use for enzyme immobilization for TEM and confocal microscopy imaging (**Fig. 6**, **Supplementary Fig. 12**). TEM images suggest that scaffold fibers are coated by GFP compared to the thinner and more articulated fibers observed in **Fig. 3c**. In addition, bundles of fibers are linked together, presumably by the conjuaged GFP cargo protein (**Fig. 6a**). Confocal microscopy, including 3D reconstructions, delineate the micrometer dimensions of radial particles that are conjugated throughout with GFP and their clustering into larger scaffolds (**Fig. 6b**), suggesting a scaffold assembling and cross-linking process that involves the hierarchical assembly of microtubes into cross-linked radial particle clusters illustrated in **Fig. 6c**.

**Fig. 6.**
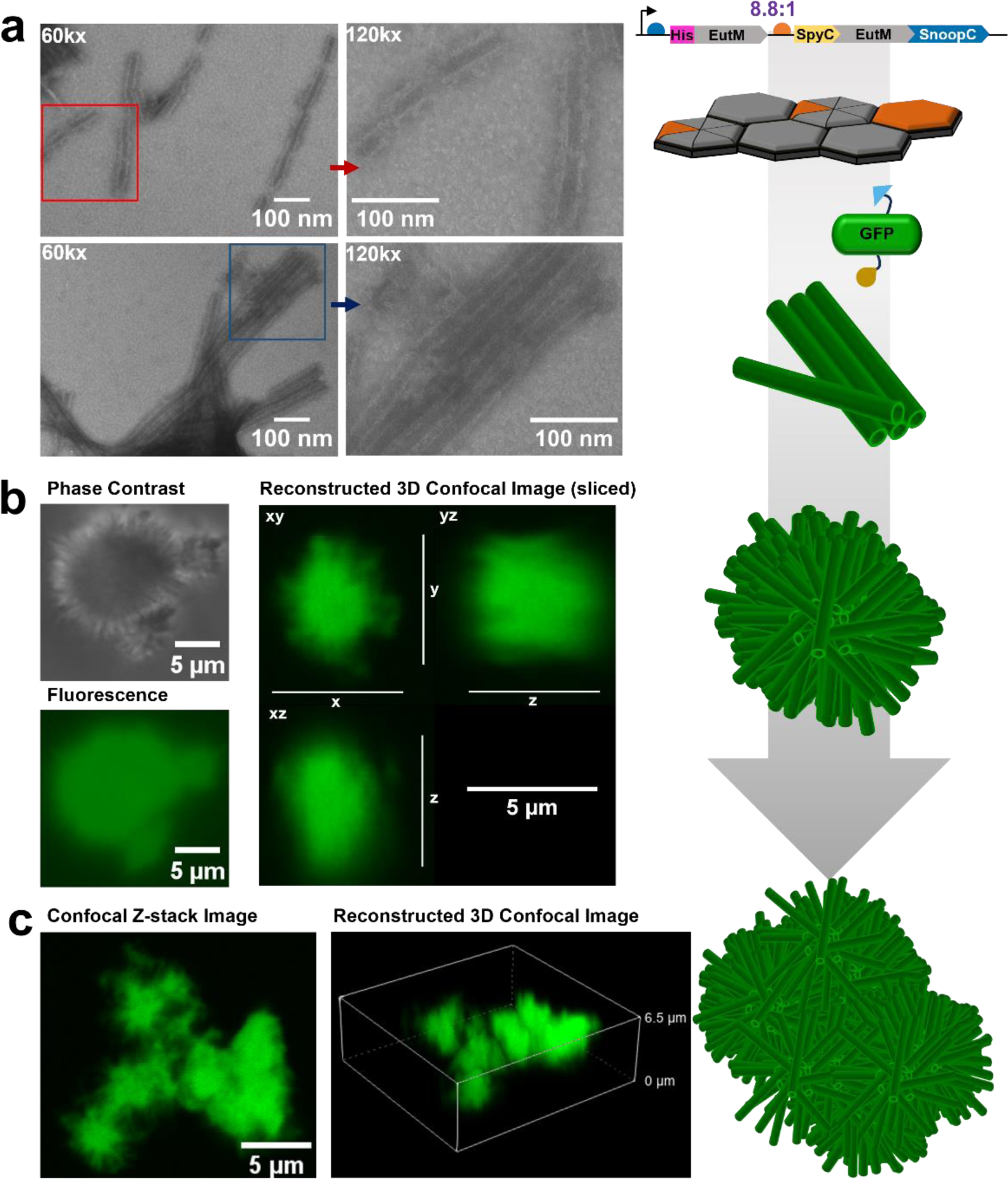
High-resolution microscopy imaging of GFP cross-linked hybrid scaffold assemblies. **a** TEM of hybrid His-EutM:SpyC-EutM-SnoopC=8.8:1 scaffolds with cross-linked His-SpyT-GFP-SnoopT cargo (see **Fig. 5a**) shows EutM scaffold nanofibers that appear to be coated (red arrows) with GFP compared to the thinner and more articulated tube-like scaffold structures observed in **Fig. 3c**. Bundles of fibers are observed that are enveloped by a film (blue arrows), suggesting cross-linking by the dual-modified GFP. Parallel and radially aligned fiber bundles can be observed that are reminiscent of the radial structures observed by light microscopy (see **Supplementary Fig. 12** for additional TEM images and magnifications). Red and blue boxes mark areas further magnified to the right. Samples are diluted to 1 mg/mL for TEM analysis. Representative images from one set of proteins are shown. **b** Phase contrast and confocal fluorescence microscopy of a single, radial scaffold particle and a larger scaffold assembly composed of clusters of radial particles from the same GFP-cross-linked hybrid scaffolds imaged by TEM. Images show the micrometer dimensions of the scaffolds. Confocal Z-stack imaging shows GFP fluorescence across the entire scaffold structures. **c** Imaging results suggest a scaffold assembling process that starts with the assembly of EutM hexamers into microtubes which form radial, microscale particles that are coated and cross-linked with GFP. Particles and bundles of microtubes cluster together into larger microscale scaffolds observed by microscopy and eventually, form the large insoluble macroscale materials that fall out of solution.

### Testing hybrid-scaffolds for enzyme immobilization

Our results showed that the designed hybrid scaffolds can be readily overexpressed and isolated from *E. coli* with configurable conjugation site densities. Further, the scaffolds are also stable and self-assemble into macroscale materials that are efficiently conjugated and cross-linked with GFP as model cargo protein. These are characteristics that are useful for the creation of customizable, functional materials for a range of biotechnological applications. As a first step towards the design of such materials, we sought to assess the utility of our scaffolding system for enzyme immobilization by replacing our model cargo protein GFP with two enzymes for the operation of a two-enzyme reaction.

As a proof-of-concept, we selected as our first enzyme for scaffold attachment the challenging, yet extensively studied and highly versatile multi-domain cytochrome P450 monooxygenase CYP102A1 known as P450BM3^18,33^. Unlike many P450 enzymes, P450BM3 can be recombinantly expressed as soluble protein, is self-sufficient due to a fused NADPH reductase domain, is known to catalyzes diverse reactions and over thousand variants have been described^18,33^. Yet, industrial use of this enzyme remains challenging due to its complex, two-domain and dimeric protein structure, undesired “uncoupling” of NADPH derived electrons to produce reactive and inactivating oxygen species and/or H_2_O_2_. To improve P450BM3 reaction efficiency and for catalysts and NADPH co-factor recycling, diverse immobilization methods have been developed, including the co-immobilization of P450BM3 with co-factor recycling enzymes^25,34–39^. From this knowledgebase, we chose for scaffold conjugation the commonly used P450BM3 variant GVQ (A74G, F87V, L188Q) (referred herein as P450BM3m). This enzyme variant oxidizes a range of substrates, including indole into the blue indigo for convenient activity assessment^40–42^. For co-factor recycling and the operation of a two-enzyme cascade (**Fig. 7a**), we selected as our second enzyme for scaffold attachment an engineered phosphite dehydrogenase (PTDH) that is known to work with P450BM3^17,37,43,44^.

**Fig. 7.**
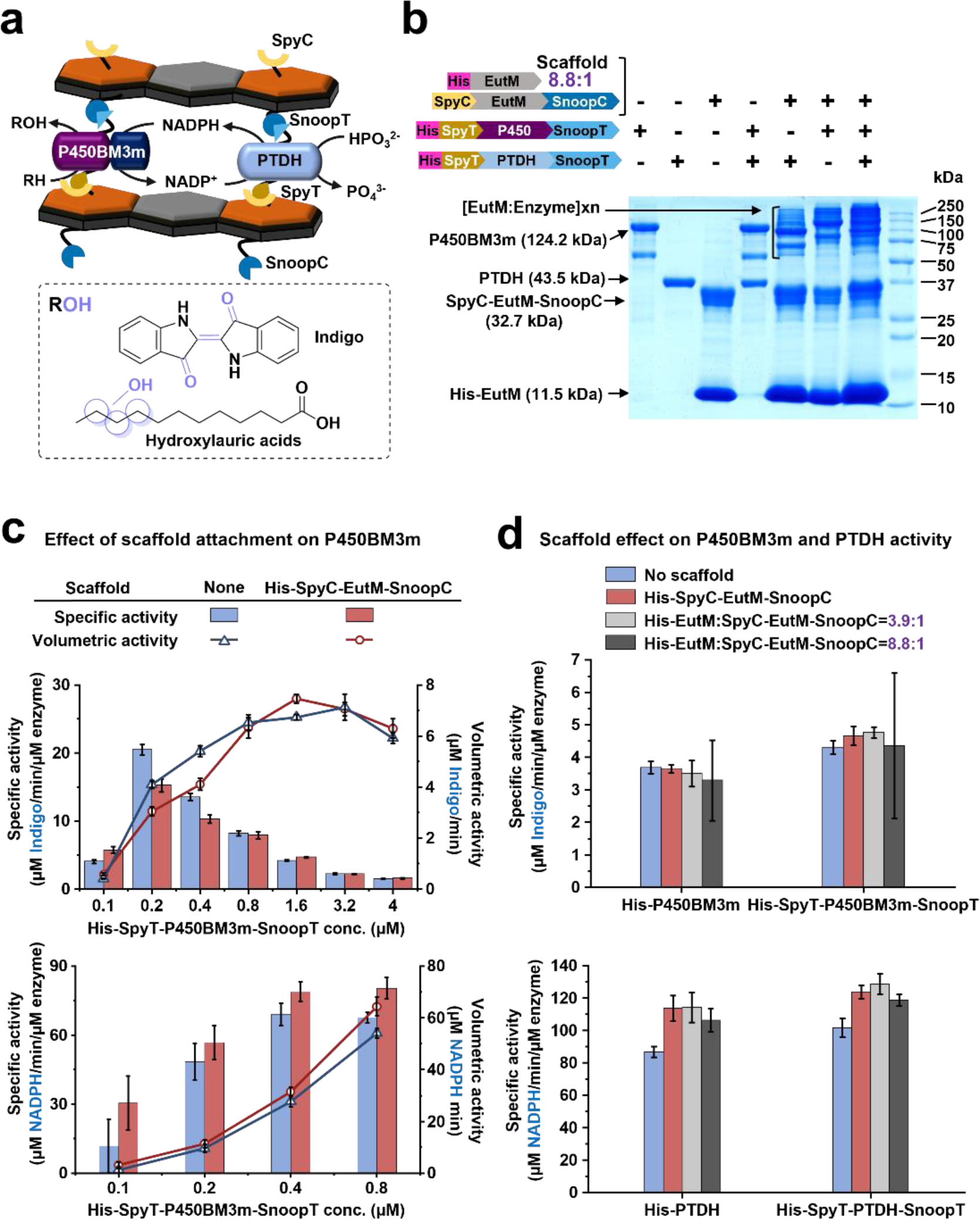
Immobilization of P450BM3m and PTDH onto hybrid EutM scaffolds. **a** P450BM3m and PTDH are co-immobilized onto His-EutM:SpyC-EutM-SnoopC=8.8:1 hybrid scaffolds via their N– and C-terminal cross-linking SnoopT and SpyT fusions. The resulting immobilized dual-enzyme system facilitates phosphite-driven NADPH co-factor recycling by PTDH for the P450BM3m catalyzed hydroxylation of indole into indoxyl (which dimerizes into indigo) or of lauric acid into 9–, 10– and 11-hydroxylauric acids. **b** Confirmation of cross-linking of SpyC-EutM-SnoopC in the His-EutM:SpyC-EutM-SnoopC=8.8:1 scaffolds with P450BM3m and PTDH. Enzymes were mixed with the hybrid scaffold at a 1:4 molar ratio corresponding to the SpyC-EutM-SnoopC building block in the scaffold. Samples in 0.1 M phosphate buffer (pH 7.0) were incubated for 1 h at 30 °C and 180 rpm prior to SDS-PAGE analysis. Control reactions were performed with and without enzymes and scaffolds as shown. The formation of high molecular weight complexes (as for GFP in **Fig. 1b**) confirms that the cross-linking fusion tags are functional. Results for His-EutM:His-SpyC-EutM-SnoopC=3.9:1 and His-SpyC-EutM-SnoopC scaffolds are found in **Supplementary Fig. 14**. **c** Effects of N– and C-terminal cross-linking of His-SpyT-P450BM3m-SnoopT to scaffolds on enzyme activity were determined by mixing the enzyme with His-SpyC-EutM-SnoopC at a 1:4 molar ratio in 0.1 M phosphate buffer (pH 7.0). After dilution to achieve different molar enzyme concentrations (0.5-20 µM), samples were incubated for 1 h at 30 °C to allow for isopeptide bond formation. Specific and volumetric activities of the immobilized enzyme samples with indole (2.5 mM) and NADPH (0.25 mM) at 30 °C and pH 7.0 were then measured with 40 µL of the immobilization mixtures in 200 µL reactions (5-fold dilution) to obtain the final enzyme concentrations shown. The formation of indigo and consumption of NADPH were spectrophotometrically monitored. Control reactions were performed without scaffolds. At P450 enzyme concentrations >0.8 µM, NADPH was consumed after 30 s, preventing reliable activity measurements. The same experiments were performed with unmodified His-P450BM3m as a control (see **Supplementary Fig. 15**). **d** Influence of different scaffolds on P450BM3m and PTDH activities was determined by immobilizing P450BM3m (8 µM) or PTDH (0.5 µM) as in **b** prior to activity measurements. Control reactions were performed without scaffolds and enzymes without SpyT and SnoopT fusions. Specific activities for P450BM3m were measured with indole as in (**c**) by monitoring indole formation. Specific activities for PTDH were measured with sodium phosphite (1 mM) and NADP^+^ (0.25 mM) by monitoring NADPH formation. Reactions were performed with 40 µL immobilization samples in 200 µL assays at 30 °C and pH 7.0. Using the same set-up, stability of the immobilized enzymes was determined by measuring specific activities immediately and after up to 168 h (30 °C, pH 7.0) after mixing of enzymes and scaffolds (**Supplementary Figs. 16 & 17**). For (**b-c**), detailed protocols with protein concentrations are provided in the Methods. For (**c**) and (**d**), data are shown as mean values ± SD and error bars represent the standard deviations of four replicates with one set of purified proteins.

For scaffold conjugation and cross-linking, we modified the N– and C-termini of P450BM3m and PTDH with Spy/Snoop-Catcher and –Tag fusions as we did for GFP. While comparing the kinetic properties of the dual-modified and unmodified enzymes with indole or lauric acid and NADPH as substrates (**Table 1 & 2**) we found that fusion of the larger Catcher domains (9.1 kDa SpyC, 12.6 kDa SnoopC) completely inactivated P450BM3m and reduced the activity of PTDH more than 10-fold. The addition of the smaller Spy/Snoop-Tag did not significantly impact enzyme activities and thus yielded active enzymes for conjugation to SpyC-EutM-SnoopC modified scaffolds. We noted that under the reaction conditions tested (0.1 M sodium phosphate buffer, pH 7.0, 30°C) P450BM3m exhibited non-Michaelis-Menten, sigmoidal kinetics (**Supplementary Fig. 13**), which has been reported for P450BM3 with non-natural substrates like indole and different buffer conditions^42,45–49^. This sigmoidal behavior was more pronounced when measuring steady-state kinetics with varying concentrations of NADPH (5 mM indole) as indicated by a more than 3-fold higher Hill coefficient (2.0 vs. 6.6 for His-SpyT-P450BM3m-SnoopT) (**Table 1**), suggesting homotropic cooperativity and/or NADPH uncoupling.

**Table 1:** P450BM3m kinetic parameters. Enzyme reactions were performed at 30 °C in sodium phosphate buffer (pH 7.0) with 0.4 µM His-tagged wild-type dual-modified P450BM3m (see Methods). The His-SnoopC-P450BM3m-SpyC fusion protein was inactive, and no kinetic parameters could be measured. Data are shown as mean values ± SD and error bars represent the standard deviations of four replicates with one set of purified proteins. Kinetic fitting curves are shown in **Supplementary Fig. 13**.

| Enzyme | $K_m$ (μM) | $K_{cat}$ ($s^{-1}$) | $K_{cat}/K_m$ ($M^{-1} s^{-1}$) | Hill Coefficient |
| --- | --- | --- | --- | --- |
| **Indole<sup>1</sup>** |  |  |  |  |
| His-P450BM3m | 631.8 ± 46.8 | 0.1 ± 0.01 | 2.3 $\times 10^2$ | 2.7 ± 0.3 |
| His-SpyT-P450BM3m-SnoopT | 691.3 ± 18.3 | 0.2 ± 0.003 | 2.8 $\times 10^2$ | 2.0 ± 0.1 |
| **Indole<sup>2</sup>** |  |  |  |  |
| His-P450BM3m | 179.3 ± 22.2 | 0.6 ± 0.03 | 3.3 $\times 10^3$ | 1.6 ± 0.2 |
| His-SpyT-P450BM3m-SnoopT | 302.1 ± 27.8 | 0.8 ± 0.03 | 2.6 $\times 10^3$ | 1.3 ± 0.1 |
| **NADPH<sup>1</sup>** |  |  |  |  |
| His-P450BM3m | 70.7 ± 2.3 | 0.1 ± 0.01 | 2.1 $\times 10^3$ | 7.0 ± 1.2 |
| His-SpyT-P450BM3m-SnoopT | 74.4 ± 0.8 | 0.2 ± 0.004 | 2.4 $\times 10^3$ | 6.6 ± 0.2 |
| **Lauric acid<sup>2</sup>** |  |  |  |  |
| His-P450BM3m | 121.1 ± 14 | 2.2 ± 0.1 | 1.8 $\times 10^4$ | 1.5 ± 0.3 |
| His-SpyT-P450BM3m-SnoopT | 95.8 ± 6.2 | 2.5 ± 0.1 | 2.6 $\times 10^4$ | 1.8 ± 0.3 |

**Table 2:**
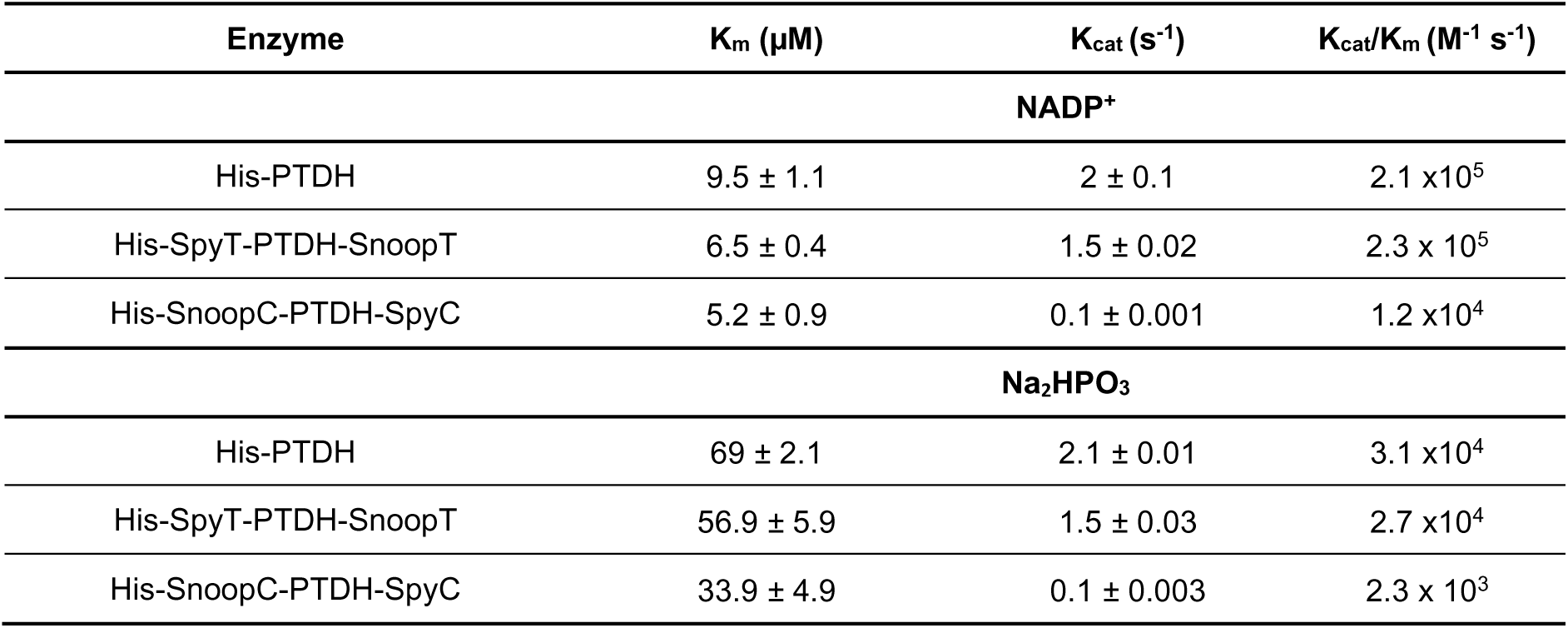
PTDH kinetic parameters. Enzyme reactions were performed at 30 °C in sodium phosphate buffer (pH 7.0) with 0.03-0.25 µM PTDH (see Methods). Data are shown as mean values ± SD and error bars represent the standard deviations of four replicates with one set of purified proteins.

Each dual-modified or unmodified (control) enzyme (His-SpyT-P450BM3m-SnoopT, His-SpyT-PTDH-SnoopT) was then individually conjugated His-SpyC-EutM-SnoopC and hybrid His-EutM:SpyC-EutM-SnoopC scaffolds at a 1:4 molar ratio to the dual-modified EutM building block. At this molar ratio, we obtained efficient conjugation of GFP cargo and are expecting to achieve sufficient spacing of the immobilized catalysts^12^, especially considering the large size of P450BM3m (248 kDa for the dimer). Both enzymes were efficiently cross-linked individually as well as together to scaffolds and formed higher molecular weight complexes as observed with the GFP cargo (**Fig. 7b**, **Supplementary Fig. 14**). Because P450BM3m is a highly dynamic protein that associates as a functional dimer^50^, we first determined its specific and volumetric activities at different enzyme concentrations for indigo formation and NADPH consumption when conjugated at either terminus to His-SpyC-EutM-SnoopC scaffolds (1:4 molar ratio enzyme to scaffold building block) (**Fig. 7c**). The highest volumetric activity for indigo formation was achieved with 1.6 µM of dual-modified P450BM3m attached to scaffolds. Scaffold attachment also benefited NADPH oxidation activity. Interestingly, control reactions with unmodified P450BM3m and scaffold protein show a similar trend, suggesting that the scaffolds provide a beneficial reaction environment for the enzyme (**Supplementary Fig. 15)**.

Next, we compared specific activities of P450BM3m and PTDH immobilized to different scaffolds (1:4 molar ratio of enzyme to conjugating scaffold building block). Control reactions were performed with no scaffolds and with unmodified enzymes. The final reactions were performed with the optimal P450BM3m concentration of 1.6 µM (which gave the highest volumetric activity, **Fig. 7c**) or with 0.1 µM of the much more active PTDH (**Tables 1 & 2**). As observed before, scaffolds did not reduce enzyme activities and cross-linking slightly increased specific activities of P450BM3m or PTDH (**Fig. 7d, Supplementary Figs. 16 & 17**). Scaffold type did not significantly affect the enzymes’ specific activities, except that activites with His-EutM:SpyC-EutM-SnoopC=8.8:1 hybrid scaffolds were slightly lower. Due to the high turbidity of these macroscale scaffolds that also adsorbed some of the formed indigo (yielding blue scaffold material), P450 activity measurements were particularly challenging, resulting in large standard deviations. Finally, we measured the stability of the enzymes, by incubating the scaffolded and unscaffolded enzymes and the controls under the above conditions (pH 7.0, 30°C) and concentrations (8 µM P450BM3m or 0.5 µM PTDH conjugated at a 1:4 molar ratio of enzyme to conjugating scaffold building block) for up to 168 h (7 days) prior to actvity measurements with five-fold diluted samples containing 1.6 µM P450BM3m or 0.1 µM PTDH (**Supplementary Figs. 16 & 17**). Under these incubation conditions, P450BM3m was remakable stable with little loss of activity with or without scaffolds present. PTDH was less stable with the unmodified and dual-modified enzymes retaining 50% and 35%, respectively, of their activities after 7 days. In the presence of scaffolds, the stabilities of the enzymes increased about 15%. Interestingly, only the hybrid scaffolds increased dual-modified PTDH stability, while scaffold protein addition regardless of type stabilized the unmodified PTDH, suggesting that surface attachment and the resulting different electrostatic environments matter for enzyme stability.

### Co-Immobilization of P450BM3m and PTDH operation of two-enzyme reaction

Cross-linking studies with the individual, dual-modified enzymes demonstrated that the designed scaffolding system functions as an effective enzyme immobilization platform that is even compatible with a large, multi-domain enzyme like P450BM3m. In addition, the scaffolds can stabilize less stable enzymes as we have observed before^12,14^. As a final test, we then co-immbolized P450BM3m and PTDH onto our scaffolds for co-factor recycling and to demonstrate catalysis with more than one enzyme (**Fig. 7a**). Dual-modified P450BM3m and PTDH are efficiently co-immobilized to hybrid His-EutM:SpyC-EutM-SnoopC=8.8:1 scaffolds, forming large molecular weight complexes similar to those observed with the conjugated, individual enzymes (**Fig. 7b**). Negative staining TEM of diluted hybrid scaffolds (**Fig. 8a**, **Supplementary Fig. 18**) with the co-immobilzed enymes show parallel aligned tubes that are enveloped by a thick film, indicating successful coating and cross-linking of scaffolds with the two cargo enzymes as observed with GFP (**Fig. 6a, Supplementary Fig. 12)**.

**Fig. 8.**
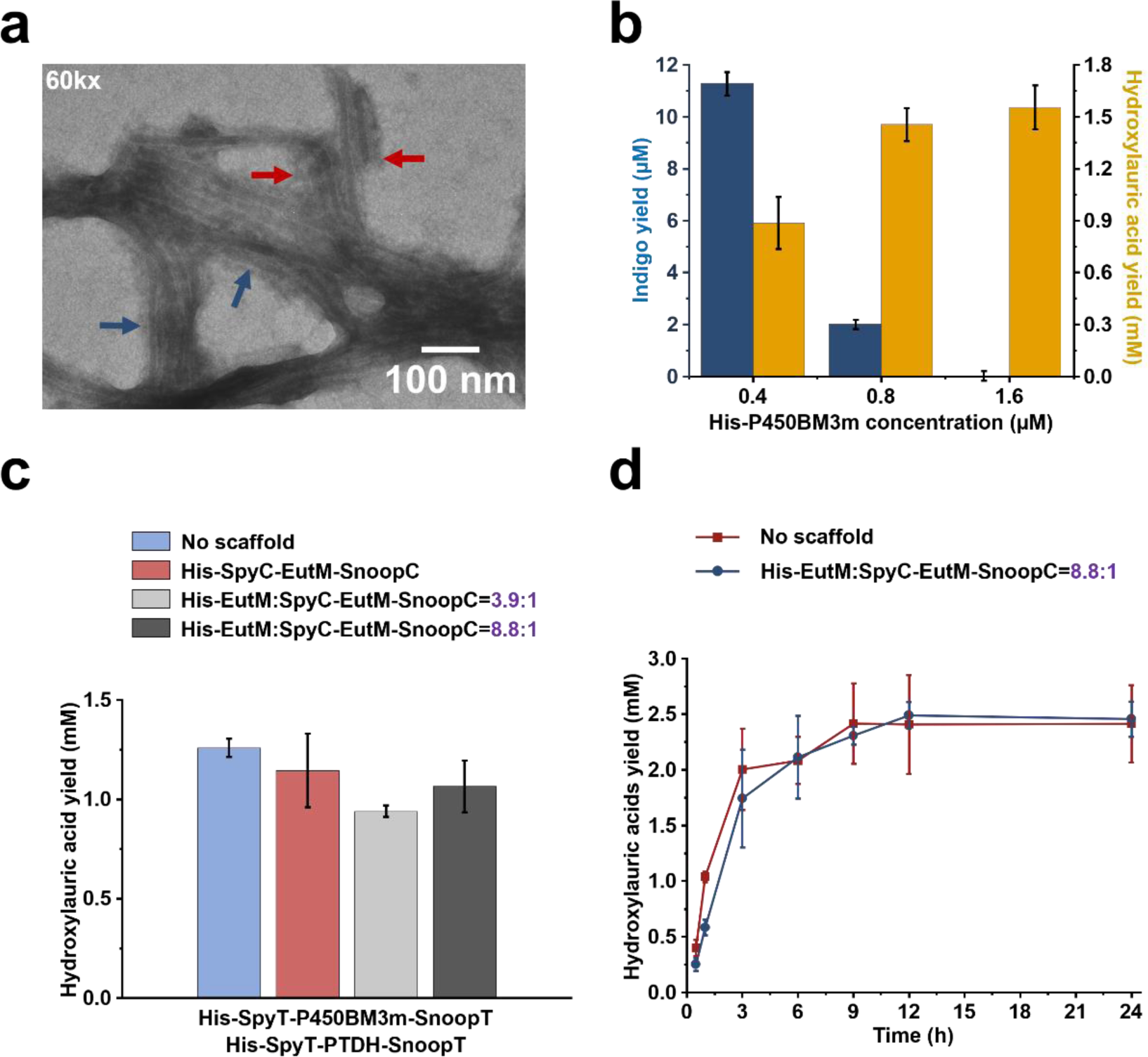
Co-immobilization of P450BM3m and PTDH onto hybrid scaffolds. **a**. TEM of His-EutM:SpyC-EutM-SnoopC=8.8:1 hybrid scaffolds with co-immobilized P450BM3m and PTDH shows cross-linked tube-like structures enveloped in a dense coating as in **Fig. 6a** with GFP. Samples were prepared by co-immobilizing 15 µM of each enzyme with 120 µM His-EutM:SpyC-EutM-SnoopC=8.8:1 as in **Fig. 7b**. Samples were diluted (1 mg/mL) prior to TEM. Additional images and magnifications (10k-120kx) are provided in **Supplementary Fig. 18**. **b.** Testing of coupled reaction system with free His-PTDH and His-P450BM3m. For indole conversion (blue), reactions were performed at 30 °C and pH 7.0 in 0.1M phosphate buffer with1.6 µM His-PTDH and 0.4, 0.8, or 1.6 µM His-P450BM3m and 2.5 mM indole, 10 mM sodium phosphite and 0.25 mM NADP^+^. After 15 min, indigo was spectrophotometrically quantified. For lauric acid conversion (gold), reactions were performed under the same conditions and concentrations but with 2.5 mM lauric acid. Reactions were stopped after 15 min and hydroxylauric acids (combined 9-, 10-, 11-hydroxylauric acids, see **Supplementary Figs. 22 & 23**) were quantified by GC-FID. **c** Demonstration of coupled reaction with P450BM3m and PTDH co-immobilized onto scaffolds. Dual-modified P450BM3m and PTDH (6.4 µM each) were co-immobilized onto scaffolds (or no scaffolds as a control) at a 1:4 molar ratio of enzymes to cross-linking scaffold building block in the scaffolds as in **Fig. 7d**. 1 mL lauric acid conversion reactions were then performed for 10 min with 500 µL of enzyme mixture (3.2 µM final enzyme concentrations) with the conditions in **b**. Relative amounts of hydroxy lauric acid conversion products for the reactions are shown in **Supplementary Table 2**. **d** Two-phase reaction system for lauric acid conversion. A 5 mL reaction was set up with 2.5 mL immobilized enzyme mixtures as in **c**. but with 0.8 µM His-SpyT-P450BM3m-SnoopT, 3.2 µM His-SpyT-PTDH-SnoopT, 50 mM sodium phosphite, 0.5 mM NADP^+^ and 20 mM lauric acid in 20% (v/v) dodecane. Reactions were performed at 30°C,120 rpm and hydroxylauric acid products quantified. Detailed protocols with protein concentrations for scaffolds and enzymes are provided in the Methods. Data in (**b**), (**c**) and (**d**) are shown as mean values ± SD and error bars represent the standard deviations of four replicates with one set of purified proteins.

During optimization of the coupled reaction of P450BM3m and PTDH with the free enzymes for NADPH recycling with indole as substrate, we unexpectedly found that at higher P450BM3m concentrations (including the optimal scaffold immobilization concentration of 1.6 µM determined in **Fig. 7c** for the highest volumetric activity) and 0.25 mM NADPH, indigo production was either completely abolished or greatly inhibited in the coupled system compared to P450BM3m only reactions with 0.25 mM NADPH (**Fig. 8b, Supplementary Fig. 19)**. We suspected that this inhibition may be caused by the production of inactivating oxygen species and/or H_2_O ^34,51,52^ due to uncoupling of NADPH electron transfer with the non-native indole as substrate due to the maintenance of a high NADPH concentration by the co-factor recycling PTDH. The inhibition of indole conversion can be replicated with both the unmodified and dual-modified His-P450BM3m enzyme in reactions with both high NADPH (3.5 mM) and P450BM3m (1.6 µM) concentrations (**Supplementary Fig. 19**). We then tested the same coupled P450BM3m and PTDH reaction system with lauric acid as a natural substrate of P450BM3 under the same conditions. The catalytic efficiency of P450BM3m with lauric acid was two orders of magnitude higher compared to indole (**Table 1**). In addition, the NADPH coupling efficiency of P450BM3m for lauric acid was determined to be at least 10-fold higher than for indole (54.0 ± 17.8% compared to 3.3 ± 0.5%, see **Supplementary Methods**). No inhibitory effect on lauric acid hydroxylation was observed at higher P450 enzyme concentrations in the coupled reaction system with the free enzymes (**Fig. 8b)**. We therefore proceeded to evaluate scaffold co-immobilization of our model two-enzyme system with lauric acid to minimize uncoupling effects upon immobilization.

After testing different P450BM3m and PTDH concentrations (**Supplementary Fig. 20**), we co-immobilized P450BM3m and PTDH at a 1:1 ratio (final enzyme concentrations in the reaction: 3.2 µM each) onto scaffolds. Conjugation to His-SpyC-EutM-SnoopC, His-EutM:SpyC-EutM-SnoopC=3.9:1 or =8.8:1 scaffolds decreased hydroxylauric acid yields 10%, 25%, and 15%, respectively, compared to the free enzyme system, which is comparable to the reduction of lauric acid activity obtained for carrier immobilized P450BM3^34^ (**Fig. 8c**). Scaffold conjugation did not affect hydroxylation site selectivity of P450BM3m with 9-hydroxylauric acid being the major product followed by 10– and 11-hydroxylauric acid **(Supplementary Table 2**). The lower product yields of the scaffolded reaction systems may be due to minor diffusion limitations of the substrate in the cross-linked scaffold material but more likely also to higher electron uncoupling when the two enzymes are co-localized in proximity and thus, exposure of P450BM3m to high local NADPH concentrations. To test recycling and reuse of the scaffolds, we repeated this reaction scaled up to 3 mL with the P450BM3m and PTDH co-immobilized onto the His-EutM:SpyC-EutM-SnoopC=8.8:1 hybrid scaffolds (**Supplementary Fig. 21**) for four cycles. After each 30 min reaction, the scaffolded enzymes were recovered by centrifugation and reused in the next reaction cycle. Hydroxylauric acid yields and protein recovery after each reaction cycle were measured for comparison. While the P450BM3m lost over 60% of its activity after the first reaction cycle and with almost all the enzyme inactivated after three cycles, the hybrid scaffold material proved to be remarkably robust with ∼82% of the protein material recovered after 4 cycles despite the use of a relatively low protein concentration (4.4 mg/mL) in the reaction mix.

Finally, we tested a two-phase reaction system with 20% (v/v) dodecane, 2% DMSO and 20 mM lauric acid (soluble up to 2 mM in water)^42^ for lauric acid conversion with a 4-fold decreased P450BM3m concentration to slow conversion time and allow for longer time course measurements (**Fig. 8d**). Compared to the soluble enzyme system, scaffold immobilization did not benefit initial lauric acid conversion and had only a small effect on the total turnover number (TTN) of the scaffolded P450BM3m which was 3110.7 ± 148.0 compared to a TTN of 3009.9 ± 554.1 for the free enzyme after 12 h of reaction when maximal conversion is reached. The time course of the conversions shows that like in the reuse experiment (**Supplementary Fig. 21**), P450BM3m is rapidly inactivated (even at the reduced enzyme loading), presumably due to the production of H_2_O_2_. This together with a slow mass transfer of lauric acid from the dodecane phase in the bi-phasic reaction^42^ results in only ∼12% conversion of lauric acid.

To increase conversion yields and TTN, additional optimization efforts such as fine tuning enzyme activities, optimizing the reaction system for substrate delivery, use of wild-type P450BM3, catalase incorporation^34^ or separate immobilization of P450BM3m and its NADPH recycling system^36^ will be required to mitigate P450 inactivation that are beyond the scope of this work. For demonstration purposes, the genetically programable protein-based scaffolding system compares favorably with other more mature biocatalyst immobilization methods^34,35,37,53,54^. It allows for the cross-linking of a challenging and complex enzyme without any significant reduction in activity. Furthermore, the macroscale protein carrier material does not appear to significantly impede substrate diffusion, probably due to its assembly into scaffold structures composed of radial microtube bundles with large surface areas for enzyme conjugation and cross-linking.

## CONCLUSIONS

We successfully developed a programmable protein scaffold with two controllable conjugation sites for cargo attachment and cross-linking using the isopeptide bond forming Spy/Snoop-Catcher and –Tag system. By controlling the co-expression levels of unmodified and with conjugation-sites modified EutM, we were able to readily produce macroscale protein materials with different building block ratios for customizable functionalization. We observed that higher levels of unmodified EutM or EutM modified at the N– and C-terminus with the smaller Spy/Snoop-Tags promoted self-assembly into larger and denser scaffold materials composed of microtubular, radial structures. Although the Spy/Snoop-Catcher dual-modified EutM building blocks assemble into smaller, soluble scaffolds, co-expression with unmodified EutM building blocks drives the assembly of large, insoluble hybrid scaffolds that are stable under a range of conditions. Instead of sheet-like structures, the scaffolds form hollow, rolled-up microtubes that increase in diameter and thickness when hybrid scaffolds are co-assembled from His-EutM and SnoopT-EutM-SpyT. These microtubes organize into clusters of radial scaffold particles with large surface areas to which tagged cargo proteins can be rapidly attached, coating surfaces and cross-linking tubes into larger bundles. We furthermore demonstrated that our designed materials are well suited for enzyme immobilization, supporting covalent attachment of a complex P450 enzyme without loss of activity and change of product profile despite immobilization at either end of the enzyme. Co-immobilization of two enzymes is also readily achieved, but for our P450BM3m model system co-immobilization of PTDH exacerbated rapid P450 inactivation, presumably due to high local NADPH concentrations produced by PTDH, leading to creation of reactive uncoupling products. While this challenge may be overcome by additional optimization of the system, this is outside the scope of this work. Nevertheless, the recycling experiments demonstrated that the designed protein scaffolds yield robust, macroscale materials that can be recovered from reactions with >80% yield after several rounds of centrifugation even when used at low protein concentrations – which is an important property for future applications in heterogenous reaction systems.

We observed that the genetic fusion of the larger Spy/Snoop-Catcher domains to cargo enzymes negatively impacted enzyme function. Our scaffolding system can be easily adapted to not only allow for the inclusion of alternative genetically encoded protein-protein interaction mechanisms for attachment, but also separate cross-linking from attachment of enzymes or other proteins. Numerous native or designed proteins can be envisioned to be utilized as cross-linkers to control material properties, while the attachment of enzyme or other proteins will yield functional materials e.g. for biomanufacturing, as functional coatings or conducting and sensing materials. Together with the prospect of eventually programming the fabrication of the entire material in a recombinant system, our robust scaffolding system provides a highly adaptable and customizable platform for the design of a multitude of functional materials.

## METHODS

Methods and materials, including image and statistical analysis and reproducibility statement are provided in the **Supplementary Information.**

## DATA AVAILABILITY

All data generated or analyzed in this study are included in this manuscript and accompanying Supplementary Information, Supplementary Data and Source Data files. Plasmids created in this study can be made available subject to an MTA that can be requested by contacting the corresponding author Prof. Schmidt-Dannert, who will respond to requests within a week. Protein Data Bank data set (PDB: 3I6P) was used to generate EutM hexamer structure representation in **Fig. 1 & 2**.

## ACKNOWLEDGEMENTS

This research was sponsored by grants from the National Science Foundation (CBET-1916030) and Department of Energy Advanced Research Project Agency (ARPA-E) (DE-AR0001510) and supported by seed grants from the BioTechnology Institute at the University of Minnesota.

## AUTHOR CONTRIBUTIONS

R.Z. contributed to the experimental design, created genetic constructs, isolated, and characterized scaffold, performed microscopy, enzyme assays, analyzed data and wrote the manuscript. S.-Y.K. contributed to the microscopy analysis of the scaffolds and writing of the manuscript. F.G. contributed to the GC analysis and manuscript writing. E.L.P. contributed to the design and construction of plasmids and the writing of the manuscript. C.S.-D. conceived, devised, conceptualized, and directed the overall project, analyzed data, and wrote the manuscript together with R.Z.

## REFERENCES

1 Miserez, A., Yu, J. & Mohammadi, P. Protein-based biological materials: molecular design and artificial production. Chemical Reviews 123, 2049–2111, doi:10.1021/acs.chemrev.2c00621 (2023).

2 Seo, M.-J. & Schmidt-Dannert, C. Organizing multi-enzyme systems into programmable materials for biocatalysis. Catalysts 11, 409, doi:10.3390/catal11040409 (2021).

3. 3 Wallin, K., Zhang, R. & Schmidt-Dannert, C. in Engineered Living Materials (ed Wil V. Srubar Iii) 51–94 (Springer International Publishing, 2022).

4 Shen, Y. et al. From protein building blocks to functional materials. ACS Nano 15, 5819–5837, doi:10.1021/acsnano.0c08510 (2021).

5 Kortemme, T. De novo protein design – From new structures to programmable functions. Cell 187, 526–544, doi:10.1016/j.cell.2023.12.028 (2024).

6 Mout, R. et al. De novo design of modular protein hydrogels with programmable intra– and extracellular viscoelasticity. Proceedings of the National Academy of Sciences 121, e2309457121, doi:10.1073/pnas.2309457121 (2024).

7 Hilditch, A. T. et al. Assembling membraneless organelles from de novo designed proteins. Nature Chemistry 16, 89–97, doi:10.1038/s41557-023-01321-y (2024).

8 Zhang, W. et al. Rationally designed protein building blocks for programmable hierarchical architectures. Frontiers in Chemistry 8, doi:10.3389/fchem.2020.587975 (2020).

9 Hirschi, S., Ward, T. R., Meier, W. P., Mueller, D. J. & Fotiadis, D. Synthetic biology: bottom-up assembly of molecular systems. Chemical Reviews 122, 16294–16328, doi:10.1021/acs.chemrev.2c00339 (2022).

10 Pokhrel, A., Kang, S.-y. & Schmidt-Dannert, C. Ethanolamine bacterial microcompartments: from structure, function studies to bioengineering applications. Current Opinion in Microbiology 62, 28–37, doi:10.1016/j.mib.2021.04.008 (2021).

11 Zhang, G., Schmidt-Dannert, S., Quin, M. B. & Schmidt-Dannert, C. Protein-based scaffolds for enzyme immobilization. Methods Enzymology 617, 323–362, doi:10.1016/bs.mie.2018.12.016 (2019).

12 Zhang, G., Quin, M. B. & Schmidt-Dannert, C. Self-assembling protein scaffold system for easy *in vitro* coimmobilization of biocatalytic cascade enzymes. ACS Catalysis 8, 5611–5620, doi:10.1021/acscatal.8b00986 (2018).

13 Schmidt-Dannert, S., Zhang, G., Johnston, T., Quin, M. B. & Schmidt-Dannert, C. Building a toolbox of protein scaffolds for future immobilization of biocatalysts. Applied Microbiology and Biotechnology 102, 8373–8388, doi:10.1007/s00253-018-9252-6 (2018).

14 Zhang, G., Johnston, T., Quin, M. B. & Schmidt-Dannert, C. Developing a protein scaffolding system for rapid enzyme immobilization and optimization of enzyme functions for biocatalysis. ACS Synthetic Biology 8, 1867–1876, doi:10.1021/acssynbio.9b00187 (2019).

15 Veggiani, G. et al. Programmable polyproteams built using twin peptide superglues. Proceedings of the National Academy of Sciences 113, 1202–1207, doi:10.1073/pnas.1519214113 (2016).

16 Manea, F., Garda, V. G., Rad, B. & Ajo-Franklin, C. M. Programmable assembly of 2D crystalline protein arrays into covalently stacked 3D bionanomaterials. Biotechnology and Bioengineering 117, 912–923, doi:10.1002/bit.27261 (2020).

17 Beyer, N. et al. P450(BM3) fused to phosphite dehydrogenase allows phosphite-driven selective oxidations. Applied Microbiology Biotechnology 101, 2319–2331, doi:10.1007/s00253-016-7993-7 (2017).

18 Whitehouse, C. J. C., Bell, S. G. & Wong, L.-L. P450BM3 (CYP102A1): connecting the dots. Chemical Society Reviews 41, 1218–1260, doi:10.1039/C1CS15192D (2012).

19 Bolivar, J. M., Woodley, J. M. & Fernandez-Lafuente, R. Is enzyme immobilization a mature discipline? Some critical considerations to capitalize on the benefits of immobilization. Chemical Society Reviews 51, 6251–6290, doi:10.1039/d2cs00083k (2022).

20 Chen, Z. et al. Self-assembly systems to troubleshoot metabolic engineering challenges. Trends in Biotechnology, doi:10.1016/j.tibtech.2023.06.009.

21 Caparco, A. A., Dautel, D. R. & Champion, J. A. Protein mediated enzyme immobilization. Small 18, doi:10.1002/smll.202106425 (2022).

22 Jäger, V. D. et al. Tailoring the properties of (catalytically)-active inclusion bodies. Microbial cell factories 18, 33, doi:10.1186/s12934-019-1081-5 (2019).

23 Kowalski, A. E. et al. Porous protein crystals as scaffolds for enzyme immobilization. Biomaterials Science 7, 1898–1904, doi:10.1039/C8BM01378K (2019).

24 Diener, M., Kopka, B., Pohl, M., Jaeger, K.-E. & Krauss, U. Fusion of a coiled-coil domain facilitates the high-level production of catalytically active enzyme inclusion bodies. ChemCatChem 8, 142–152, doi:10.1002/cctc.201501001 (2016).

25 Yin, L., Guo, X., Liu, L., Zhang, Y. & Feng, Y. Self-assembled multimeric-enzyme nanoreactor for robust and efficient biocatalysis. ACS Biomaterials Science & Engineering 4, 2095–2099, doi:10.1021/acsbiomaterials.8b00279 (2018).

26 Ledesma-Fernandez, A., Velasco-Lozano, S., Santiago-Arcos, J., Lopez-Gallego, F. & Cortajarena, A. L. Engineered repeat proteins as scaffolds to assemble multi-enzyme systems for efficient cell-free biosynthesis. Nature Communications 14, doi:10.1038/s41467-023-38304-z (2023).

27 Noël, C. R., Cai, F. & Kerfeld, C. A. Purification and characterization of protein nanotubes assembled from a single bacterial microcompartment shell subunit. Advanced Materials Interfaces 3, 1500295, doi:10.1002/admi.201500295 (2016).

28 Kaczmarczyk, A., Vorholt, J. A. & Francez-Charlot, A. Cumate-inducible gene expression system for sphingomonads and other alphaproteobacteria. Applied and Environmental Microbiology 79, 6795–6802, doi:10.1128/AEM.02296-13 (2013).

29 Salis, H. M., Mirsky, E. A. & Voigt, C. A. Automated design of synthetic ribosome binding sites to control protein expression. Nature Biotechnology 27, 946–950, doi:10.1038/nbt.1568 (2009).

30 Seo, S. O. & Schmidt-Dannert, C. Development of a synthetic cumate-inducible gene expression system for *Bacillus*. Applied Microbiology and Biotechnology 103, 303–313, doi:10.1007/s00253-018-9485-4 (2019).

31 Vick, J. E. et al. Optimized compatible set of BioBrick vectors for metabolic pathway engineering. Applied Microbiology and Biotechnology 92, 1275–1286, doi:10.1007/s00253-011-3633-4 (2011).

32 Faulkner, M., Zhao, L. S., Barrett, S. & Liu, L. N. Self-assembly stability and variability of bacterial microcompartment shell proteins in response to the environmental change. Nanoscale Research Letters 14, doi:10.1186/s11671-019-2884-3 (2019).

33 Fansher, D. J., Besna, J. N., Fendri, A. & Pelletier, J. N. Choose your own adventure: a comprehensive database of reactions catalyzed by cytochrome P450 BM3 variants. ACS Catal, 5560–5592, doi:10.1021/acscatal.4c00086 (2024).

34 Valikhani, D., Bolivar, J. M., Dennig, A. & Nidetzky, B. A tailor-made, self-sufficient and recyclable monooxygenase catalyst based on coimmobilized cytochrome P450 BM3 and glucose dehydrogenase. Biotechnology and Bioengineering 115, 2416–2425, doi:10.1002/bit.26802 (2018).

35 Solé, J., Caminal, G., Schürmann, M., Alvaro, G. & Guillén, M. Co-immobilization of P450 BM3 and glucose dehydrogenase on different supports for application as a self-sufficient oxidative biocatalyst. Journal of Chemical Technology and Biotechnology 94, 244–255, doi:10.1002/jctb.5770 (2019).

36 Maurer, S. C., Schulze, H., Schmid, R. D. & Urlacher, V. Immobilisation of P450 BM-3 and an NADP+ cofactor recycling system: towards a technical application of heme-containing monooxygenases in fine chemical synthesis. Advanced Synthesis & Catalysis 345, 802–810, doi:10.1002/adsc.200303021 (2003).

37 Valikhani, D., Bolivar, J. M. & Pelletier, J. N. An overview of cytochrome P450 immobilization strategies for drug metabolism studies, biosensing, and biocatalytic applications: challenges and opportunities. ACS Catalysis 11, 9418–9434, doi:10.1021/acscatal.1c02017 (2021).

38 Bahrami, A. et al. Kinetics of enzymatic hydroxylation by free and MNPs-immobilized NADH-dependent cytochrome P450 BM3 from *Bacillus megaterium*. Industrial & Engineering Chemistry Research 58, 808–815, doi:10.1021/acs.iecr.8b04513 (2019).

39 Buergler, M. B., Dennig, A. & Nidetzky, B. Process intensification for cytochrome P450 BM3-catalyzed oxy-functionalization of dodecanoic acid. Biotechnology and Bioengineering 117, 2377–2388, doi:10.1002/bit.27372 (2020).

40 Li, Q.-S., Schwaneberg, U., Fischer, P. & Schmid, R. D. Directed evolution of the fatty-acid hydroxylase P450 BM-3 into an indole-hydroxylating catalyst. Chemistry – A European Journal 6, 1531–1536, doi:10.1002/(SICI)1521-3765(20000502)6:9<1531::AID-CHEM1531>3.0.CO;2-D (2000).

41 Appel, D., Lutz-Wahl, S., Fischer, P., Schwaneberg, U. & Schmid, R. D. A P450BM-3 mutant hydroxylates alkanes, cycloalkanes, arenes and heteroarenes. Journal of Biotechnology 88, 167–171, doi:10.1016/s0168-1656(01)00249-8 (2001).

42 Maurer, S. C. et al. Catalytic hydroxylation in biphasic systems using CYP102A1 mutants. Advanced Synthesis & Catalysis 347, 1090–1098, doi:10.1002/adsc.200505044 (2005).

43 Woodyer, R., Zhao, H. & van der Donk, W. A. Mechanistic investigation of a highly active phosphite dehydrogenase mutant and its application for NADPH regeneration. The FEBS Journal 272, 3816–3827, doi:10.1111/j.1742-4658.2005.04788.x (2005).

44 Beyer, N., Kulig, J. K., Fraaije, M. W., Hayes, M. A. & Janssen, D. B. Exploring PTDH– P450BM3 variants for the synthesis of drug metabolites. ChemBioChem 19, 326–337, doi:10.1002/cbic.201700470 (2018).

45 Huang, W.-C. et al. Filling a Hole in Cytochrome P450 BM3 Improves Substrate Binding and Catalytic Efficiency. Journal of Molecular Biology 373, 633–651, doi:10.1016/j.jmb.2007.08.015 (2007).

46 Li, Q.-S., Ogawa, J., Schmid, R. D. & Shimizu, S. Indole hydroxylation by bacterial cytochrome P450 BM-3 and modulation of activity by cumene hydroperoxide. Bioscience, Biotechnology, and Biochemistry 69, 293–300, doi:10.1271/bbb.69.293 (2005).

47 Rowlatt, B. et al. Chain length-dependent cooperativity in fatty acid binding and oxidation by cytochrome P450BM3 (CYP102A1). Protein & Cell 2, 656–671, doi:10.1007/s13238-011-1082-6 (2011).

48 Kong, F. et al. Evolving a P450BM3 peroxygenase for the production of indigoid dyes from indoles. ChemCatChem 14, e202201151, doi:10.1002/cctc.202201151 (2022).

49 Mendoza-Avila, J., Chauhan, K. & Vazquez-Duhalt, R. Enzymatic synthesis of indigo-derivative industrial dyes. Dyes and Pigments 178, 108384, doi:10.1016/j.dyepig.2020.108384 (2020).

50 Jeffreys, L. N. et al. Characterization of the structure and interactions of P450 BM3 using hybrid mass spectrometry approaches. Journal of Biological Chemistry 295, 7595–7607, doi:10.1074/jbc.RA119.011630 (2020).

51 Cirino, P. C. & Arnold, F. H. Regioselectivity and activity of cytochrome P450 BM-3 and mutant F87A in reactions driven by hydrogen peroxide. Advanced Synthesis & Catalysis 344, 932–937, doi:10.1002/1615-4169 (200210)344:9<932::AID-ADSC932>3.0.CO;2-M (2002).

52 Vidal-Limón, A., Águila, S., Ayala, M., Batista, C. V. & Vazquez-Duhalt, R. Peroxidase activity stabilization of cytochrome P450BM3 by rational analysis of intramolecular electron transfer. Journal of Inorganic Biochemistry 122, 18–26, doi:10.1016/j.jinorgbio.2013.01.009 (2013).

53 Bahrami, A. et al. Noncovalent immobilization of optimized bacterial cytochrome P450 BM3 on functionalized magnetic nanoparticles. Industrial & Engineering Chemistry Research 56, 10981–10989, doi:10.1021/acs.iecr.7b02872 (2017).

54 Nöth, M. et al. MicroGelzymes: pH-independent immobilization of cytochrome P450 BM3 in microgels. Biomacromolecules 21, 5128–5138, doi:10.1021/acs.biomac.0c01262 (2020).

