## Supplementary Information for "Design of a genetically programmable and customizable protein scaffolding system for the hierarchical assembly of robust, functional macroscale materials"

140 Gortner Laboratory

University of Minnesota

1479 Gortner Avenue

St. Paul, MN 55108

### Supplementary Methods

#### Materials and chemicals

All chemical reagents were purchased from Sigma-Aldrich (St. Louis, MO, USA) unless otherwise noted. Lauric acid (≥99.0% GC, catalog #61610), palmitic acid (≥98.5%, catalog #76120), and 12-hydroxylauric acid (≥97%, catalog #55499) were obtained from Honeywell Fluka™ (Morris Plain, NJ, USA). Except for the PCRBio Verifi Mix from Genesee Scientific (Morrisville, NC, USA) used for colony PCR, all other molecular biology reagents (HiFi® DNA Assembly Master mix for Gibson assembly and other enzymes) were obtained from New England Biolabs (Ipswhich, MA, USA). Oligonucleotides were purchased from Integrated DNA Technologies, Inc. (Coralville, IA, USA).

For purification and PAGE analysis of proteins, Spectra/Por Dialysis Tubing (MWCO 6-8 kDa) from Spectrum Life Sciences (Rancho Dominguez, CA, USA) was used for dialysis and HisPur™ Ni-NTA resin from ThermoFisher™ Scientific (Waltham, MA, USA) for metal affinity chromatography. For P450BM3m purification, Roche cOmplete™ Protease Inhibitor Cocktail tablets purchased from Sigma-Aldrich were used. The Pierce™ BCA Protein Assay Kit from ThermoFisher was used to quantify protein concentrations. Except for Coomassie brilliant blue from Sigma-Aldrich, all other reagents for PAGE (TEMED, Precision Plus Protein™ prestained protein standard (catalog #161-0373), 30% acrylamide and bis-acrylamide solution (37.5:1)) were obtained from Bio-Rad (Hercules, CA, USA). A Milli-Q water purification system (MilliporeSigma, Burlington, MA, USA) was used to filter deionized water to prepare ultrapure water with a final electrical resistance higher than 18.2 MΩ $cm^{-1}$.

#### Bacterial strains, media, and general cloning methods

Cloning and plasmid propagation were done in *E. coli* TOP10 (Invitrogen, Carlsbad, CA, USA) while *E. coli* BL21 (DE3) (New England Biolabs, Ipswhich, MA, USA) was used for expression of proteins for purification. *E. coli* strains were grown in LB (Luria broth; tryptone 10 g/L, NaCl 5 g/L, yeast extract 10 g/L) medium supplemented with appropriate antibiotics (100 µg/mL ampicillin (LB-Amp) or 30 µg/mL kanamycin (LB-Km)) for plasmid maintenance. For cytochrome P450BM3m protein expression and purification, LB medium was replaced with Hyper Broth™ (Athenea ES, Baltimore, MD, USA).

Plasmid transformation into *E. coli* followed standard molecular biology techniques. Transformants were confirmed by colony PCR and all gene and plasmid sequences were verified by Sanger sequencing (ACGT Inc., Wheeling, IL, USA) and complete plasmid Nanopore sequencing (Plasmidsaurus, Eugene, OR, USA).

### Plasmid construction

Plasmids were constructed using a combination of methods, including Gibson Assembly (HiFi® DNA assembly kit from New England Biolabs), T5 exonuclease-dependent assembly[1] for fragment assembly and site-directed mutagenesis (Q5® kit, New England Biolabs) for short insertions, deletions and mutations as described previously[2,3].

For amplification and cloning of His-tagged GFP cargo and EutM with and without SpyTag/SpyCatcher fusions, we used previously described plasmids as templates and backbones (pCT5BB or pET28a) [4]. The PTDH nucleotide sequence and SnoopCatcher sequence were synthesized by Genewiz® (South Plainfield, NJ. USA). The wildtype P450BM3 gene was amplified from genomic DNA isolated from *Bacillus megaterium*[3,5] and three amino acid substitutions (Ala74Gly, Phe87Val, Leu188Gln) were introduced into the cloned gene by site-directed mutagenesis to yield the indigo producing variant P450BM3m[5]. Shorter Snoop/SpyTag and GS-linker sequences and ribosome binding sites were inserted by site-directed mutagenesis. All plasmids used and constructed are listed in **Supplementary Table 3**. Amino acid sequences and encoding nucleotide sequences for EutM scaffolds, GFP cargo proteins and enzymes are provided in **Supplementary Table 4 & 5**.

Briefly, cargo protein cloning started with the assembly of GFP into the *Nde*I site of pET28a, followed by the insertion of Spy/Snoop Tag/Catcher fusions and GS-linkers up- and downstream of GFP. GFP was replaced by PTDH or P450BM3m to yield the corresponding cargo protein expression plasmids. Plasmids for EutM scaffold protein expression were constructed using pCT5BB-His-EutM[2,4,6] as a template to create pCT5BB-His-SnoopT-EutM-SpyT and pCT5BB-His-SpyC-EutM-SnoopC plasmids for the expression of dual-modified EutM proteins. Hybrid scaffold expression plasmids were constructed by amplifying His-EutM from pCT5BB-His-EutM along with its upstream RBS (strong native RBS in pCT5BB referred to as RBSA) and inserting it upstream of the RBSA site of the dual-modified EutM expression cassettes. Site-directed mutagenesis was used to change His-tags and delete RBSs. RBS with different strengths in addition to the native RBSA from pCT5BB were selected based on previous work [7] and predictions using the online RBS calculator developed by the Salis group (https://salislab.net/software/predict_rbs_calculator) **(Supplementary Table 1)**.

### Protein expression in *E. coli*

For the expression of EutM scaffolds from the cumate inducible promoter on pCT5BB plasmids, single colonies of *E. coli* BL21 (DE3) transformants were inoculated into 50 mL of LB-Amp and grown overnight (30 ºC, 180 rpm). Overnight cultures were diluted 100-fold into fresh LB-Amp (200 mL in 1 L flasks or 400 mL in 2 L flasks) and grown at 37 ºC, 180 rpm until $OD_{600}$ = 0.6-1.0 when protein expression was induced with 50 µM cumate. Induced cultures were grown for 16-20 h at 37 ºC, 180 rpm and cells harvested by centrifugation at 4,000 x*g* for 25 mins at 4 ºC. Cell pellets were stored at -80 ºC until needed.

For the expression of GFP cargo proteins from the T7-promoter on pET28a plasmids, LB-Amp was replaced with LB-Km and protein expression and cell harvest followed the same procedure, except that protein expression was induced with 0.1 mM isopropyl β-D-1-thiogalactopyranoside (IPTG). For PTDH cargo protein expression from the T7-promoter on a pET28a plasmid, the temperature of the overnight culture was lowered to 25 °C prior to induction and the induced cultures were grown for 16-20 h at 180 rpm until harvest of cell pellets which were stored at -80 ºC until needed.

For cytochrome P450BM3m cargo protein expression from the T7-promoter on pET28a plasmids, *E. coli* BL21(DE3) transformants were grown overnight in 50 mL LB-Km (30 ºC, 180 rpm). Overnight cultures were diluted 1:20 into Hyper Broth™ (AthenaES®, catalog #0107, 400 mL in 2 L flasks) supplemented with 30 µg/mL kanamycin, 1 mM $MgSO_4$, 0.1 mM $FeCl_2$, and 1x trace elements[8] and grown (37 ºC, 180 rpm) until $OD_{600}$ = 1.0-1.5 (~3-4 h). Cultures were cooled down in an ice-water bath for 20 min and then induced with 0.1 mM IPTG. To ensure sufficient heme biosynthesis, δ-aminolevulinic acid was added (1 mM final concentration) at this time. Induced cultures were grown at 22 ºC and 140 rpm for 16-20 h until harvest and storage as described above.

**EutM scaffold and GFP cargo protein purification**

For the purification of the soluble dual-modified EutM scaffolds and GFP cargo proteins, *E. coli* cells were suspended in lysis buffer without urea (20 mM imidazole, 50 mM Tris-HCl, 250 mM NaCl, pH 8.0) and disrupted by sonication (30 mins, power 40%, pulse on 1 s, and pulse off 2 s on ice with a Branson Sonifier). The lysed cells were centrifuged (10,000 x*g*, 30 min, 4 °C) and the His-tagged proteins in the supernatant purified using the Batch Protocol with the HisPur™ Ni-NTA resin according to the manufacturer's instructions (ThermoFischer™). Briefly, clarified supernatant was mixed with resin and incubated for 1 h at 4 ºC. The mixture was then loaded onto a gravity-flow column, washed with lysis buffer and bound proteins eluted with five resin-bed volumes of elution buffer (250 mM imidazole, 50 mM Tris-HCl, 250 mM NaCl, pH 8.0).

For the purification of wild-type, hybrid-EutM scaffolds, samples for assembly testing and Native

PAGE analysis of dual-modified EutM scaffolds (**Supplemental Fig. 1&3)**, 4 M urea was added to both lysis and elution buffers to disassemble and solubilize large EutM scaffolds. The purified, eluted proteins were then concentrated with an Amicon® Ultra Centrifugal Filter (3 kDa MWCO) to a concentration of 20-40 mg/mL if needed. To remove imidazole and/or urea, purified and concentrated proteins were dialyzed against 50 mM Tris-HCl buffer (pH 7.5) (or other buffers where indicated) at 4 ºC overnight using Spectra/Por™ dialysis tubing (6-8 kDa MWCO).

**PTDH and P450BM3m cargo enzyme purification**

*E. coli* cells were resuspended and lysed by sonication as described above, except that for P450BM3m purification the cell density for lysis was controlled to 4 mL buffer per gram of cell wet weigh and 1 mg/mL lysozyme and Roche cOmplete™ Protease Inhibitor was added to the buffer according to the manufacturer's instructions. Proteins were purified by Ni-NTA affinity chromatography as described above with the exception that the column was washed with a buffer (50 mM Tris-HCl, 250 mM NaCl, pH 8.0) containing 45 mM imidazole and proteins were eluted with five volumes of elution buffer. The buffer of the eluted protein samples was exchanged to a 0.1 M sodium phosphate buffer (pH 7.0) with a PD-10 desalting column (GE HealthCare, Buckinghamshire, UK). Proteins were concentrated to 40-80 mg/mL with an Amicon® Ultra Centrifugal Filter (10 kDa MWCO), mixed 1:1 v/v with glycerol and aliquots flash-frozen in liquid-nitrogen for storage at -80 ºC until needed. All protein purifications were performed at least three times from fresh transformed *E. coli* strains.

**SDS-PAGE and protein concentration analysis**

Purity and protein sample compositions of scaffolds, cargo proteins and enzymes were analyzed by 15% SDS-PAGE following standard methods with samples diluted 6x with loading buffer and denatured for 20 min at 100 ºC prior to loading. Protein concentrations were measured with the Pierce BCA assay kit using the manufacturer's 60 ºC protocol.

**Native PAGE analysis of EutM scaffolds**

Purified EutM scaffold proteins were normalized to 2 mg/mL with elution buffer containing 4 M urea and then mixed with 2x Native PAGE Sample Buffer (62.5 mM Tris-HCl, 40% glycerol, 0.01% bromophenol blue, pH 6.8), separated on 4-15% Mini-PROTEAN® TGX Stain-Free™ Protein Gels (Bio-Rad catalog #4568083) and stained with Bio-Safe™ Coomassie Stain (Bio-Rad catalog #1610786) (**Supplementary Fig. 1**)

**GFP cargo-crosslinking to different EutM scaffolds**

Purified GFP cargo proteins (dual-modified and unmodified control) and EutM scaffold or hybrid scaffold proteins (20-40 mg/mL in 0.1 M sodium phosphate buffer (pH 7.0)) were mixed with the iso-peptide bond forming partners (e.g. 50 µM His-SpyT-GFP-SnoopT with 50 µM His-SpyC-EutM-SnoopC scaffold or with 50 µM SpyC-EutM-SnoopC in the His-EutM:SpyC-EutM-SnoopC scaffold). Reactions were performed in a 0.1 M sodium phosphate buffer (pH 7.0) (200 µL final volume) prior to mixing with sample buffer for SDS-PAGE analysis (**Fig. 2b**, **Supplementary Fig. 9**) or for imaging by microscopy or TEM (**Figs. 5 & 6**, **Supplementary Fig. 12**). To characterize cargo cross-linking under different conditions, reactions were performed with a 1:1 molar ratio of 50 µM cross-linking partner proteins under different conditions, including for 1 h at different temperatures (4, 25, 30, and 37 °C) and at 25 °C for up to 24 h with samples taken at different intervals (**Supplementary Fig. 2**). In addition, reactions were performed with different molar ratios of partner proteins by mixing 50 µM of the EutM scaffold partner with 12.5, 25, 50, 100, and 200 µM GFP cargo protein for 1 h at 25 °C and 180 rpm prior to analysis by microscopy (**Supplementary Figs. 10 & 11**).

The final scaffold and GFP cargo protein concentrations in the samples for these reactions were: i) for 50 µM EutM scaffolds: 1.7 mg/mL for His-SpyC-EutM-SnoopC (34.3 kDa) (w/o His-Tag in hybrid scaffolds = 32.7 kDa, 1.6 mg/mL), 0.8 mg/mL for His-SnoopT-EutM-SpyT (15.8 kDa) (w/o His-Tag in hybrid scaffolds = 14.5 kDa, ~ 0.7 mg/mL); ii) for 50 µM of dual-modified EutM in the hybrid scaffold designs containing different ratios of His-EutM (11.5 kDa, 50 µM = 0.6 mg/mL): 3.9 mg/mL His-EutM:SpyC-EutM-SnoopC=3.9:1, 6.7 mg/mL His-EutM:SpyC-EutM-SnoopC=8.8:1, 2.6 mg/mL His-EutM:SnoopT-EutM-SpyT=3.3:1, 5.1 mg/mL His-EutM:SnoopT-EutM-SpyT=7.6:1, iii) for the different concentrations of the GFP cargo protein: 0.7, 1.3, 2.6, 5.2, 10.4 mg/mL corresponding to 12.5, 25, 50, 100, 200 µM His-SnoopC-GFP-SpyC, 0.4, 0.8, 1.7, 3.3, 6.6 mg/mL corresponding to 12.5, 25, 50, 100, 200 µM His-SnoopT-GFP-SpyT, 1.5 mg/mL corresponding to 50 µM His-GFP.

#### ***In vitro*** **scaffold co-assembly of EutM scaffold building blocks**

To investigate *in vitro* co-assembly of purified EutM scaffold building blocks into hybrid scaffolds, purified scaffold building blocks (2 mg/mL) in elution buffer with urea were mixed at a 5:1 molar ratio (His-EutM:dual-modified EutM) and dialyzed against a 50 mM Tris-HCl buffer (pH 7.5). The assembled scaffolds were then analyzed by Native PAGE and compared to controls with single building blocks (**Supplementary Fig. 1**). The same samples were also analyzed by SDS-PAGE after separating soluble (S) and insoluble scaffolds as pellet (P) by centrifugation at 12,000 x*g* for 2 min (**Supplementary Fig. 3**).

### Hybrid scaffold characterization

The total yield of hybrid EutM scaffolds from 200 mL cultures was determined by measuring the protein concentration in the eluted, purified protein fraction (5 mL) after metal affinity chromatography. Expression, purification, and subsequent characterization experiments were done with samples obtained from three different cultures for each genetic construct. The EutM scaffold building block ratios in the different hybrid scaffolds were determined by measuring protein concentrations of EutM proteins by densitometry of SDS-PAGE gels with a standard curve of 0.1-1.0 mg/mL of purified His-EutM. Quantification was performed using the ImageJ (version 1.530) software following the protocol described by the Starr Lab and originally written by Luke Miller[9] (**Fig. 3b, Supplementary Fig. 4**). Representative SDS-PAGE gels are shown in **Supplementary Fig. 5**.

To assess scaffold assembly behavior, purified hybrid scaffolds in elution buffer with urea were normalized to 2 mg/mL with elution buffer and then dialyzed against 50 mM Tris-HCl buffer (pH 7.5) at 4 ºC overnight using Spectra/Por™ dialysis tubing (6-8 kDa MWCO). During dialysis, insoluble, larger scaffolds assembled resulting in the formation of a white protein material. These insoluble scaffolds (Pellet) were separated from soluble scaffolds by centrifugation at 12,000 x*g* for 2 min at room temperature. The protein concentration of the soluble fraction (S) and the insoluble scaffolds (P, Pellet) after resuspension in 50 mM Tris-HCl buffer (pH 7.5) was measured to calculate the percentage of insoluble assembled scaffolds of the total scaffold protein concentration as: [P] / [P+ S] in % (**Fig. 3b, Supplementary Fig. 4**).

To investigate the influence of pH, temperature, and NaCl concentrations on scaffold assembly, purified hybrid scaffolds in elution buffer were normalized to 3 mg/mL with elution buffer, and first dialyzed as described above into the following buffers: 0.1 M sodium acetate (pH 5), 0.1 M sodium phosphate (pH 6, pH 7, or pH 7.5), and 0.1 M Tris-HCl (pH 7.5, pH 8 or pH 9). The buffer exchanged scaffolds were then normalized to 2 mg/mL with the same buffers without and with NaCl to achieve final concentrations of 0, 100, or 250 mM NaCl. Samples were then aliquoted and incubated for 24 h at 4, 25, 30, and 37 ºC. Scaffold assembly behavior was then measured as described above to quantify the percentage of insoluble scaffold protein in the samples (**Fig. 4, Supplementary Figs. 7 & 8**).

### Phase contrast and fluorescence light microscopy

For imaging cargo loading onto EutM scaffolds (**Figs. 5 & 6b**, **Supplementary Figs. 10 & 11**), 10 µL of protein sample was loaded onto a glass slide and covered with a coverslip. A Leica DM4000 microscope controlled by the Leica Application Suite X (version 3.7.4.23463) and equipped with a 100x oil-immersion objective and filters for phase contrast or fluorescence imaging was used for slide examination and image capture. GFP fluorescence was visualized using a L5 fluorescence cube (BP 480/40, dichromatic mirror 505, suppression filter BP527/30) with a 1.0 second exposure time.

#### Confocal fluorescence microscopy

For imaging of 3D-features of EutM scaffolds cross-linked with GFP cargo (**Fig. 6b**), 10 µL of protein sample was applied to a glass slide and covered with a coverslip for examination with a Nikon A1plus Ti2 microscope equipped with a 60x 1.42 oil lambda D objective (University of Minnesota Imaging Center). The refraction index was set to 1.51 and a GFP fluorescence cube (excitation 488 nm, emission 525 nm) with a pinhole size equal to 35.76 was used for illumination. Images were captured using a Nikon A1plus camera and Nikon's NIS Elements software (version 5.30.02). Z-stacks were acquired using the microscope's ZDrive for capturing 52-55 slices with a step size of 0.1 µm.

#### Transmission Electron Microscopy (TEM)

Scaffold protein samples were diluted to 1 mg/mL in 0.1 M sodium phosphate buffer (pH 7.0) and then, 10 µL of samples were dropped onto a 200 mesh Formvar/Carbon grid (Electron Microscopy Sciences) and let adsorb for 5 min. Fluid was removed and 10 µL Trump's fixative (Electron Microscopy Sciences, Hatfield, PA, catalog #11750) applied for 5 min, then removed (with filter paper) and the grid rinsed three times with ultra-pure water. A drop of 1% aqueous uranyl acetate was applied to the grids and immediately removed to avoid overstaining. A JEOL-JEM1400Plus transmission electron microscope with a LaB6 tungsten filament at 60 kV was used to examine grids. Images were captured using an Advanced Microscopy Techniques XR16 camera with an AMT capture Engine software (version 7.0.0187) (University of Minnesota Imaging Center) (**Figs. 3c, 6a & 8a, Supplementary Figs. 6, 12, & 18**).

#### Image analysis

Images were cropped and scale bars added using ImageJ (version 1.54f). Confocal images were analyzed using Nikon's NIS Elements AR Analysis software (version 5.42.04) for 3D-reconstruction of the observed structures. The screenshot function of the analysis software was used to capture images for the slice and volume views of the reconstructed structures (**Fig. 6b**).

#### Quantification of P450BM3m concentration

Carbon monoxide (CO) difference spectra analysis[10] was used to determine the concentration of active P450BM3m. For this, purified P450BM3m enzyme was diluted into 0.1 M sodium phosphate buffer (pH 7.0) to a final concentration of 0.5-1 mg/mL and absorbance between 400-500 nm was measured with a 200 µL sample aliquot using a Varioskan LUX Multimode microplate reader (path length = 0.58 cm) (ThermoFisher Scientific). The P450BM3m solution was then saturated with CO (bubbling gas for 40-60 sec) and reduced by the addition of few grains of sodium hydrosulfite ($Na_2S_2O_4$). Another 200 µL sample aliquot was taken and the absorbance was measured again between 400 and 500 nm. Functional P450BM3m concentration was calculated with this reading using the following equation: (Absorption at 450 nm – Absorption at 490 nm)/ε x d; ε = 91 $mM^{-1}$ $cm^{-1}$ at 450 nm; d = path length[11,12]. Measurements were performed in triplicate with three separate samples.

**P450BM3m kinetic measurements**

Kinetic parameters (**Table 1, Supplementary Fig. 13**) for P450BM3m (with and without Spy/Snoop-Tag or -Catcher fusions) was determined spectrophotometrically with a Varioskan LUX multimode microplate reader by monitoring NADPH concentrations at 340 nm (ε = 6.22 $mM^{-1}$ $cm^{-1}$) with indole or lauric acid as substrates. For reactions with indole, indigo formation was also spectrophotometrically quantified at 670 nm (ε = 3.9 $mM^{-1}$ $cm^{-1}$). Reactions were started by the addition of NADPH.

Assays were performed with four separate replicate samples and corresponding no-enzyme control reactions at 30 ºC and 600 rpm (pulsed with 10s on and 10s off and low force setting) with a Varioskan LUX multimode microplate reader. Assays were carried out in 0.1 M sodium phosphate buffer (pH 7.0) with a total reaction volume of 200 µL per sample. To determine the $k_{cat}/K_m$ for lauric acid and indole, the reactions contained: 40 µL of P450BM3m (final concentration 0.4 µM), 2 µL of 5-500 mM indole in DMSO (final indole concentration 0.05-5 mM) or 16 µL of 0.625-25 mM lauric acid in DMSO (final lauric acid concentration 0.05-2 mM) and 10 µL of 5 mM NADPH (final NADPH concentration 0.25 mM). To determine the $k_{cat}/K_m$ of the NADPH cofactor, the NADPH concentration was varied from 0.025 to 0.25 mM NADPH (10 µL of 0.5-5 mM NADPH) and 5 mM indole (2 µL 500 mM indole in DMSO) was used as the substrate. Kinetic parameters were calculated using the Hill fitting function of Origin (version 2022b).

**PTDH kinetic measurements**

Enzyme activity for PTDH (with and without Spy/Snoop-Tag or -Catcher fusions) was determined by monitoring NADPH concentrations at 340 nm (ε = 6.22 $mM^{-1}$ $cm^{-1}$) and at 30 ºC and 600 rpm (pulsed with 10s on and 10s off and a low force setting) with a Varioskan LUX multimode microplate reader. Assays were performed with four separate samples and with no-enzyme control reactions. Assays were done in a total volume of 200 µL containing: 140 µL of 0.1 M sodium phosphate buffer (pH 7.0); 10 µL of 20 mM sodium phosphite ($Na_2HPO_3 \cdot 5H_2O$ final concentration 1 mM); 40 µL PTDH (final concentrations: 0.03 µM for His-SpyT-PTDH-SnoopT, 0.05 µM for His-PTDH, 0.25 µM for the much less active His-SnoopC-PTDH-SpyC) and 10 µL of 5 mM $NADP^+$ (final concentration 250 µM). For the determination of kinetic parameters, either the concentration of $NADP^+$ or sodium phosphite were fixed in the assay at a concentration of 0.25 mM or 1 mM, respectively and the concentrations of the second substrate varied from 0.02-1 mM for $Na_2HPO_3$ or 0.005-0.25 mM for $NADP^+$. All reactions were started with the addition of $NADP^+$ and performed in four replicates. Kinetic parameters were calculated using the Michaelis-Menten fitting function of Origin (version 2022b) (**Table 2**).

#### P450BM3m NADPH coupling efficiency

To determine the NADPH coupling efficiency, reactions were performed as described above with 0.4 µM P450BM3m, 1 mM indole or lauric acid and 1 mM NADPH in 0.1 M sodium phosphate buffer (pH 7.0) at 30 ºC. For indole, 200 µL reactions were followed spectrophotometrically at 340 nm and 670 nm until complete consumption of NADPH after 1.5 h. The coupling efficiency for indole was then calculated as the percentage of indoxyl (two molecules of indoxyl form one molecule of indigo) relative to the consumed NADPH. For lauric acid, 1 mL reactions were carried out for 1.5 h (30 ºC, 180 rpm) and reactions stopped by adding 10% (v/v) saturated NaCl $H_2SO_4$ (6 g NaCl in 10 mL 50% $H_2SO_4$). Lauric acid hydroxylation products were then identified and quantified following extraction and derivatization by GC-MS and GC-FID as described below. The coupling efficiency was calculated as the percentage of produced hydroxylauric acid products relative to the consumed NADPH. All assays were performed with four separate samples.

#### P450BM3m and PTDH cross-linking to EutM scaffolds

Cross-linking of P450BM3m and PTDH cargo proteins individually or combined to EutM scaffolds was confirmed by SDS-PAGE (**Fig. 7b and Supplementary Fig. 14)** as described for GFP cargo above except that the molar ratio of enzyme cargo to isopeptide bond forming EutM partner was increased to 1:4. Scaffolds and enzymes were mixed in 0.1 M sodium phosphate buffer (pH 7.0) at 30 ºC for 1 h before loading onto a gel. Individual enzyme immobilization reactions contained the following final concentrations: 15 µM His-SpyT-P450BM3m-SnoopT (1.9 mg/mL) or 15 µM His-SpyT-PTDH-SnoopT (0.7 mg/mL) mixed with 60 µM His-SpyC-EutM-SnoopC (2.0 mg/mL) or 60 µM SpyC-EutM-SnoopC in hybrid scaffolds His-EutM:SpyC-EutM-SnoopC=3.9:1 (4.7 mg/mL) or His-EutM:SpyC-EutM-SnoopC=8.8:1 (8.0 mg/mL). For co-immobilization, 15 µM His-SpyT-P450BM3m-SnoopT (1.9 mg/mL) or 15 µM His-SpyT-PTDH-SnoopT (0.7 mg/mL) (30 µM total enzyme cargo) were mixed with 120 µM SpyC-EutM-SnoopC in hybrid scaffold His-EutM:SpyC-EutM-SnoopC=8.8:1 (16 mg/mL). This co-immobilized sample was also analyzed with TEM as described above (**Fig. 8a and Supplementary Fig. 18**). Enzyme or scaffold only reactions served as controls for SDS-PAGE analysis.

#### Effect of scaffold immobilization on P450BM3m activity and stability

To assess the effect of scaffold attachment on P450BM3m activity (**Fig. 7c** and **Supplementary Fig. 15**), 20 µM P450BM3m (dual-modified and unmodified as control, 2.5 mg/mL) were mixed at a 1:4 molar ratio with 80 µM His-SpyC-EutM-SnoopC (2.7 mg/mL) in 0.1 M sodium phosphate buffer (pH 7.0). This mixture was then quickly diluted with the same buffer to achieve final enzyme concentrations ranging from (0.5, 1.0, 2.0, 8.0, 16.0, 20 µM) following by 1 h incubation at 30 °C and 180 rpm for cross-linking. A 40 µL reaction mixture was then taken out from each sample to quantify specific and volumetric P450BM3m activities in 200 µL assays as described above by monitoring both NADPH oxidation and indigo formation with 0.25 mM NADPH and 2.5 mM indole. The final P450BM3m concentrations in the assays (after 5 x dilution) ranged from 0.1-4 µM. Note that NADPH oxidation could not be quantified in assays containing 1.6 µM or higher enzyme concentrations as NADPH was completely consumed after 30 s.

To compare the effect of scaffold type on P450BM3m activity (**Fig. 7d**), 8 µM His-P450BM3m (1.0 mg/mL) or His-SpyT-P450BM3m-SnoopT (1.0 mg/mL) were mixed with 32 µM His-SpyC-EutM-SnoopC (1.1 mg/mL) or 32 µM SpyC-EutM-SnoopC in His-EutM:SpyC-EutM-SnoopC=3.9:1 (2.5 mg/mL) or His-EutM:SpyC-EutM-SnoopC=8.8:1 (4.3 mg/mL), incubated for 1 h at 30 °C and 180 rpm and the specific activity for indigo measured with 40 µL reaction mixture (final P450BM3m concentration 1.6 µM (0.2 mg/mL) in assay) as described above. To determine the effect of scaffolds on P450BM3m stability (**Supplementary Fig. 16**), the above reaction mixtures were incubated at 30 °C and 40 µL samples removed at 0-168 h for measurement of specific activity and indigo formation as described above. All measurements were performed with four replicates for each enzyme immobilization reaction. Control reactions contained no scaffolds.

#### Effect of scaffold immobilization on PTDH activity and stability

To determine the effect of hybrid scaffold on PTDH activity, 0.5 µM His-PTDH or His-SpyT-PTDH-SnoopT (0.02 mg/mL) were mixed at a molar ration of 1:4 with 2 µM His-SpyC-EutM-SnoopC (0.07 mg/mL) or 2 µM SpyC-EutM-SnoopC in His-EutM:SpyC-EutM-SnoopC=3.9:1 (0.2 mg/mL) or His-EutM:SpyC-EutM-SnoopC=8.8:1 (0.3 mg/mL) and incubated within 0.1 M sodium phosphate buffer (pH 7.0) at 30 °C and 180 rpm for 1 h. 40 µL reaction mixture was then assayed in a 200 µL assay to determine the specific PTDH activity as described above with 1 mM sodium phosphite and 0.25 mM $NADP^+$ in 0.1 M sodium phosphate buffer (pH 7.0) (**Fig. 7d**). The effect of scaffolds on PTDH stability were measured by incubating the reaction mixtures for up to 168 h at 30 °C and removing 40 µL samples after set time intervals for specific activity measurements. All measurements were performed with four replicates for each enzyme immobilization reaction. Control reactions contained no scaffolds (**Supplementary Fig. 17**).

**Implementation of coupled reaction system with free P450BM3m and PTDH**

To test and optimize the coupled P450BM3m-PTDH system for indole or lauric acid conversion, reactions were performed with fixed PTDH and ratios of P450BM3m for the conversion of 2.5 mM indole or lauric acid. (**Fig. 8b**). For indole conversion, indigo formation was quantified spectrophotometrically (see kinetic assay above) after 15 min at 30 °C (600 rpm) in 200 µL reactions containing 20 µL 100 mM sodium phosphite (final concentration 10 mM), 10 µL 5 mM $NADP^+$ (final concentration 0.25 mM), 2 µL 250 mM indole in DMSO (final concentration 2.5 mM), 40 µL enzyme mixture with 8 µM (0.3 mg/mL) His-PTDH and 2, 4, or 8 µM (0.2-1.0 mg/mL) His-P450BM3m (final concentration 1.6 µM His-PTDH; 0.4, 0.8, or 1.6 µM His-P450BM3m) in 0.1 M sodium phosphate buffer (pH 7.0). For lauric acid conversion, hydroxylauric acid formation was quantified by GC-FID (see GC analysis below) after 15 mins at 30 °C (180 rpm) in 1 mL reactions containing 100 µL of 100 mM sodium phosphite (final concentration 10 mM), 50 µL of 5 mM $NADP^+$ (final concentration 0.25 mM), 80 µL of 31.25 mM lauric acid in DMSO (final concentration 2.5 mM), 200 µL of enzyme mixture with 8 µM His-PTDH and 2, 4, or 8 µM His-P450BM3m (final concentration 1.6 µM His-PTDH; 0.4, 0.8, or 1.6 µM His-P450BM3m) in 0.1 M sodium phosphate buffer (pH 7.0). Higher enzyme concentrations were also tested for hydroxylauric acid formation by conducting reactions under the same conditions with equimolar concentrations (1.6, 3.2, 6.4, 12.8 or 25.6 µM) of P450BM3m and PTDH (**Supplemental Fig. 20**). All assays were performed with four separate samples.

**Influence of P450BM3m and NADPH concentrations on indole oxidation**

To characterize inhibition of indigo formation by high P450BM3m and NADPH concentration (**Supplementary Fig. 19**), P450BM3m spectrophotometric assays were performed (see above) with 0.25 mM or 3.5 mM NADPH and with either 0.4 µM or 1.6 µM (0.05 or 0.2 mg/mL) His-P450BM3m or His-SpyT-P450BM3m-SnoopT. Indigo formation was monitored and quantified at 670 nm after 20 min at 30 °C. Assays were performed with four separate samples.

**Scaffolded P450BM3m-PTDH reactions system for lauric acid conversion**

Small scale reactions (1 mL) with both enzymes were set up with and without (control) scaffolds to characterize effects on conversion (**Fig. 8c**). For this, 6.4 µM His-SpyT-P450BM3m-SnoopT (0.8 mg/mL) and 6.4 µM His-SpyT-PTDH-SnoopT (0.3 mg/mL) were mixed at a 1:4 molar ratio of enzymes to cross-linking scaffold building blocks with 51.2 µM His-SpyC-EutM-SnoopC (1.8 mg/mL) or 51.2 µM SpyC-EutM-SnoopC in His-EutM:SpyC-EutM-SnoopC=3.9:1 (4.0 mg/mL) or in His-EutM:SpyC-EutM-SnoopC=8.8:1 (6.9 mg/mL) hybrid scaffolds. The mixtures were then incubated at 30 °C and 180 rpm for 1 h to allow for cross-link formation. Conversions were then performed in 1 mL reactions by mixing 500 µL of the scaffolded enzymes (or enzymes only control) with 270 µL 0.1 M sodium phosphate buffer (pH 7.0), 100 µL 100 mM sodium phosphite (final concentration 10 mM), 80 µL 31.25 mM lauric acid in DMSO (final concentration 2.5 mM) and 50 µL 5 mM $NADP^+$ (final concentration 0.25 mM), to start the reactions. The final P450BM3m and PTDH concentrations were 3.2 µM. After 10 min incubation at 30°C and 180 rpm, reactions were stopped (see assay for coupling efficiency), and lauric acid hydroxylation products extracted and quantified (see below). All reactions were performed with four separate samples.

For larger scale conversion reactions (5 mL) with 20% (v/v) dodecane (**Fig. 8d**) followed over 24 h, 1.6 µM His-SpyT-P450BM3m-SnoopT (0.2 mg/mL) and 6.4 µM His-SpyT-PTDH-SnoopT (0.3 mg/mL) were first mixed with 20 µM hybrid His-EutM:SpyC-EutM-SnoopC=8.8:1 (2.7 mg/mL) (or without scaffolds as control) and incubated at 30°C and 180 rpm for 1 h to allow for the co-immobilization of enzymes and cross-linking of scaffolds. Conversion reactions were then performed in 5 mL reactions by mixing 2.5 mL of the scaffolded enzymes (or enzymes only control) with 1.4 mL of 0.1 M sodium phosphate buffer (pH 7.0), 0.5 mL of 0.5 M sodium phosphite (final concentration 50 mM), 0.1 mL DMSO (final concentration 2% (v/v)), and 1 mL of 100 mM lauric acid in dodecane (final lauric acid concentration 20 mM and 20% (v/v) dodecane). Reactions were started with 0.5 mL of 5 mM $NADP^+$ (final concentration 0.5 mM). The final enzyme concentrations in the reactions were 0.8 µM His-SpyT-P450BM3m-SnoopT and 3.2 µM His-SpyT-PTDH-SnoopT. After 0.5, 1, 3, 6, 9, 12, and 24 h incubation at 30°C and 120 rpm, 100 µL and 20 µL aliquots were removed from the aqueous and dodecane phases, respectively, and combined. Reactions were stopped (see assay for coupling efficiency) for the quantification of lauric acid conversion products (see below). All reactions were performed with four separate samples.

**Recycling of scaffold co-immobilized P450BM3m and PTDH for lauric acid conversion**

Dual-modified P450BM3m and PTDH (6.4 µM each) were first co-immobilized onto hybrid EutM:His-SpyC-EutM-SnoopC scaffolds at a 1:4 molar ratio of enzymes to cross-linking scaffold building block as described above. For the first reaction cycle, conversions were performed in 3 mL reactions by mixing 1.5 mL of the scaffolded enzymes with 0.81 mL of 0.1 M sodium phosphate buffer (pH 7.0), 0.3 mL of 0.1 M sodium phosphite (final concentration 10 mM), and 0.24 mL of 31.25 mM lauric acid in DMSO (final lauric acid concentration 2.5 mM and 8% (v/v) DMSO). Reactions were started with 0.15 mL of 5 mM $NADP^+$ (final concentration 0.25 mM). The final enzyme concentrations in the reactions were 3.2 µM His-SpyT-P450BM3m-SnoopT and 3.2 µM His-SpyT-PTDH-SnoopT. After 30 min incubation at 30°C and 180 rpm, a 100 µL aliquot was removed for the quantification of lauric acid conversion products (see below). Another 100 µL was removed to measure protein concentration for SDS-PAGE analysis (**Supplemental Fig. S21**). The remaining sample was spun down at 5000 x*g* for 5 min at 4 ºC to recover the scaffolded enzymes. After removing the supernatant, the material was reused under the same conditions and in the same volume for the next reaction cycle. All reactions were performed with four separate samples.

**Gas chromatography (GC) analysis of lauric acid hydroxylation**

Stopped enzyme reaction samples were extracted twice with an equal volume of ethyl acetate. The organic extracts were collected and 10 µL of 10 mM of palmitic acid (corresponding to 1 mM after derivatization) was added as internal reference prior to evaporation and resuspension in 50 µL dimethylformamide (DMF). Resuspended samples were then derivatized for GC analysis with 50 µL BSTFA with 1% TMCS (N, O-bis(trimethylsilyl)trifluoroacetamide with 1% trimethylchlorosilane) (MilliporeSigma, Burlington, MA, USA) at 60°C for 30 min. For quantification by GC analysis of hydroxylauric acid products and the lauric acid substrate, derivatization reactions were similarly performed with 1 mM palmitic acid (as internal reference) and 0.01-2 mM 12-hydroxylauric acid or 0.01-1 mM lauric acid in 50 µL DMF to calculate detector response factors for derivatized monohydroxylauric acids and lauric acid[11]. Peak areas of all hydroxylated lauric acid products were combined to quantify hydroxylation activity of P450BM3m.

Derivatized samples were then analyzed and quantified using an Agilent 6890 Plus gas chromatograph with a flame ionization detector and equipped with a capillary column (HP-5ms, 30 m x 0.25 mm x 0.25 µm, Agilent, Santa Clara, CA, USA) with the following parameters: The FID heater temperature was set to 320 ºC and flow rates for $H_2$, air, and make up gas (helium) were 40, 450, and 45 mL/min, respectively. A representative GC-FID chromatogram with retention times of derivatized fatty acids is shown in **Supplementary Fig. 22**.

To identify derivatized lauric acid oxidation products of samples, GC mass spectrometry (GC-MS) was performed with an Agilent 7890A GC system and a 5975C MSD detector equipped with a capillary column (HP-5ms, 30 m x 0.25 mm x 0.25 µm, Agilent, Santa Clara, CA, USA) with the following parameters: 1 µL injection volume with a 300 °C port temperature and a split ratio of 10; helium as carrier gas and a temperature gradient from 40 to 300 °C with 5 °C/min and a 10 min isothermal hold at 300 °C and a 8 min solvent delay. Representative fragmentation spectra of derivatized 9-, 10- and 11-hydroxylauric acids are shown in **Supplementary Fig. 23** and matched reported spectra[11,13] and expected sequence of retention times.

### Statistical Analysis and Reproducibility

Enzyme kinetic parameters were calculated using the Hill or Michaelis-Menten fitting function in Origin (version 2022b). R-squared values were > 0.95. Standard curves for hydroxylauric acid (R-squared values ≥ 0.99) and EutM protein densitometry (R-squared value > 0.9) analysis were created in Microsoft Excel 365 using the linear trendline function. Mean and standard deviations were calculated using Average and STDEV functions in Microsoft Excel 365. The yield, molar ratios, insoluble and soluble fractions of hybrid scaffolds were measured in triplicate with protein isolated from three independent recombinant cultures per scaffold expression construct. All scaffold proteins and enzymes were purified at least three times to isolate sufficient quantities for all experiments. Cross-linking experiments for PAGE analysis and microscopy were done at least twice to obtain representative gel and microscopy images. All enzyme reactions were done as four separate replicate samples.

### Supplementary Figures

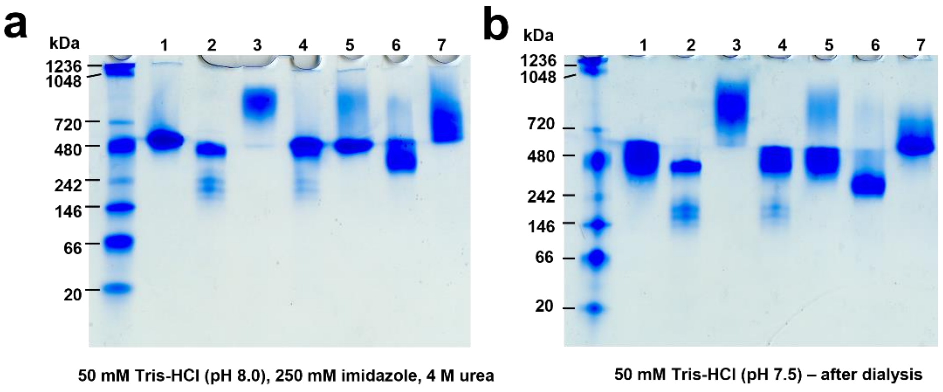

**c** **Molecular weights of EutM monomers and of possible hexamer assemblies of scaffolds with sizes observed in (a) & (b)**

| Lane | Scaffolds | Monomers | | Hexamers | | Hexamer Multimers | |
|---|---|---|---|---|---|---|---|
| | | | kDa | | kDa | # | kDa |
| 1 | His-EutM | His-EutM | 11.5 | 6x His-EutM | 69 | 6x | 414 |
| 2 | His-SpyC-EutM-SnoopC | His-SpyC-EutM-SnoopC | 34.3 | 6x His-SpyC-EutM-SnoopC | 205.8 | 2x | 411.6 |
| 3 | His-SnoopT-EutM-SpyT | His-SnoopT-EutM-SpyT | 15.8 | 6x His-SnoopT-EutM-SpyT | 94.8 | 8x | 758.4 |
| 4 | Purified proteins mixed His-EutM:His-SpyC-EutM-SnoopC 5:1 | His-EutM | 11.5 | 6x His-EutM | 69 | 6x | 414 |
| | | His-SpyC-EutM-SnoopC | 34.3 | 6x His-SpyC-EutM-SnoopC | 205.8 | 2x | 411.6 |
| 5 | Purified proteins mixed His-EutM:His-SnoopT-EutM-SpyT 5:1 | His-EutM | 11.5 | 6x His-EutM | 69 | 6x | 414 |
| | | His-SnoopT-EutM-SpyT | 15.8 | 6x His-SnoopT-EutM-SpyT | 94.8 | 8x | 758.4 |
| 6 | Co-expressed hybrid scaffold His-EutM:SpyC-EutM-SnoopC= 3.9:1 | His-EutM | 11.5 | His-EutM:SpyC-EutM-SnoopC (5x:1x) | 90.2 | 5x | 451 |
| | | SpyC-EutM-SnoopC | 32.7 | His-EutM:SpyC-EutM-SnoopC (4x:2x) | 166.2 | 2x | 322.4 |
| 7 | Co-expressed hybrid scaffold His-EutM:SnoopT-EutM-SpyT= 3.3:1 | His-EutM | 11.5 | His-EutM:SnoopT-EutM-SpyT (5x:1x) | 72 | 8x | 576 |
| | | SnoopT-EutM-SpyT | 14.5 | His-EutM:SnoopT-EutM-SpyT (4x:2x) | 75 | 8x | 600 |

**Supplementary Fig. 1. Native PAGE analysis of EutM assemblies.**

Purified EutM scaffolds normalized to 2 mg/mL in elution buffer with 4 M urea after metal affinity chromatography (**a**) and after dialysis into buffer without urea (**b**) were analyzed by native PAGE to determine the molecular weights of soluble assemblies able to migrate into the gel. Proteins loaded onto the gel are listed in (**c**) along with the molecular weights of their monomers and possible hexameric assemblies corresponding to sizes of gel separated protein bands.

Assembly sizes before and after dialysis are similar except for assemblies of the hybrid His-EutM:SpyC-EutM-SnoopC=3.9:1 scaffold (purified from *E. coli*) which appear smaller after dialysis (lane 6). Purified His-EutM mixed as a control with at a molar ratio of 5:1 with either His-SpyC-EutM-SnoopC (lane 4) or His-SnoopT-EutM-SpyT (lane 5) formed assemblies with sizes like His-EutM (lane 1). Assemblies with larger sizes are formed by His-SnoopT-EutM-SpyT (lane 3) and hybrid His-EutM:SnoopT-EutM-SpyT=3.3:1 (lane 7), while the presence of His-SpyC-EutM-SnoopC (lane 2 & 4) leads to smaller assemblies. Results shown are from one set of purified proteins.

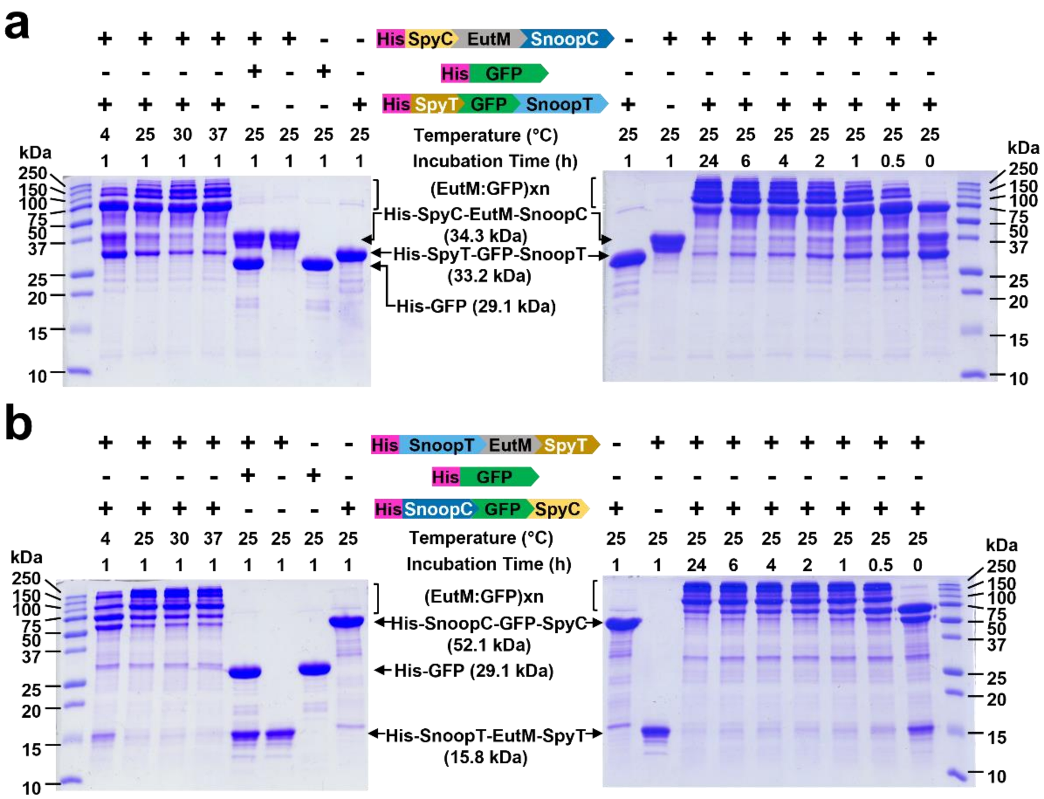

**Supplementary Fig. 2. Characterization of scaffold building block cross-linking.**

SDS-PAGE analysis of EutM building block crosslinking with GFP. Purified His-SpyC-EutM-SnoopC (50 µM) (**a**) or His-SnoopT-EutM-SpyT (50 µM) (**b**) was mixed at 1:1 molar ratio with GFP (50 µM) with and without (control) cognate Catcher/Tag fusions. Proteins were incubated in 0.1 M sodium phosphate buffer (pH 7.0) for 0-24 h at 4, 25, 30, or 37 ºC prior to analysis (see Methods for details). Higher molecular weight (EutM:GFP)xn only form with dual-modified GFP. Isopeptide bond formation is fast with the majority of EutM cross-linked to GFP after 1 h at 25 ºC. Shown data are representative for one set of purified proteins.

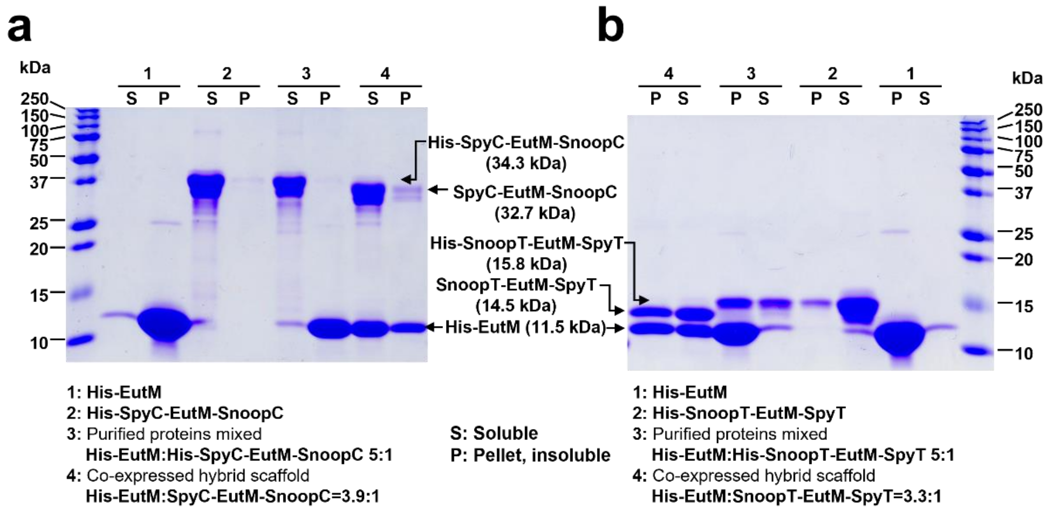

**Supplementary Fig. 3. SDS-PAGE analysis of EutM scaffold co-assembly.**

Purified single protein and EutM hybrid (co-expressed in *E. coli*) His-SpyC-EutM-SnoopC=3.9:1 (**a**) and His-SnoopT-EutM-SpyT=3.3:1 scaffold (**b**) in elution buffer with 4 M urea were normalized to 2 mg/mL and then dialyzed into 50 mM Tris-HCl buffer at pH 7.5. Insoluble (Pellet) and soluble (S) scaffold fractions were separated and analyzed to compare their compositions (see Methods for details). Purified EutM scaffolds and mixtures (at 5:1 molar ratios) of purified EutM and of double-modified EutM treated in the same way were included as controls. Representative data from one set of purified proteins are shown.

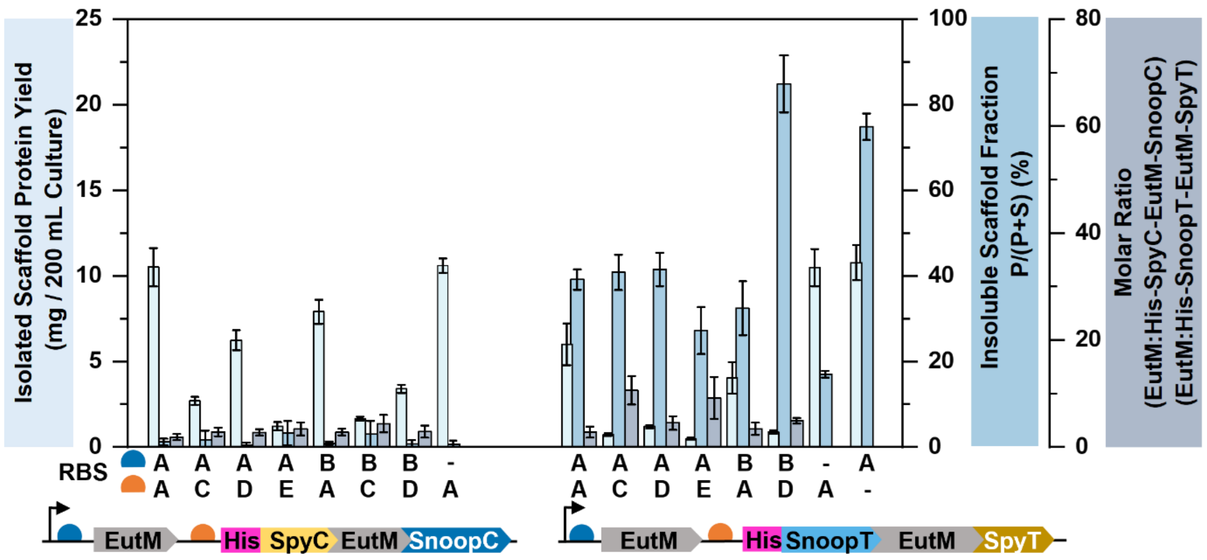

**Supplementary Fig. 4. Characterization of hybrid scaffolds co-assembled from EutM and dual-modified His-EutM.**

Eight RBS combinations (RBS A-E) were used to vary scaffold building block ratios for co-expression in *E. coli*. Scaffolds were purified from 200 mL cultures in the presence of urea and the total yield of isolated scaffolds was quantified. Hybrid scaffolds can be selectively pulled-down by His-tag affinity purification, yielding purified scaffolds that separate into a soluble fraction (S) composed of smaller scaffolds and an insoluble fraction (pellet P). The percentage of the insoluble scaffold fraction of normalized scaffolds (2 mg/mL) (as an indicator of assembly strength) was determined after dialysis into 0.1 M Tris-HCl buffer (pH 7.5) to remove urea. The molar ratios of EutM building blocks for the isolated hybrid scaffolds were analyzed by SDS-PAGE densitometry. Data are shown as mean values ± SD and error bars represent the standard deviations of replicates from three independent cultures for each expression construct. **Supplementary Fig. 5** shows representative SDS-PAGE gels used for analysis.

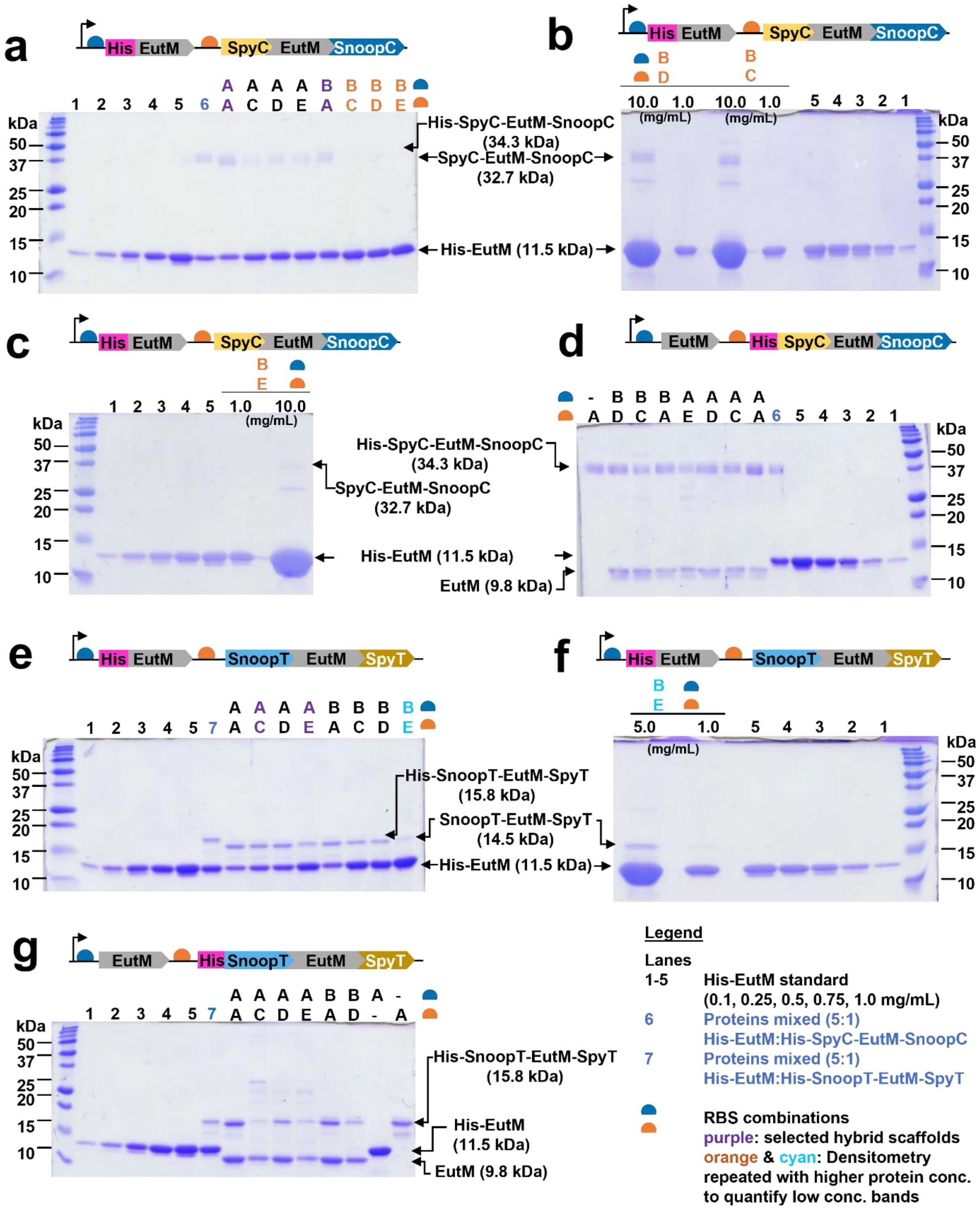

Figure legend next page.

**Supplementary Fig. 5. SDS-PAGE analysis of EutM hybrid scaffolds for protein quantification.**

(**a, b, c**) Representative SDS-PAGE gels of hybrid His-EutM:SpyC-EutM-SnoopC hybrid scaffolds co-expressed with different RBS combinations. RBS combinations B/C, B/D and B/E resulted in scaffolds with too faint bands for Spyc-EutM-SnoopC and gels were rerun (**b, c**) with higher hybrid-scaffold protein concentrations for quantification by densitometry. (**d, e**) Representative gels for hybrid EutM:His-SpyC-EutM-SnoopC and EutM:His-SnoopT-EutM-SpyT. (**f**) Rerun of hybrid EutM:His-SnoopT-EutM-SpyT scaffolds with RBS combinations B/E at higher concentrations. (**g**) Representative gel for EutM:His-SnoopT-EutM-SpyT constructs. Lanes 1-5 include His-EutM standards with known protein concentrations for densitometry quantification. Lanes 6 & 7 contain mixed EutM building blocks with fixed molar ratios for comparison. SDS-PAGE analysis was repeated nine times with three sets of purified proteins from three independent cultures to quantify the molar ratio between EutM and dual-modified EutM as presented in **Fig. 3b** and **Supplementary Fig. 4**. Representative images are shown.

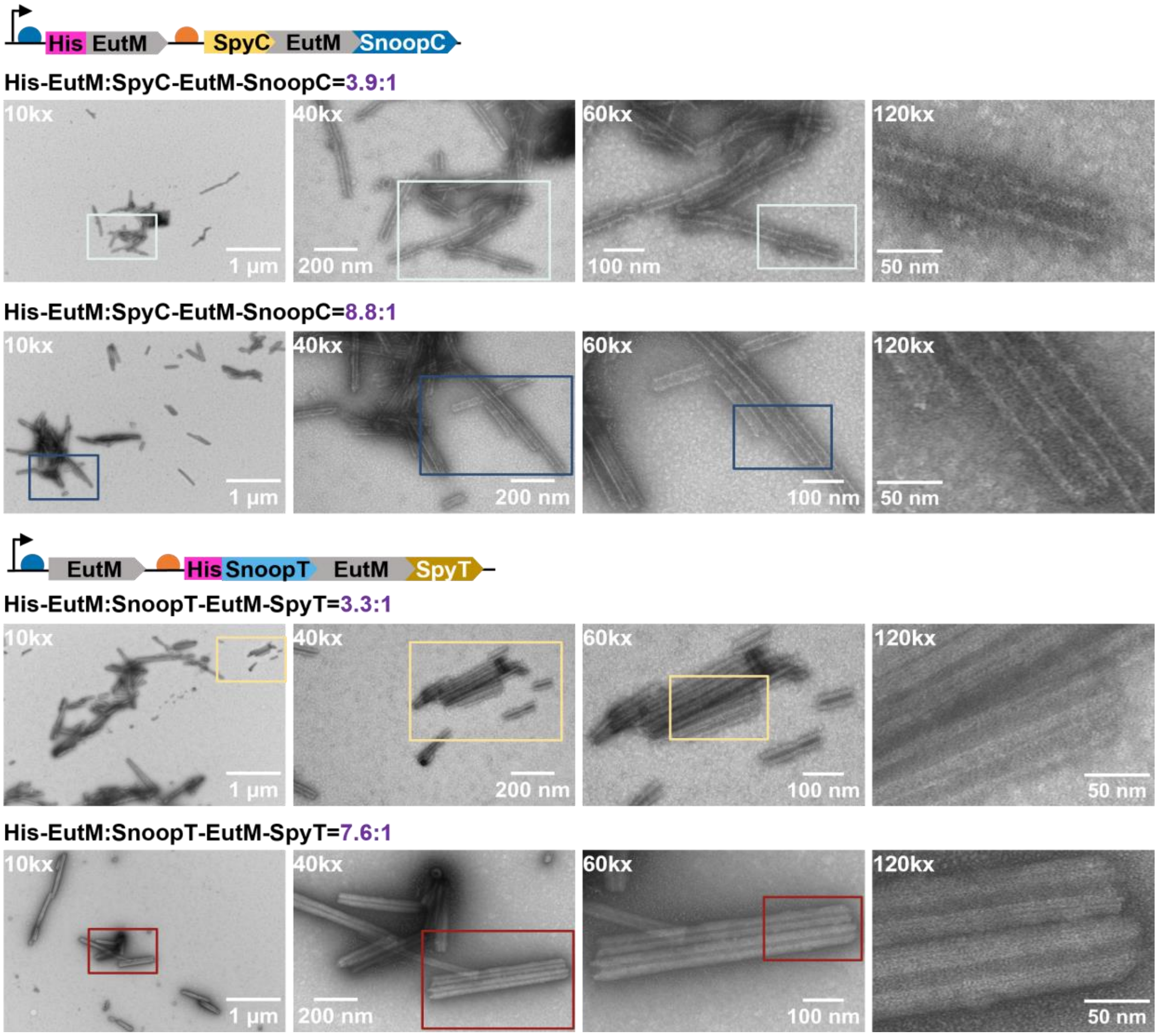

**Supplementary Fig. 6. TEM of selected hybrid EutM scaffolds.**

Purified hybrid scaffolds after dialysis into 0.1 M sodium phosphate buffer (pH 7.0) and dilution to 1 mg/mL were applied to grids for TEM. Shown are representative images from one set of purified proteins captured for one region (colored boxes) of each scaffold at different magnifications from 10,000x to 120,000x. Additional images at 60,000x for each scaffold are shown in **Fig. 3c**.

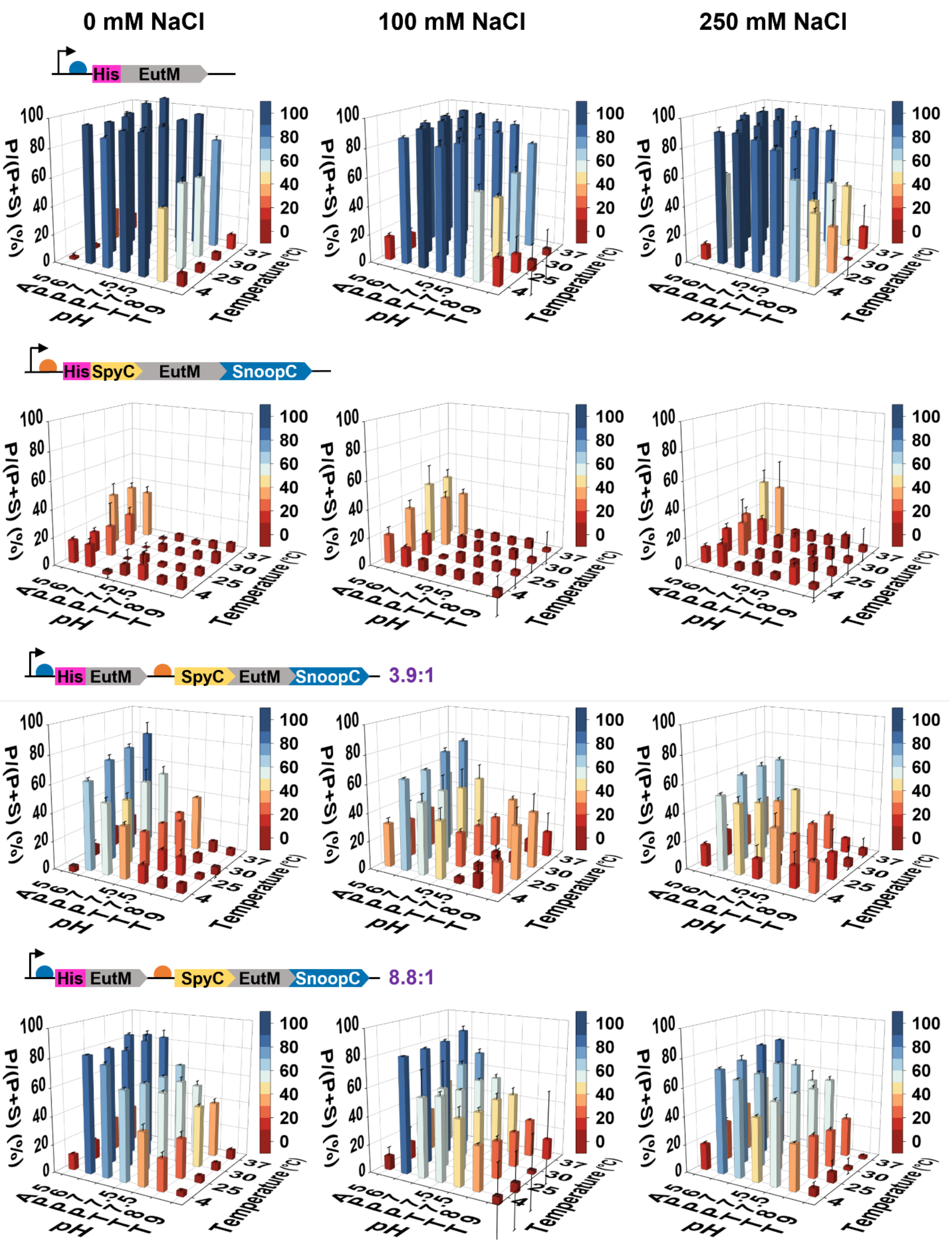

**Figure legend next page.**

**Supplementary Fig. 7. Characterization of His-EutM:SpyC-EutM-SnoopC hybrid scaffold assembly strength.**

Hybrid His-EutM:SpyC-EutM-SnoopC scaffolds with two different molar ratios (purple numbers) of co-expressed building blocks were characterized for assembly strength without and with 100 mM or 250 mM NaCl added and compared to scaffolds formed by individual building blocks. Scaffolds (3 mg/mL in urea) were dialyzed into buffers with different pH values, normalized to 2 mg/mL and then incubated for 24 h at different temperatures. The percentage of the insoluble scaffold (P) fraction (P/(P + S) as in **Fig. 3**) was then calculated as indicator of assembly strength. The following buffers were used: 0.1 M sodium acetate (A, pH 5), 0.1 M sodium phosphate (P, pH 6, 7, or 7.5), and 0.1 M Tris-HCl (T, pH 7.5, 8 or 9). Data are shown as mean values ± SD and error bars represent the standard deviations of three independent experiments per scaffold.

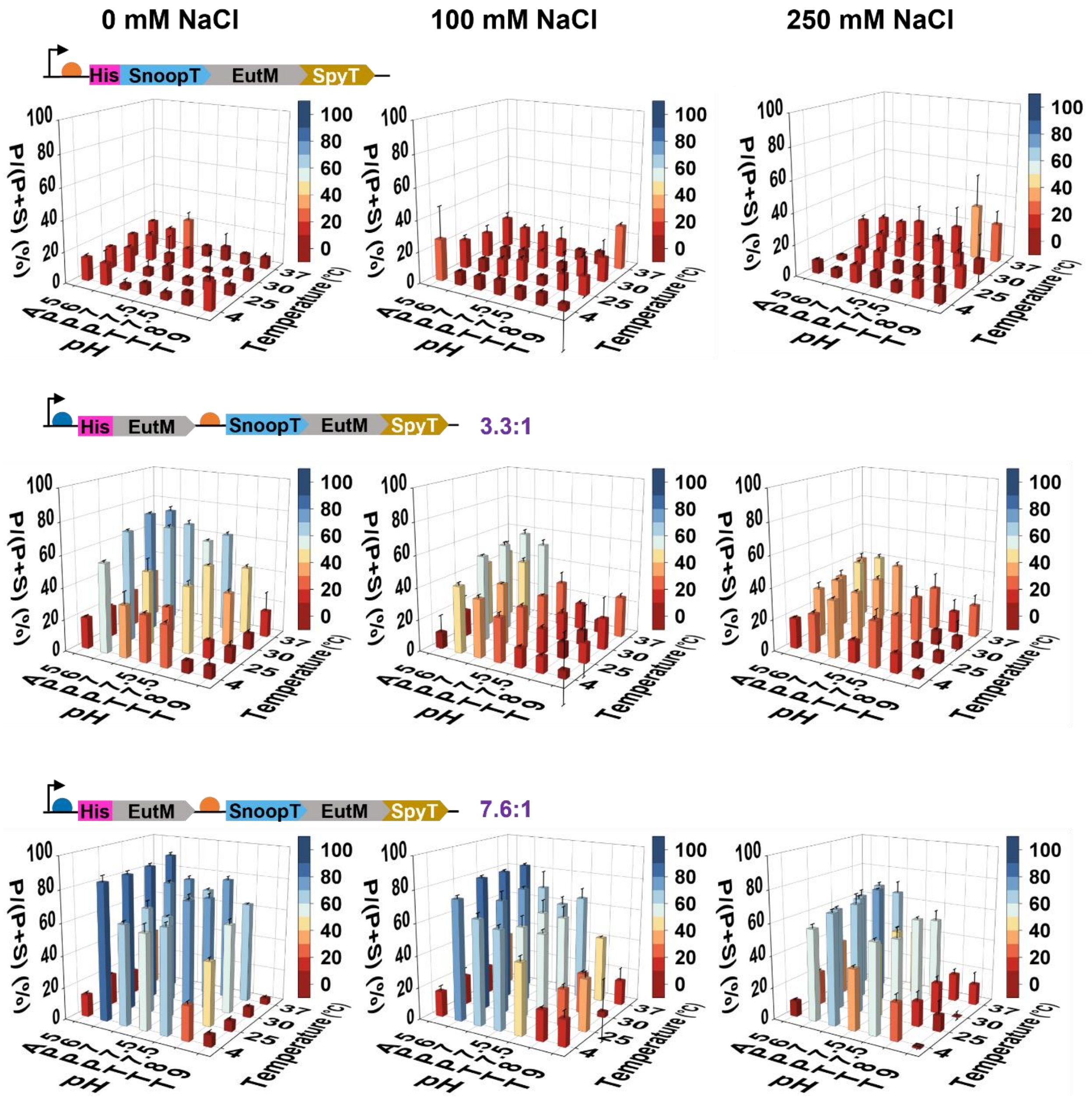

**Supplementary Fig. 8. Characterization of His-EutM:SnoopT-EutM-SpyT hybrid scaffold assembly strength.**

Hybrid His-EutM:SnoopT-EutM-SpyT scaffolds with two different molar ratios (purple numbers) of co-expressed building blocks were characterized for assembly strength without or with 100 mM or 250 mM NaCl added and compared to scaffolds formed His-SnoopT-EutM-SpyT (His-EutM, see **Supplementary Fig. 7**). Scaffolds (3 mg/mL in 4 M urea) were dialyzed into buffers with different pH values, normalized to 2 mg/mL and then incubated for 24 h at different temperatures. The percentage of the insoluble scaffold (P) fraction (P/(P + S) as in **Fig. 3**) was then calculated as indicator of assembly strength. The following buffers were used: 0.1 M sodium acetate (A, pH 5), 0.1 M sodium phosphate (P, pH 6, 7, or 7.5), and 0.1 M Tris-HCl (T, pH 7.5, 8 or 9). Data are shown as mean values ± SD and error bars represent the standard deviations of three independent experiments per scaffold.

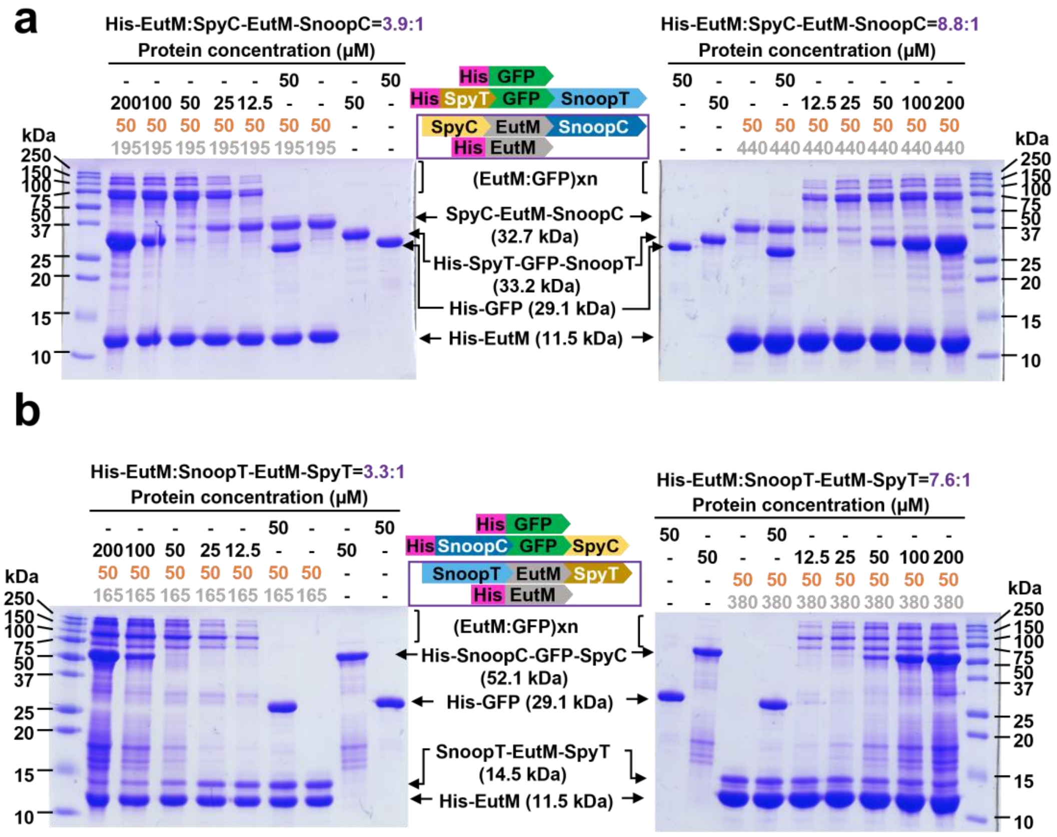

**Supplementary Fig. 9. Characterization of hybrid scaffold building block cross-linking.**

Cross-linking of EutM building blocks (purple boxes) in the four selected hybrid scaffolds (purple numbers) with GFP were analyzed by SDS-PAGE. For this purified hybrid scaffolds at concentrations corresponding to 50 µM (orange number, grey number His-EutM concentrations based on densitometry calculated scaffold building block molar ratios) of their dual modified SpyC-EutM-SnoopC (**a**) or SnoopT-EutM-SpyT (**b**) was mixed at different molar ratios with GFP (12.5-200 µM) with or without (control) cognate Catcher/Tag fusions (see Methods for details). Proteins were incubated in 0.1 M sodium phosphate buffer (pH 7.0) for 1 h at 25 ºC prior to analysis. Higher molecular weight (EutM:GFP)xn only form with dual-modified GFP. Isopeptide bond formation is fast with almost all GFP cargo cross-linked at a 50 µM concentration (1:1 ratio of cognate partner proteins) for the scaffolds with higher dual-tagged EutM building blocks at (left gels). For scaffolds with lower dual-tagged EutM building block ratios (right gels), crosslinking of the majority of GFP is achieved at a concentration of 25 µM. Shown data are representative for one set of purified proteins.

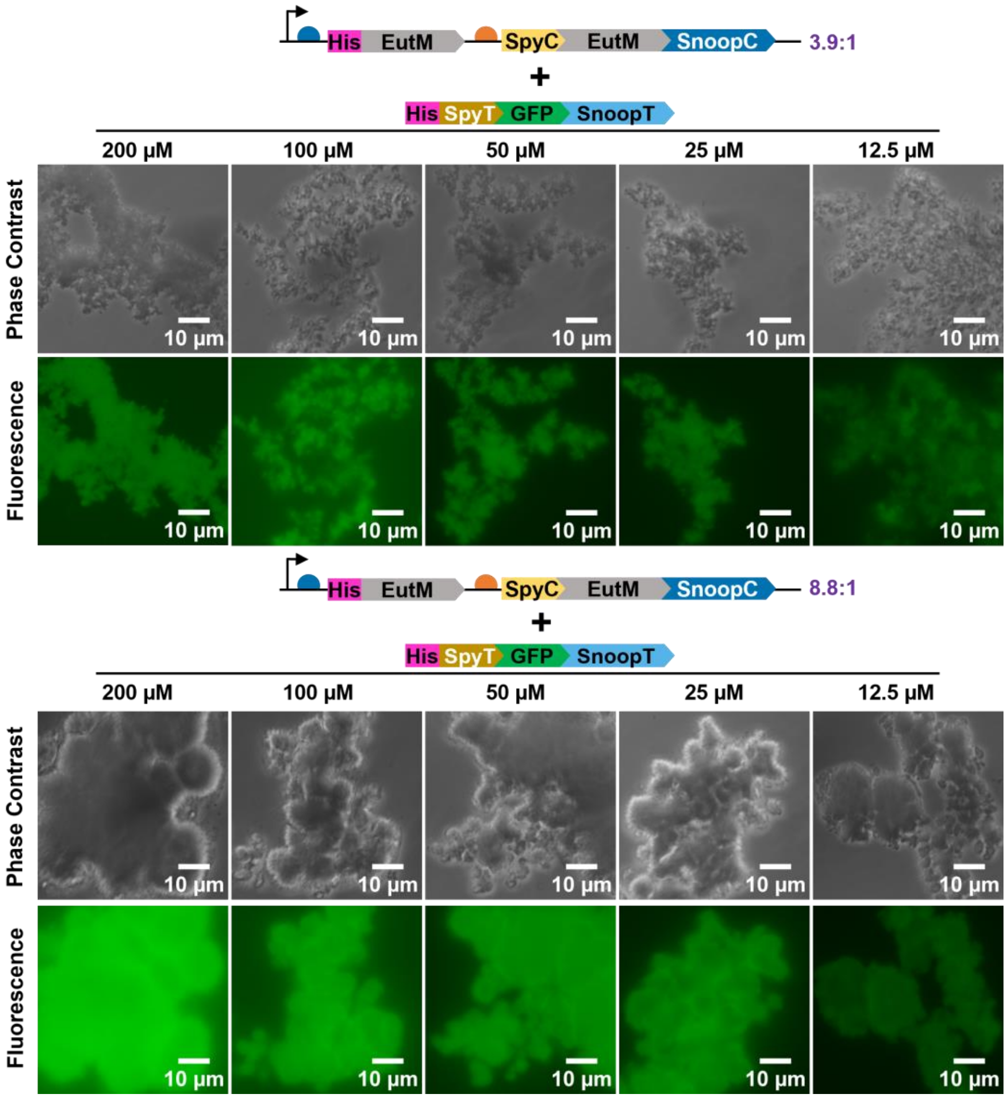

**Supplementary Fig. 10. Imaging of GFP attachment to hybrid His-EutM:SpyC-EutM-SnoopC scaffolds.**

His-EutM:SpyC-EutM-SnoopC hybrid scaffolds (molar ratios of 3.9:1 and 8.8:1 from **Fig. 3**) were mixed with His-SpyT-GFP-SnoopT at different molar ratios. Experiments were performed by mixing 12.5-200 µM GFP cargo in 0.1 M sodium phosphate buffer (pH 7.0) with hybrid scaffolds such that concentration of their modified EutM building blocks is fixed at 50 µM, resulting in molar ratios of the two isopeptide forming partner proteins from 4:1 to 1:4 (GFP:EutM) (see Methods for details). After incubation for 1 h at 25 °C and 180 rpm, samples were prepared for imaging. More dense macroscale structures are observed in hybrid scaffolds with a higher ratio of unmodified EutM (His-EutM:SpyC-EutM-SnoopC=8.8:1) and a higher concentration of GFP cargo-crosslinker appears to result in the formation of denser macroscale materials in the case of the His-EutM:SpyC-EutM-SnoopC=3.9:1 hybrid scaffolds. Representative images from one set of purified proteins are shown.

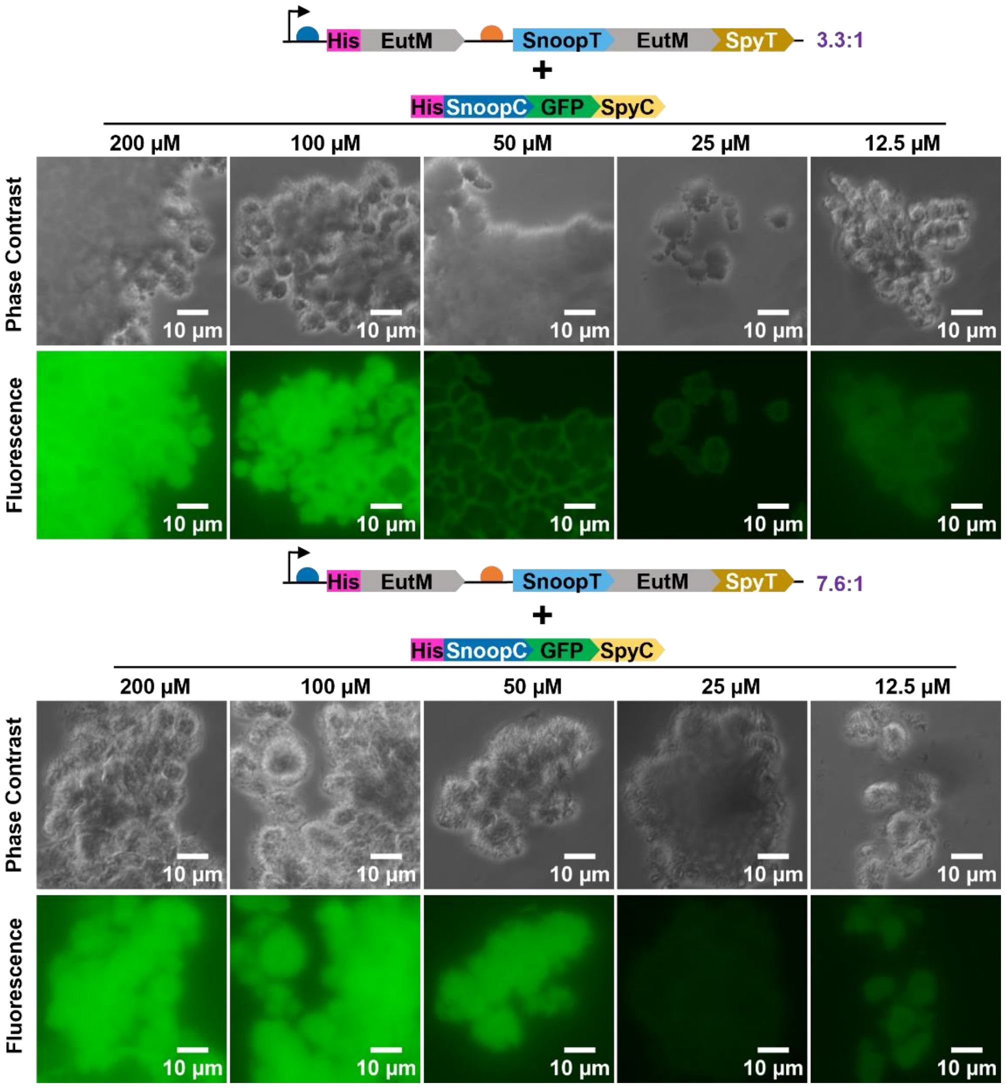

**Supplementary Fig. 11. Imaging of GFP attachment to hybrid His-EutM:SnoopT-EutM-SpyT scaffolds.**

His-EutM:SnoopT-EutM-SpyT hybrid scaffolds (molar ratios of 3.3:1 and 7.6:1 from **Fig. 3**) were mixed with His-SnoopC-GFP-SpyC at different molar ratios. Experiments were performed by mixing 12.5-200 µM GFP cargo in 0.1 M sodium phosphate buffer (pH 7.0) with hybrid scaffolds such that concentration of their modified EutM building blocks is fixed at 50 µM, resulting in molar ratios of the two isopeptide forming partner proteins from 4:1 to 1:4 (GFP:EutM) (see Methods for details). After incubation for 1 h at 25 °C and 180 rpm, samples were prepared for imaging. Both types of hybrid scaffolds form dense macroscale materials with attached GFP. Representative images from one set of purified proteins are shown.

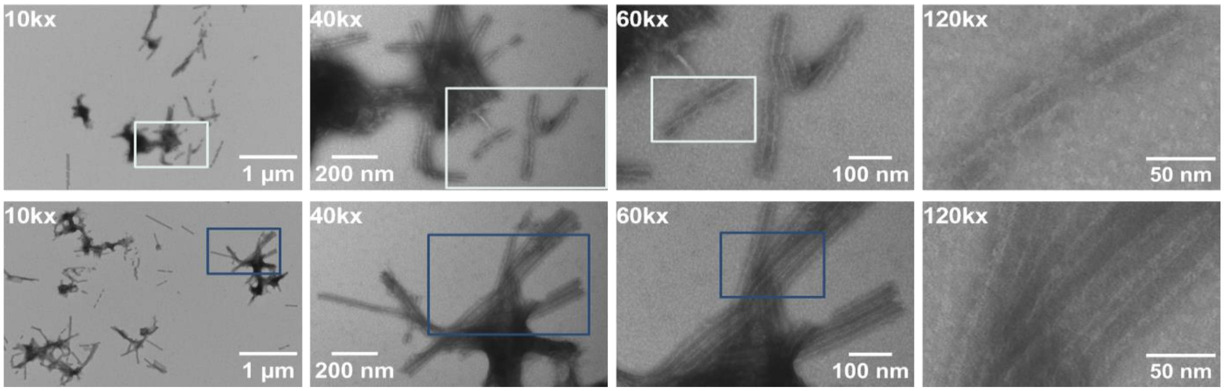

**Supplementary Fig. 12. TEM of GFP cross-linked hybrid His-EutM:SpyC-EutM-SnoopC scaffolds.**

Hybrid His-EutM:SpyC-EutM-SnoopC=8.8:1 scaffolds cross-linked with His-SpyT-GFP-SnoopT from **Fig. 5a** and **Fig. 6a** were diluted to 1 mg/mL and applied to grids for TEM imaging. Two grid positions are shown (top and bottom rows) with images captured at a range of magnifications (10k-120kx) for each location. Boxes indicate magnified regions for each of the two grid positions. The lowest magnification shows cross-linked and coated tubular structures. Individual tubes (top row) can be observed that are less articulated and thicker than those from scaffolds without attached GFP cargo (compare to **Fig. 3c** and **Supplementary Fig. 6**). Other tubes appear to be enveloped by a film and cross-linked together (lower panel). Representative images captured from one set of purified proteins are shown.

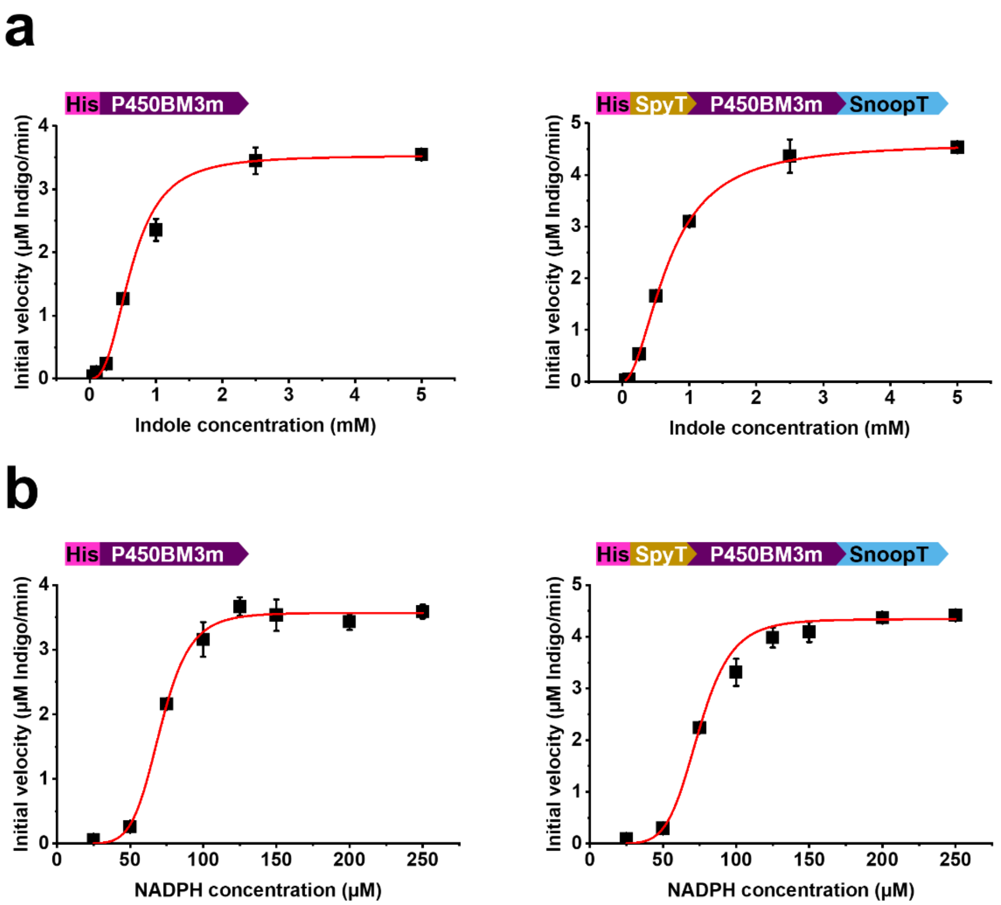

**Supplementary Fig. 13. Fitting curves for determining P450BM3m kinetic parameters.**

Enzyme kinetic parameters for unmodified and dual-modified His-P450BM3m (**Table 1**) were measured by varying the concentration of indole (**a**) or NADPH (**b**) as substrate and quantifying spectrophotometrically indigo formation. Assays were performed at 30 ºC in sodium phosphate buffer (pH 7.0) with 0.4 μM P450BM3m (see Methods for details). Initial velocities were measured, and curve fitting performed using the Hill function of Origin 2022b. Data are shown as means ± standard error and error bars represent the standard error of four independent replicates with one set of purified proteins.

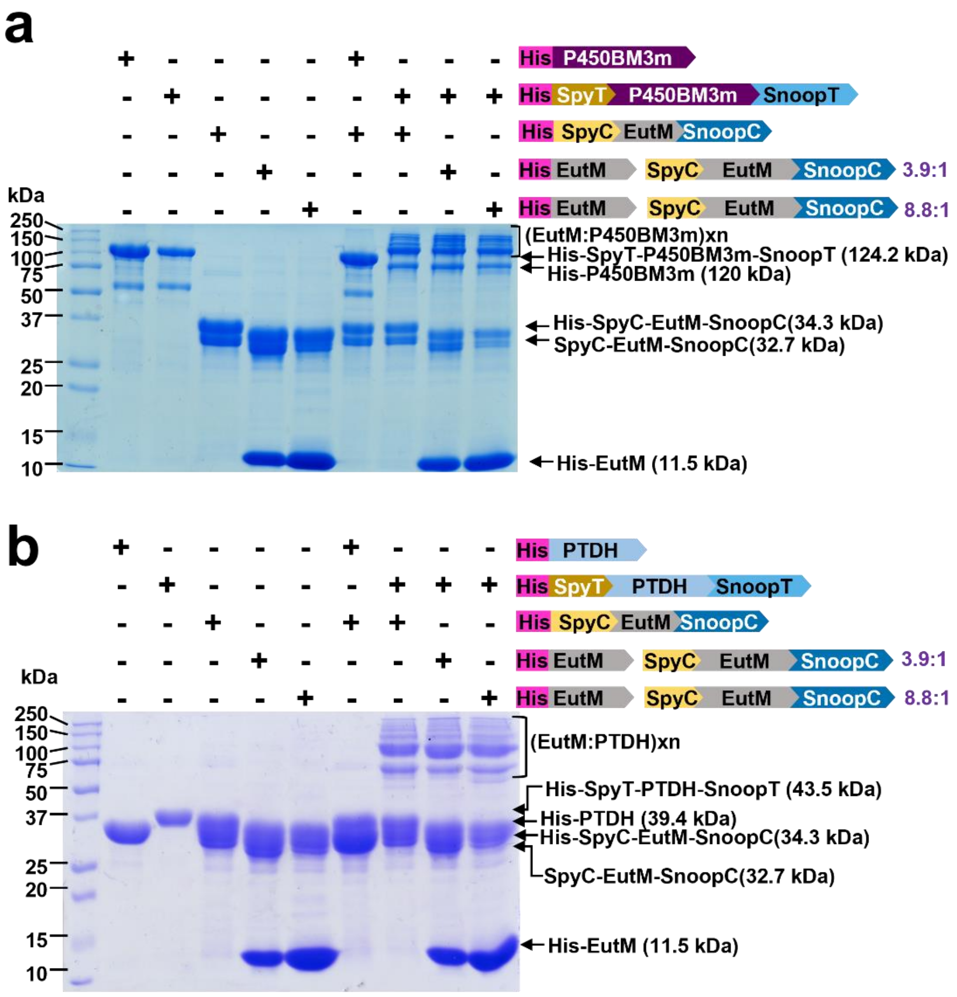

**Supplementary Fig. 14. Characterization of P450BM3m and PTDH cross-linking to hybrid scaffolds.**

Cross-linking of dual-modified or un-modified (control) His-P450BM3m and His-PTDH to the two selected hybrid His-EutM:SpyC-EutM-SnoopC scaffolds (3.9:1, 8.8:1) or to His-SpyC-EutM-SnoopC scaffolds (control) was analyzed by SDS-PAGE. Enzymes were mixed at a 1:4 molar ratio with their cognate binding partner in the scaffolds (see Methods for protein concentrations). Both enzymes are efficiently cross-linked to scaffolds, resulting in the formation of higher molecular weight complexes. Shown data are representative for one set of purified proteins.

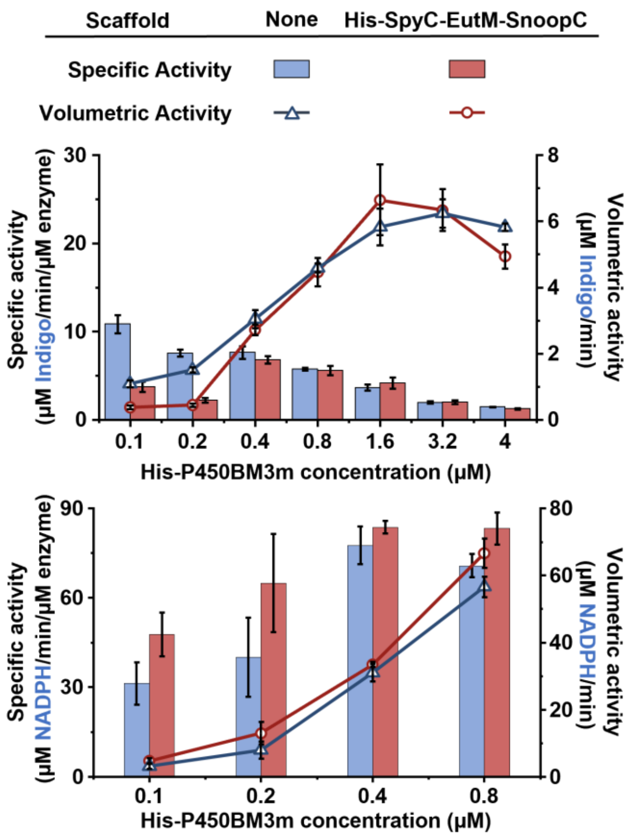

**Supplementary Fig. 15. Effect of scaffolds on unmodified His-P450BM3m activity.**

To determine the compatibility of scaffolds with P450BM3m, activity was determined by first mixing the enzyme with His-SpyC-EutM-SnoopC at a 1:4 molar ratio in 0.1 M phosphate buffer (pH 7.0). After dilution to achieve different molar enzyme concentrations (0.5-20 µM), samples were incubated for 1 h at 30 ºC to allow for isopeptide bond formation. Specific and volumetric activities of the immobilized enzyme samples with indole (2.5 mM) and NADPH (0.25 mM) at 30 ºC and pH 7.0 were then measured with 40 µL of the immobilization mixtures in 200 µL reactions (5-fold dilutions) to obtain the final enzyme concentrations shown. Indigo formation and NADPH consumption were spectrophotometrically monitored. Control reactions were performed without scaffolds. At enzyme concentrations > 0.8 µM, NADPH was consumed after 30 s, preventing reliable activity measurements. Detailed protocols with protein concentrations are provided in the Methods. Data are shown as mean values ± SD and error bars represent the standard deviations of four replicates with one set of purified proteins.

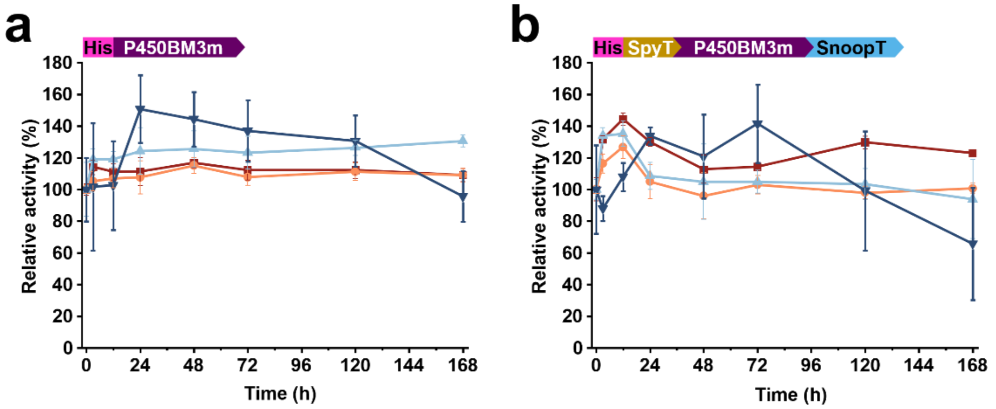

| Label | EutM Scaffold | Specific activity at t = 0 h (µM Indigo/min/µM enzyme) | |
|---|---|---|---|
| | | His-P450BM3m | His-SpyT-P450BM3m-SnoopT |
| ■ | None | 3.4 ± 0.1 | 3.3 ± 0.2 |
| ● | His-SpyC-EutM-SnoopC | 3.7 ± 0.1 | 4.0 ± 0.1 |
| ▲ | His-EutM:SpyC-EutM-SnoopC=3.9:1 | 3.1 ± 0.5 | 3.4 ± 0.2 |
| ▼ | His-EutM:SpyC-EutM-SnoopC=8.8:1 | 3.0 ± 0.6 | 2.9 ± 0.8 |

**Supplementary Fig. 16. Stability of P450BM3m mixed with EutM scaffolds.**

The influence of cross-linking to different EutM scaffolds on the stability of unmodified (control) (**a**) and dual-modified (**b**) His-P450BM3m was measured at the optimal enzyme concentration (1.6 µM) and molar ratio of 1:4 of enzyme to conjugating scaffold building block. Enzymes and scaffolds (or no scaffold controls) were mixed in 0.1 M sodium phosphate buffer (pH 7.0) and incubated at 30 °C for up to 168 h. Immediately after mixing (t = 0 h) and after different time intervals, aliquots were removed for activity measurements to determine specific activities with indole and NADPH as substrates (see Methods and **Fig. 7d** for details). Shown are specific activities measured for indigo formation at t = 0 h and relative specific activities (with t = 0 h set as 100 %) for 168 h of incubation. Data are shown as means ± SD and error bars represent the standard derivation of four replicates.

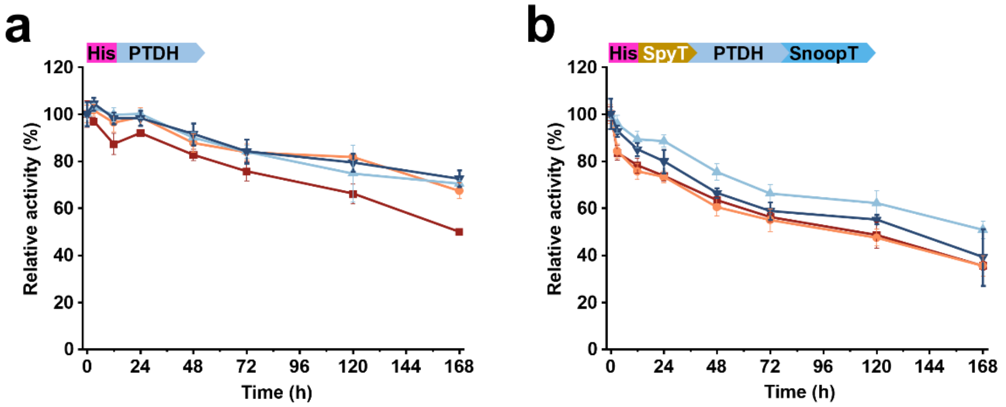

| Label | EutM Scaffold | Specific activity at t = 0 h (µM NADPH/min/µM enzyme) | |
|---|---|---|---|
| | | His-PTDH | His-SpyT-PTDH-SnoopT |
| | None | 99.1 ± 5.6 | 113.7 ± 3.4 |
| | His-SpyC-EutM-SnoopC | 126.7 ± 4.8 | 135.3 ± 5.3 |
| | His-EutM:SpyC-EutM-SnoopC=3.9:1 | 124.0 ± 5.8 | 137.4 ± 2.7 |
| | His-EutM:SpyC-EutM-SnoopC=8.8:1 | 120.5 ± 6.3 | 131.4 ± 8.4 |

**Supplementary Fig. 17. Stability of PTDH mixed with EutM scaffolds.**

The influence of cross-linking to different EutM scaffolds on the stability of unmodified (control) (**a**) and dual-modified (**b**) His-PTDH was measured at the optimal enzyme concentration (0.1 µM) and molar ratio of 1:4 of enzyme to conjugating scaffold building block. Enzymes and scaffolds (or no scaffold control) were mixed in 0.1 M sodium phosphate buffer (pH 7.0) and incubated at 30 °C for up to 168 h. Immediately after mixing (t = 0) and after different time intervals, aliquots were removed for activity measurements to determine specific activities with sodium phosphite and $NADP^+$ as substrates (see Methods and **Fig. 7d** for details). Shown are specific activities for NADPH formation at t= 0 h and relative specific activities (with t = 0 h set as 100 %) for 168 h of incubation. Data are shown as means ± SD and error bars represent the standard derivation of four replicates.

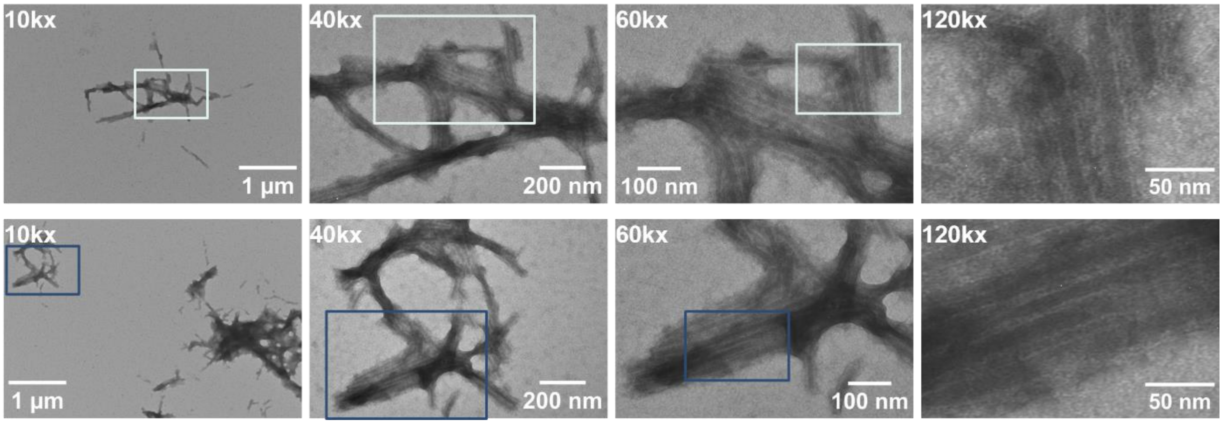

**Supplementary Fig. 18. TEM of co-immobilized P450BM3m and PTDH on hybrid scaffolds.**

Hybrid His-EutM:SpyC-EutM-SnoopC=8.8:1 scaffolds cross-linked with His-SpyT-P450BM3m-SnoopT and His-SpyT-PTDH-SnoopT from **Fig. 8a** were diluted to 1 mg/mL and applied to grids for TEM imaging. Two grid positions are shown (top and bottom rows) with images captured at a range of magnifications (10k-120kx) for each location in each row. Boxes indicate magnified regions for each of the two grid positions. Images show parallel aligned tubes that are enveloped by a film and are linked together, similar to the structures observed with GFP (compare **Fig. 6a** and **Supplementary Fig. 12**). Representative images captured from one set of purified proteins are shown.

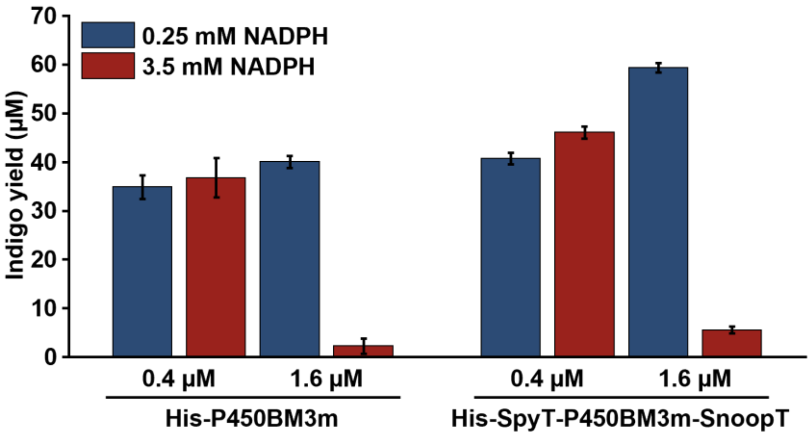

**Supplementary Fig. 19. Inhibition of indigo production by P450BM3m at higher local NADPH concentration.**

To confirm that the inhibition of indigo formation observed in the coupled reaction system with co-immobilized P450BM3m-PTDH (**Fig. 8b**) was likely caused by an uncoupling of NADPH consumption and indole oxidation at high local NADPH concentrations, P450BM3m indigo production activities of the unmodified and dual-modified enzyme were measured with 0.4 µM or 1.6 µM enzyme and with 0.25 mM or 3.5 mM NADPH. Reactions were performed as in **Fig. 8b** with 2.5 mM indole at pH 7.0 and 30 ºC. Reactions were started with the addition of NADPH. After 15 min, indigo formation was spectrophotometrically quantified. Data are shown as mean values ± SD and error bars represent the standard deviations of four replicates with one set of purified proteins.

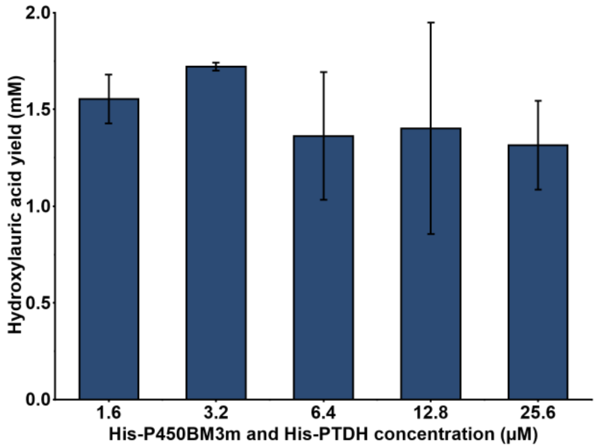

**Supplementary Fig. 20. Effect of P450BM3m and PTDH concentrations on hydroxylauric acid production.** Hydroxylauric acid production with increasing, equimolar concentrations of P450BM3m and PTDH (1.6-25.6 µM each) was evaluated in 1 mL reactions at 30 ºC and 180 rpm. The reactions contained 0.25 mM $NADP^+$, 10 mM $Na_2HPO_3$ and 2.5 mM lauric acid dissolved in DMSO (final concentration 8% v/v) in 0.1 M sodium phosphate buffer (pH 7.0). Reactions were stopped after 15 min and hydroxylauric acids were quantified by GC-FID. Data are shown as mean values ± SD and error bars represent the standard deviations of four replicates with one set of purified proteins.

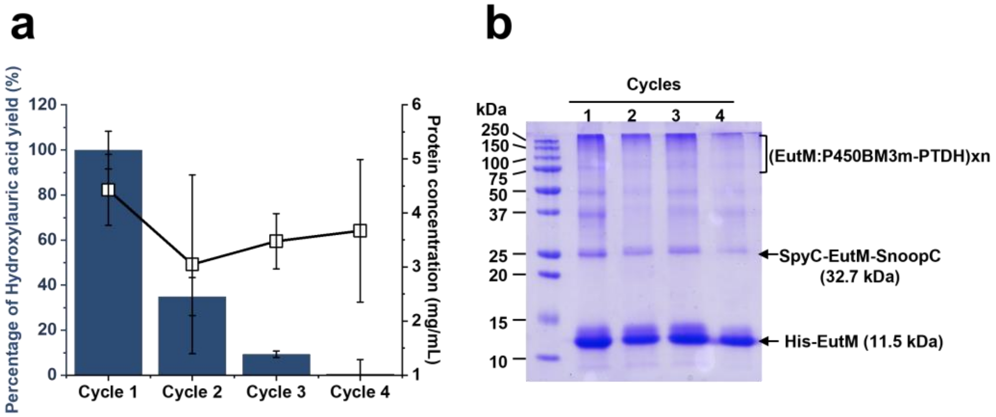

**Supplementary Fig. 21. Reuse of the EutM:P450BM3m:PTDH biomaterials for hydroxylauric acid production**

Performance of the EutM:P450BM3m:PTDH biomaterials for hydroxylauric acid production in each reaction cycle. After co-immobilizing dual-modified P450BM3m and PTDH (6.4 µM each) at a 1:4 molar ratio of enzymes to cross-linking scaffold building block onto hybrid EutM:His-SpyC-EutM-SnoopC scaffolds, 3 mL conversion reactions which consist of 3.2 µM His-SpyT-P450BM3m-SnoopT, 3.2 µM His-SpyT-PTDH-SnoopT and 25.6 µM His-EutM:SpyC-EutM-SnoopC=8.8:1, 0.1 M sodium phosphate buffer (pH 7.0), 0.25 mM $NADP^+$, 10 mM $Na_2HPO_3$ and 2 mM lauric acid added within DMSO (final concentration 8% v/v) were then performed for 30 min at 30 ºC. Two hundred µL sample were took for measurement of the yield of 9-, 10-, and 11-hydroxylauric acids and protein concentration (**a**) and SDS-PAGE analysis (**b**). The rest of the sample was spun down at 5,000 x*g* for 5 mins at 4 ºC to recover enzyme materials, and then equal volume of same reaction components were added to start next cycle. The yield of hydroxylauric acids for cycle 2, 3 and 4 were normalized to cycle 1. Data are shown as mean values ± SD and error bars represent the standard deviations of four replicates with one set of purified proteins.

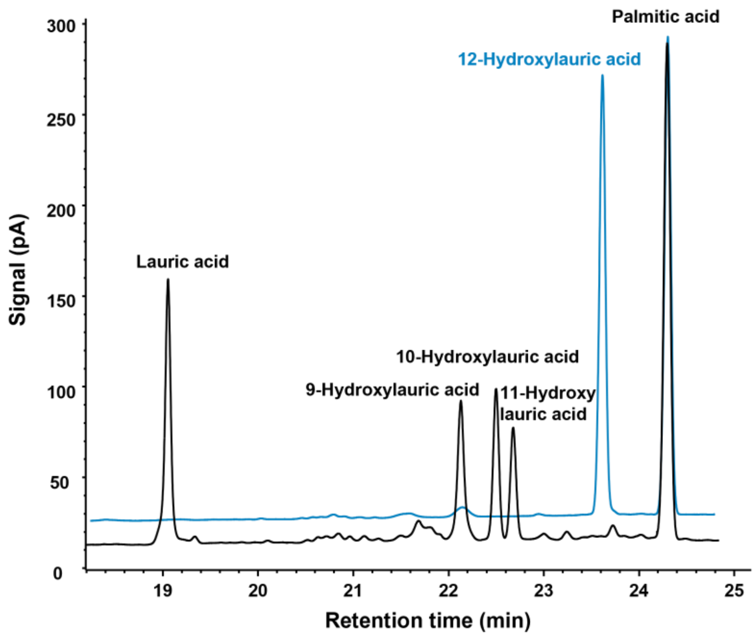

**Supplementary Fig. 22. GC-FID chromatogram for the analysis of lauric acid conversion by P450BM3m.**

Representative chromatograms are shown for the analysis of TMS derivatized lauric acid enzyme conversion reactants and products (black trace) and 12-hydroxylauric acid (response factor quantification standard) and palmitic acid (internal standard) (blue line). Shown representative coupled enzyme reaction (**Fig. 8b**) was performed with 0.8 µM His-P450BM3m,1.6 µM His-PTDH, 10 mM sodium phosphite, 0.25 mM $NADP^+$ and 2.5 mM lauric acid in a 1 mL reaction volume for 15 min. Palmitic acid was added as internal standard during TMS derivatization for GC analysis. TMS-derivatized hydroxylauric acid P450 conversion products were quantified using 12-hydroxylauric acid as reference. See Methods for details. Conversion products were identified by their mass fragmentation spectra (see **Supplementary Fig. 23**).

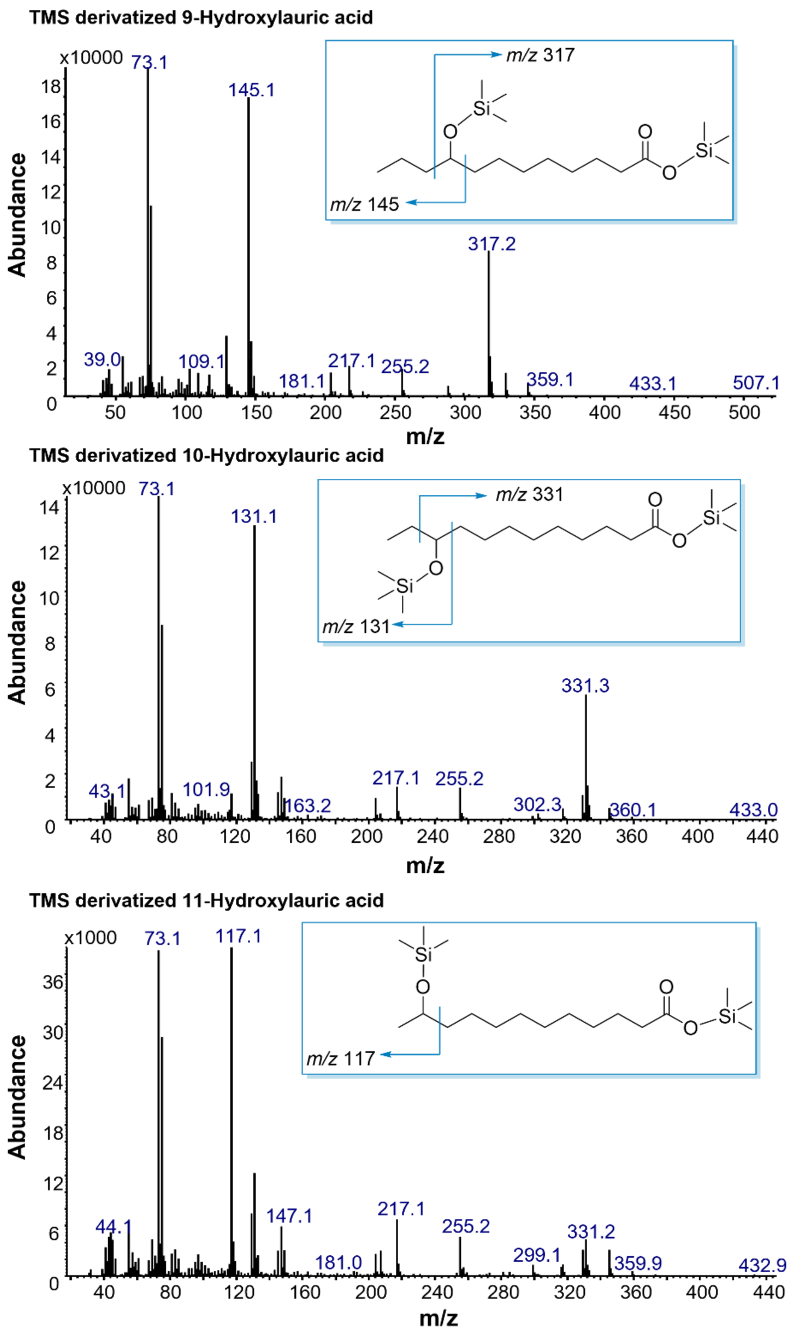

**Supplementary Fig. 23. Fragmentation spectra for TMS derivatized hydroxylauric acids.**

Representative mass fragmentation spectra obtained after GC-MS analysis of TMS-derivatized hydroxylauric acid conversion products from the P450BM3m-PTDH coupled reaction shown in **Supplementary Fig. 22**. Insets show trimethyl-silylated hydroxylauric acid structures and *m/z* of expected fragments. See Methods for details.

### Supplementary Tables

**Supplementary Table 1. Nucleotide sequences of Ribosome Binding Site (RBS).**

| RBS | Nucleic acid sequence | Predicted strength for SpyC-EutM-SnoopC | | Source |
|---|---|---|---|---|
| | | Translation Rate (au) | $\Delta G_{total}$ (kcal/mol) | |
| A | ATGAACGAAGGAGGGATCTGGATCC | 2159.7 | -1.24 | pCT5BB[3,14] |
| B | AACAAAATGAGGAGGTACTGAG | 27028.83 | -6.86 | [7] |
| C | CCCTTTAATGAAAAACAAAGATCACATCTAGACAGCATATCAATT | 597.4 | 1.61 | [7] |
| D | CGAGCTGGTACTTAAAAAC | 100.13 | 5.58 | [7] |
| E | TGAAAATAGAATAGTAACGTACGTTACCATCCATCT | 21.95 | 8.95 | [7] |

**Supplementary Table 2. Hydroxylauric acid product profiles of scaffolded and unscaffolded P450BM3m-PTDH coupled reaction system.**

Relative amounts of hydroxylauric acid products from reactions of P450BM3m and PTDH co-immobilized on different scaffolds from **Fig. 8c** are shown. Scaffolding and the type of scaffolds had no major effect on the hydroxylation site selectivity of P450 for lauric acid.

| **Scaffold** | **Relative amounts of hydroxylauric acid products (%)** | | |
|---|---|---|---|
| | **9-hydroxylauric acid** | **10-hydroxylauric acid** | **11-hydroxylauric acid** |
| None | 42.9 ± 5.6 | 34.9 ± 3.9 | 22.2 ± 1.7 |
| His-SpyC-EutM-SnoopC | 43.2 ± 3.4 | 34.1 ± 1.2 | 22.8 ± 2.2 |
| His-EutM:SpyC-EutM-SnoopC=3.9:1 | 43.7 ± 2.1 | 34.3 ± 1.3 | 22 ± 0.9 |
| His-EutM:SpyC-EutM-SnoopC=8.8:1 | 39.7 ± 2 | 36.5 ± 1.3 | 23.8 ± 0.7 |

**Supplementary Table 3. Plasmids and strains used in this study.**

| Plasmid | Description | Source |
|---|---|---|
| ***Plasmids for expression of EutM scaffold*** | | |
| pCT5BB-His-EutM | *eutM* (NP_461400) containing a N-terminal 6xHis tag fused via a GS-linker to generate N-His-GS-EutM, pCT5 promoter, $Amp^R$ | [2,4,6] |
| pCT5BB-His-SpyC-EutM-SnoopC | Derived from pCT5BB-His-EutM: EutM shell protein with N-terminal His tag and SpyCatcher and C-terminal SnoopCatcher | This study |
| pCT5BB-His-SnoopT-EutM-SpyT | Derived from pCT5BB-His-EutM: EutM shell protein with N-terminal His tag and SnoopTag and C-terminal SpyTag | This study |
| ***Plasmids for expression of hybrid His-EutM:SpyC-EutM-SnoopC scaffold*** | | |
| pCT5BB-RBSA-His-EutM-RBSA-SpyC-EutM-SnoopC | Derived from pCT5BB-His-SpyC-EutM-SnoopC: co-express His-EutM and SpyC-EutM-SnoopC under control of the same cumate inducible promoter pCT5, and their transcription initiated by their own RBSA, respectively. | This study |
| pCT5BB-RBSA-His-EutM-RBSC-SpyC-EutM-SnoopC | Derived from pCT5BB-RBSA-His-EutM-RBSA-SpyC-EutM-SnoopC: replace the RBSA of SpyC-EutM-SnoopC with RBSC | This study |
| pCT5BB-RBSA-His-EutM-RBSD-SpyC-EutM-SnoopC | Derived from pCT5BB-RBSA-His-EutM-RBSA-SpyC-EutM-SnoopC: replace the RBSA of SpyC-EutM-SnoopC with RBSD | This study |
| pCT5BB-RBSA-His-EutM-RBSE-SpyC-EutM-SnoopC | Derived from pCT5BB-RBSA-His-EutM-RBSA-SpyC-EutM-SnoopC: replace the RBSA of SpyC-EutM-SnoopC with RBSE | This study |
| pCT5BB-RBSB-His-EutM-RBSA-SpyC-EutM-SnoopC | Derived from pCT5BB-RBSA-His-EutM-RBSA-SpyC-EutM-SnoopC: replace the RBSA of His-EutM with RBSB | This study |
| pCT5BB-RBSB-His-EutM-RBSC-SpyC-EutM-SnoopC | Derived from pCT5BB-RBSA-His-EutM-RBSC-SpyC-EutM-SnoopC: replace the RBSA of His-EutM with RBSB | This study |
| pCT5BB-RBSB-His-EutM-RBSD-SpyC-EutM-SnoopC | Derived from pCT5BB-RBSA-His-EutM-RBSD-SpyC-EutM-SnoopC: replace the RBSA of His-EutM with RBSB | This study |
| pCT5BB-RBSB-His-EutM-RBSE-SpyC-EutM-SnoopC | Derived from pCT5BB-RBSA-His-EutM-RBSE-SpyC-EutM-SnoopC: replace the RBSA of His-EutM with RBSB | This study |
| ***Plasmids for expression of hybrid His-EutM:SnoopT-EutM-SpyT scaffold*** | | |
| pCT5BB-RBSA-His-EutM-RBSA-SnoopT-EutM-SpyT | Derived from pCT5BB-His-SnoopT-EutM-SpyT: co-express His-EutM and SnoopT-EutM-SpyT under control of the same cumate inducible promoter pCT5, and their transcription was initiated by their own RBSA, respectively. | This study |
| pCT5BB-RBSA-His-EutM- | Derived from pCT5BB-RBSA-His-EutM-RBSA-SnoopT-EutM-SpyT: | This study |

| Plasmid | Description | Source |
|---|---|---|
| RBSC-SnoopT-EutM-SpyT | replace the RBSA of SnoopT-EutM-SpyT with RBSC | |
| pCT5BB-RBSA-His-EutM-RBSD-SnoopT-EutM-SpyT | Derived from pCT5BB-RBSA-His-EutM-RBSA-SnoopT-EutM-SpyT: replace the RBSA of SnoopT-EutM-SpyT with RBSD | This study |
| pCT5BB-RBSA-His-EutM-RBSE-SnoopT-EutM-SpyT | Derived from pCT5BB-RBSA-His-EutM-RBSA-SnoopT-EutM-SpyT: replace the RBSA of SnoopT-EutM-SpyT with RBSE | This study |
| pCT5BB-RBSB-His-EutM-RBSA-SnoopT-EutM-SpyT | Derived from pCT5BB-RBSA-His-EutM-RBSA-SnoopT-EutM-SpyT: replace the RBSA of His-EutM with RBSB | This study |
| pCT5BB-RBSB-His-EutM-RBSC-SnoopT-EutM-SpyT | Derived from pCT5BB-RBSA-His-EutM-RBSC-SnoopT-EutM-SpyT: replace the RBSA of His-EutM with RBSB | This study |
| pCT5BB-RBSB-His-EutM-RBSD-SnoopT-EutM-SpyT | Derived from pCT5BB-RBSA-His-EutM-RBSD-SnoopT-EutM-SpyT: replace the RBSA of His-EutM with RBSB | This study |
| pCT5BB-RBSB-His-EutM-RBSE-SnoopT-EutM-SpyT | Derived from pCT5BB-RBSA-His-EutM-RBSE-SnoopT-EutM-SpyT: replace the RBSA of His-EutM with RBSB | This study |
| ***Plasmids for expression of hybrid EutM:His-SpyC-EutM-SnoopC scaffold*** | | |
| pCT5BB-RBSA-EutM-RBSA-His-SpyC-EutM-SnoopC | Derived from pCT5BB-His-SpyC-EutM-SnoopC: co-express EutM and His-SpyC-EutM-SnoopC under control of the same cumate inducible promoter pCT5, and their transcription initiated by their own RBSA, respectively. | This study |
| pCT5BB-RBSA-EutM-RBSC-His-SpyC-EutM-SnoopC | Derived from pCT5BB-RBSA-EutM-RBSA-His-SpyC-EutM-SnoopC: replace the RBSA of His-SpyC-EutM-SnoopC with RBSC | This study |
| pCT5BB-RBSA-EutM-RBSD-His-SpyC-EutM-SnoopC | Derived from pCT5BB-RBSA-EutM-RBSA-His-SpyC-EutM-SnoopC: replace the RBSA of His-SpyC-EutM-SnoopC with RBSD | This study |
| pCT5BB-RBSA-EutM-RBSE-His-SpyC-EutM-SnoopC | Derived from pCT5BB-RBSA-EutM-RBSA-His-SpyC-EutM-SnoopC: replace the RBSA of His-SpyC-EutM-SnoopC with RBSE | This study |
| pCT5BB-RBSB-EutM-RBSA-His-SpyC-EutM-SnoopC | Derived from pCT5BB-RBSA-EutM-RBSA-His-SpyC-EutM-SnoopC: replace the RBSA of EutM with RBSB | This study |
| pCT5BB-RBSB-EutM-RBSC-His-SpyC-EutM-SnoopC | Derived from pCT5BB-RBSA-EutM-RBSC-His-SpyC-EutM-SnoopC: replace the RBSA of EutM with RBSB | This study |

| Plasmid | Description | Source |
|---|---|---|
| pCT5BB-RBSB-EutM-RBSD-His-SpyC-EutM-SnoopC | Derived from pCT5BB-RBSA-EutM-RBSD-His-SpyC-EutM-SnoopC: replace the RBSA of EutM with RBSB | This study |
| ***Plasmids for expression of hybrid EutM:His-SnoopT-EutM-SpyT scaffold*** | | |
| pCT5BB-RBSA-EutM-RBSA-His-SnoopT-EutM-SpyT | Derived from pCT5BB-His-SnoopT-EutM-SpyT: co-express His-EutM and SnoopT-EutM-SpyT under control of the same cumate inducible promoter pCT5, and their transcription initiated by their own RBSA, respectively. | This study |
| pCT5BB-RBSA-EutM-RBSC-His-SnoopT-EutM-SpyT | Derived from pCT5BB-RBSA-EutM-RBSA-His-SnoopT-EutM-SpyT: replace the RBSA of His-SnoopT-EutM-SpyT with RBSC | This study |
| pCT5BB-RBSA-EutM-RBSD-His-SnoopT-EutM-SpyT | Derived from pCT5BB-RBSA-EutM-RBSA-His-SnoopT-EutM-SpyT: replace the RBSA of His-SnoopT-EutM-SpyT with RBSD | This study |
| pCT5BB-RBSA-EutM-RBSE-His-SnoopT-EutM-SpyT | Derived from pCT5BB-RBSA-EutM-RBSA-His-SnoopT-EutM-SpyT: replace the RBSA of His-SnoopT-EutM-SpyT with RBSE | This study |
| pCT5BB-RBSB-EutM-RBSA-His-SnoopT-EutM-SpyT | Derived from pCT5BB-RBSA-EutM-RBSA-His-SnoopT-EutM-SpyT: replace the RBSA of EutM with RBSB | This study |
| pCT5BB-RBSB-EutM-RBSD-His-SnoopT-EutM-SpyT | Derived from pCT5BB-RBSA-EutM-RBSD-His-SnoopT-EutM-SpyT: replace the RBSA of EutM with RBSB | This study |
| ***Plasmids for expression of reporters and cargo proteins*** | | |
| pET28a | T7 promoter, $Km^R$ (6xHis-tag and thrombin site downstream of MCS in pET28a) | Invitrogen |
| pET28a-His-GFP | Derived from pET28a: GFP fluorescence protein with N-terminal His tag and thrombin site | This study |
| pET28a-His-SnoopC-GFP-SpyC | Derived from pET28a-His-GFP: GFP fluorescence protein with N-terminal His tag, thrombin site, and SnoopCatcher and C-terminal SpyCatcher | This study |
| pET28a-His-SpyT-GFP-SnoopT | Derived from pET28a-His-GFP: GFP fluorescence protein with N-terminal His tag, thrombin site, and SpyTag and C-terminal SnoopTag | This study |

| Plasmid | Description | Source |
|---|---|---|
| pET28a-His-PTDH | Derived from pET28a-His-GFP: replace the GFP with synthesized PTDH gene fragment | This study |
| pET28a-His-SnoopC-PTDH-SpyC | Derived from pET28a-His-SnoopC-GFP-SpyC: replace the GFP with synthesized PTDH gene fragment | This study |
| pET28a-His-SpyT-PTDH-SnoopT | Derived from pET28a-His-SpyT-GFP-SnoopT: replace the GFP with synthesized PTDH gene fragment | This study |
| pET28a-His-P450BM3m | Derived from pET28a-His-GFP: replace the GFP with P450BM3m gene fragment | This study |
| pET28a-His-SnoopC-P450BM3m-SpyC | Derived from pET28a-His-SnoopC-GFP-SpyC: replace the GFP with P450BM3m gene fragment | This study |
| pET28a-His-SpyT-P450BM3m-SnoopT | Derived from pET28a-His-SpyT-GFP-SnoopT: replace the GFP with P450BM3m gene fragment | This study |
| **Strain** | **Description** | **Source** |
| *Escherichia coli* Top10 | General cloning host | Invitrogen |
| *Escherichia coli* BL21 (DE3) | Protein expression | New England Biolabs |

**Supplementary Table 4. Amino acid sequences of proteins and peptides in this study.**

| Name | Amino acid sequences |
|---|---|
| EutM | MEALGMIETRGLVALIEASDAMVKAARVKLVGVKQIGGGLCTAMVRGDVAACKAATDAGA AAAQRIGELVSVHVIPRPHGDLEEVFPISFKGDSNI |
| SpyCatcher (SpyC) | DSATHIKFSKRDEDGKELAGATMELRDSSGKTISTWISDGQVKDFYLYPGKYTFVETAAPD GYEVATAITFTVNEQGQVTVNG |
| SpyTag (SpyT) | AHIVMVDAYKPTK |
| SnoopCatcher (SnoopC) | KPLRGAVFSLQKQHPDYPDIYGAIDQNGTYQNVRTGEDGKLTFKNLSDGKYRLFENSEPA GYKPVQNKPIVAFQIVNGEVRDVTSIVPQDIPATYEFTNGKHYITNEPIPPK |
| SnoopTag (SnoopT) | KLGDIEFIKVNK |
| His-EutM | MHHHHHHGSGSGSGSGSGSEALGMIETRGLVALIEASDAMVKAARVKLVGVKQIGGGLCT AMVRGDVAACKAATDAGAAAAQRIGELVSVHVIPRPHGDLEEVFPISFKGDSNI |
| His-SpyC-EutM-SnoopC | MHHHHHHGSGSGSGSGSGDSATHIKFSKRDEDGKELAGATMELRDSSGKTISTWISDGQ VKDFYLYPGKYTFVETAAPDGYEVATAITFTVNEQGQVTVNGSGSGSGSEALGMIETRGLV ALIEASDAMVKAARVKLVGVKQIGGGLCTAMVRGDVAACKAATDAGAAAAQRIGELVSVHV IPRPHGDLEEVFPISFKGDSNIVDGSGGSGGKPLRGAVFSLQKQHPDYPDIYGAIDQNGTY QNVRTGEDGKLTFKNLSDGKYRLFENSEPAGYKPVQNKPIVAFQIVNGEVRDVTSIVPQDIP ATYEFTNGKHYITNEPIPPK |
| SpyC-EutM-SnoopC | MDSATHIKFSKRDEDGKELAGATMELRDSSGKTISTWISDGQVKDFYLYPGKYTFVETAAP DGYEVATAITFTVNEQGQVTVNGSGSGSGSEALGMIETRGLVALIEASDAMVKAARVKLVG VKQIGGGLCTAMVRGDVAACKAATDAGAAAAQRIGELVSVHVIPRPHGDLEEVFPISFKGD SNIVDGSGGSGGKPLRGAVFSLQKQHPDYPDIYGAIDQNGTYQNVRTGEDGKLTFKNLSD GKYRLFENSEPAGYKPVQNKPIVAFQIVNGEVRDVTSIVPQDIPATYEFTNGKHYITNEPIPP K |
| His-SnoopT-EutM-SpyT | MHHHHHHGSGGSGGKLGDIEFIKVNKGSGSGSGSGSGSMEALGMIETRGLVALIEASDAM VKAARVKLVGVKQIGGGLCTAMVRGDVAACKAATDAGAAAAQRIGELVSVHVIPRPHGDL EEVFPISFKGDSNIGSGGGSGGGSGSAHIVMVDAYKPTK |
| SnoopT-EutM-SpyT | MKLGDIEFIKVNKGSGSGSGSGSGSMEALGMIETRGLVALIEASDAMVKAARVKLVGVKQI GGGLCTAMVRGDVAACKAATDAGAAAAQRIGELVSVHVIPRPHGDLEEVFPISFKGDSNIG SGGGSGGGSGSAHIVMVDAYKPTK |
| His-Thrombin-GFP | MGSSHHHHHHSSGLVPRGSHMMSKGEELFTGVVPILVELDGDVNGHKFSVRGEGEGDAT NGKLTLKFICTTGKLPVPWPTLVTTLTYGVQCFSRYPDHMKRHDFFKSAMPEGYVQERTIS FKDDGTYKTRAEVKFEGDTLVNRIELKGIDFKEDGNILGHKLEYNFNSHNVYITADKQKNGI KANFKIRHNVEDGSVQLADHYQQNTPIGDGPVLLPDNHYLSTQSVLSKDPNEKRDHMVLL |

| Name | Amino acid sequences |
|---|---|
| | EFVTAAGITHGMDELYK |
| His-Thrombin-SnoopC-GFP-SpyC | MGSSHHHHHHSSGLVPRGSHMGSGGSGGKPLRGAVFSLQKQHPDYPDIYGAIDQNGTYQNVRTGEDGKLTFKNLSDGKYRLFENSEPAGYKPVQNKPIVAFQIVNGEVRDVTSIVPQDIPATYEFTNGKHYITNEPIPPKGGSGGSGMSKGEELFTGVVPILVELDGDVNGHKFSVRGEGEGDATNGKLTLKFICTTGKLPVPWPTLVTTLTYGVQCFSRYPDHMKRHDFFKSAMPEGYVQERTISFKDDGTYKTRAEVKFEGDTLVNRIELKGIDFKEDGNILGHKLEYNFNSHNVYITADKQKNGIKANFKIRHNVEDGSVQLADHYQQNTPIGDGPVLLPDNHYLSTQSVLSKDPNEKRDHMVLLEFVTAAGITHGMDELYKSGSGSGGDSATHIKFSKRDEDGKELAGATMELRDSSGKTISTWISDGQVKDFYLYPGKYTFVETAAPDGYEVATAITFTVNEQGQVTVNG |
| His-Thrombin-SpyT-GFP-SnoopT | MGSSHHHHHHSSGLVPRGSHMMGSSGAHIVMVDAYKPTKGSGGSGMSKGEELFTGVVPILVELDGDVNGHKFSVRGEGEGDATNGKLTLKFICTTGKLPVPWPTLVTTLTYGVQCFSRYPDHMKRHDFFKSAMPEGYVQERTISFKDDGTYKTRAEVKFEGDTLVNRIELKGIDFKEDGNILGHKLEYNFNSHNVYITADKQKNGIKANFKIRHNVEDGSVQLADHYQQNTPIGDGPVLLPDNHYLSTQSVLSKDPNEKRDHMVLLEFVTAAGITHGMDELYKSGSGSGGKLGDIEFIKVNK |
| His-Thrombin-PTDH | MGSSHHHHHHSSGLVPRGSHMMLPKLVITHRVHEEILQLLAPHCELITNQTDSTLTREEILRRCRDAQAMMAFMPDRVDADFLQACPELRVIGCALKGFDNFDVDACTARGVWLTFVPDLLTVPTAELAIGLAVGLGRHLRAADAFVRSGKFRGWQPRFYGTGLDNATVGFLGMGAIGLAMADRLQGWGATLQYHARKALDTQTEQRLGLRQVACSELFASSDFILLALPLNADTLHLVNAELLALVRPGALLVNPCRGSVVDEAAVLAALERGQLGGYAADVFEMEDWARADRPQQIDPALLAHPNTLFTPHIGSAVRAVRLEIERCAAQNILQALAGERPINAVNRLPKANPAADSRSAAG |
| His-Thrombin-SnoopC-PTDH-SpyC | MGSSHHHHHHSSGLVPRGSHMGSGGSGGKPLRGAVFSLQKQHPDYPDIYGAIDQNGTYQNVRTGEDGKLTFKNLSDGKYRLFENSEPAGYKPVQNKPIVAFQIVNGEVRDVTSIVPQDIPATYEFTNGKHYITNEPIPPKGGSGGSGMLPKLVITHRVHEEILQLLAPHCELITNQTDSTLTREEILRRCRDAQAMMAFMPDRVDADFLQACPELRVIGCALKGFDNFDVDACTARGVWLTFVPDLLTVPTAELAIGLAVGLGRHLRAADAFVRSGKFRGWQPRFYGTGLDNATVGFLGMGAIGLAMADRLQGWGATLQYHARKALDTQTEQRLGLRQVACSELFASSDFILLALPLNADTLHLVNAELLALVRPGALLVNPCRGSVVDEAAVLAALERGQLGGYAADVFEMEDWARADRPQQIDPALLAHPNTLFTPHIGSAVRAVRLEIERCAAQNILQALAGERPINAVNRLPKANPAADSRSAAGSGSGSGGDSATHIKFSKRDEDGKELAGATMELRDSSGKTISTWISDGQVKDFYLYPGKYTFVETAAPDGYEVATAITFTVNEQGQVTVNG |
| His-Thrombin-SpyT-PTDH-SnoopT | MGSSHHHHHHSSGLVPRGSHMMGSSGAHIVMVDAYKPTKGSGGSGMLPKLVITHRVHEEILQLLAPHCELITNQTDSTLTREEILRRCRDAQAMMAFMPDRVDADFLQACPELRVIGCALKGFDNFDVDACTARGVWLTFVPDLLTVPTAELAIGLAVGLGRHLRAADAFVRSGKFRGWQPRFYGTGLDNATVGFLGMGAIGLAMADRLQGWGATLQYHARKALDTQTEQRLGLRQVACSELFASSDFILLALPLNADTLHLVNAELLALVRPGALLVNPCRGSVVDEAAVLAALERGQLGGYAADVFEMEDWARADRPQQIDPALLAHPNTLFTPHIGSAVRAVRLEIERCAAQNILQALAGE |

| Name | Amino acid sequences |
| --- | --- |
| | RPINAVNRLPKANPAADSRSAAGSGSGSGGKLGDIEFIKVNK |
| His-Thrombin-P450BM3m | MGSSHHHHHHSSGLVPRGSHMMTIKEMPQPKTFGELKNLPLLNTDKPVQALMKIADELGEI<br>FKFEAPGRVTRYLSSQRLIKEACDESRFDKNLSQGLKFVRDFAGDGLVTSWTHEKNWKKA<br>HNILLPSFSQQAMKGYHAMMVDIAVQLVQKWERLNADEHIEVPEDMTRLTLDTIGLCGFNY<br>RFNSFYRDQPHPFITSMVRALDEAMNKQQRANPDDPAYDENKRQFQEDIKVMNDLVDKIIA<br>DRKASGEQSDDLLTHMLNGKDPETGEPLDDENIRYQIITFLIAGHETTSGLLSFALYFLVKNP<br>HVLQKAAEEAARVLVDPVPSYKQVKQLKYVGMVLNEALRLWPTAPAFSLYAKEDTVLGGE<br>YPLEKGDELMVLIPQLHRDKTIWGDDVEEFRPERFENPSAIPQHAFKPFGNGQRACIGQQF<br>ALHEATLVLGMMLKHFDFEDHTNYELDIKETLTLKPEGFVVKAKSKKIPLGGIPSPSTEQSAK<br>KVRKKAENAHNTPLLVLYGSNMGTAEGTARDLADIAMSKGFAPQVATLDSHAGNLPREGA<br>VLIVTASYNGHPPDNAKQFVDWLDQASADEVKGVRYSVFGCGDKNWATTYQKVPAFIDET<br>LAAKGAENIADRGEADASDDFEGTYEEWREHMWSDVAAYFNLDIENSEDNKSTLSLQFVD<br>SAADMPLAKMHGAFSTNVVASKELQQPGSARSTRHLEIELPKEASYQEGDHLGVIPRNYE<br>GIVNRVTARFGLDASQQIRLEAEEEKLAHLPLAKTVSVEELLQYVELQDPVTRTQLRAMAAK<br>TVCPPHKVELEALLEKQAYKEQVLAKRLTMLELLEKYPACEMKFSEFIALLPSIRPRYYSISS<br>SPRVDEKQASITVSVVSGEAWSGYGEYKGIASNYLAELQEGDTITCFISTPQSEFTLPKDPE<br>TPLIMVGPGTGVAPFRGFVQARKQLKEQGQSLGEAHLYFGCRSPHEDYLYQEELENAQSE<br>GIITLHTAFSRMPNQPKTYVQHVMEQDGKKLIELLDQGAHFYICGDGSQMAPAVEATLMKS<br>YADVHQVSEADARLWLQQLEEKGRYAKDVWAG |
| His-Thrombin-SnoopC-P450BM3m-SpyC | MGSSHHHHHHSSGLVPRGSHMGSGGSGGKPLRGAVFSLQKQHPDYPDIYGAIDQNGTY<br>QNVRTGEDGKLTFKNLSDGKYRLFENSEPAGYKPVQNKPIVAFQIVNGEVRDVTSIVPQDIP<br>ATYEFTNGKHYITNEPIPPKGGSGGSGMTIKEMPQPKTFGELKNLPLLNTDKPVQALMKIAD<br>ELGEIFKFEAPGRVTRYLSSQRLIKEACDESRFDKNLSQGLKFVRDFAGDGLVTSWTHEKN<br>WKKAHNILLPSFSQQAMKGYHAMMVDIAVQLVQKWERLNADEHIEVPEDMTRLTLDTIGLC<br>GFNYRFNSFYRDQPHPFITSMVRALDEAMNKQQRANPDDPAYDENKRQFQEDIKVMNDL<br>VDKIIADRKASGEQSDDLLTHMLNGKDPETGEPLDDENIRYQIITFLIAGHETTSGLLSFALYF<br>LVKNPHVLQKAAEEAARVLVDPVPSYKQVKQLKYVGMVLNEALRLWPTAPAFSLYAKEDT<br>VLGGEYPLEKGDELMVLIPQLHRDKTIWGDDVEEFRPERFENPSAIPQHAFKPFGNGQRA<br>CIGQQFALHEATLVLGMMLKHFDFEDHTNYELDIKETLTLKPEGFVVKAKSKKIPLGGIPSPS<br>TEQSAKKVRKKAENAHNTPLLVLYGSNMGTAEGTARDLADIAMSKGFAPQVATLDSHAGN<br>LPREGAVLIVTASYNGHPPDNAKQFVDWLDQASADEVKGVRYSVFGCGDKNWATTYQKV<br>PAFIDETLAAKGAENIADRGEADASDDFEGTYEEWREHMWSDVAAYFNLDIENSEDNKSTL<br>SLQFVDSAADMPLAKMHGAFSTNVVASKELQQPGSARSTRHLEIELPKEASYQEGDHLGVI<br>PRNYEGIVNRVTARFGLDASQQIRLEAEEEKLAHLPLAKTVSVEELLQYVELQDPVTRTQLR<br>AMAAKTVCPPHKVELEALLEKQAYKEQVLAKRLTMLELLEKYPACEMKFSEFIALLPSIRPR<br>YYSISSSPRVDEKQASITVSVVSGEAWSGYGEYKGIASNYLAELQEGDTITCFISTPQSEFTL |

| Name | Amino acid sequences |
| --- | --- |
| | PKDPETPLIMVGPGTGVAPFRGFVQARKQLKEQGQSLGEAHLYFGCRSPHEDYLYQEELENAQSEGIITLHTAFSRMPNQPKTYVQHVMEQDGKKLIELLDQGAHFYICGDGSQMAPAVEATLMKSYADVHQVSEADARLWLQQLEEKGRYAKDVWAGSGSGSGGDSATHIKFSKRDEDGKELAGATMELRDSSGKTISTWISDGQVKDFYLYPGKYTFVETAAPDGYEVATAITFTVNEQGQVTVNG |
| His-<u>Thrombin</u>-SpyT-P450BM3m-SnoopT | MGSSHHHHHHSSG<u>LVPRGS</u>HMMGSSGAHIVMVDAYKPTKGSGGSGMTIKEMPQPKTFGELKNLPLLNTDKPVQALMKIADELGEIFKFEAPGRVTRYLSSQRLIKEACDESRFDKNLSQGLKFVRDFAGDGLVTSWTHEKNWKKAHNILLPSFSQQAMKGYHAMMVDIAVQLVQKWERLNADEHIEVPEDMTRLTLDTIGLCGFNYRFNSFYRDQPHPFITSMVRALDEAMNKQQRANPDDPAYDENKRQFQEDIKVMNDLVDKIIADRKASGEQSDDLLTHMLNGKDPETGEPLDDENIRYQIITFLIAGHETTSGLLSFALYFLVKNPHVLQKAAEEAARVLVDPVPSYKQVKQLKYVGMVLNEALRLWPTAPAFSLYAKEDTVLGGEYPLEKGDELMVLIPQLHRDKTIWGDDVEEFRPERFENPSAIPQHAFKPFGNGQRACIGQQFALHEATLVLGMMLKHFDFEDHTNYELDIKETLTLKPEGFVVKAKSKKIPLGGIPSPSTEQSAKKVRKKAENAHNTPLLVLYGSNMGTAEGTARDLADIAMSKGFAPQVATLDSHAGNLPREGAVLIVTASYNGHPPDNAKQFVDWLDQASADEVKGVRYSVFGCGDKNWATTYQKVPAFIDETLAAKGAENIADRGEADASDDFEGTYEEWREHMWSDVAAYFNLDIENSEDNKSTLSLQFVDSAADMPLAKMHGAFSTNVVASKELQQPGSARSTRHLEIELPKEASYQEGDHLGVIPRNYEGIVNRVTARFGLDASQQIRLEAEEEKLAHLPLAKTVSVEELLQYVELQDPVTRTQLRAMAAKTVCPPHKVELEALLEKQAYKEQVLAKRLTMLELLEKYPACEMKFSEFIALLPSIRPRYYSISSSPRVDEKQASITVSVVSGEAWSGYGEYKGIASNYLAELQEGDTITCFISTPQSEFTLPKDPETPLIMVGPGTGVAPFRGFVQARKQLKEQGQSLGEAHLYFGCRSPHEDYLYQEELENAQSEGIITLHTAFSRMPNQPKTYVQHVMEQDGKKLIELLDQGAHFYICGDGSQMAPAVEATLMKSYADVHQVSEADARLWLQQLEEKGRYAKDVWAGSGSGSGGKLGDIEFIKVNK |

**Supplementary Table 5. Nucleotides sequences of proteins and peptides in this study.**

| Name | Nucleotide sequence |
|---|---|
| EutM | **ATG**GAAGCATTAGGAATGATTGAAACCCGGGGCCTGGTTGCGCTGATTGAGGCCTCCGATG<br>CGATGGTAAAAGCCGCGCGCGTGAAGCTGGTCGGCGTGAAGCAGATTGGCGGTGGCCTGT<br>GTACTGCCATGGTGCGTGGCGATGTGGCGGCGTGCAAAGCCGCAACCGATGCTGGCGCCG<br>CTGCGGCGCAGCGCATTGGCGAGTTGGTCTCCGTACACGTGATTCCACGCCCGCACGGCG<br>ATCTGGAAGAAGTGTTCCCGATCAGCTTCAAAGGCGACAGCAACATT |
| SpyCatcher (SpyC) | GATAGTGCTACCCATATTAAATTCTCAAAACGTGATGAGGACGGCAAAGAGTTAGCTGGTGC<br>AACTATGGAGTTGCGTGATTCATCTGGTAAAACTATTAGTACATGGATTTCAGATGGACAAGT<br>GAAAGATTTCTACCTGTATCCAGGAAAATATACATTTGTCGAAACCGCAGCACCAGACGGTTA<br>TGAGGTAGCAACTGCTATTACCTTTACAGTTAATGAGCAAGGTCAGGTTACTGTAAATGGC |
| SpyTag (SpyT) | GCCCACATCGTGATGGTGGACGCCTACAAGCCGACGAAG |
| SnoopCatcher (SnoopC) | AAGCCGCTGCGTGGTGCCGTGTTTAGCCTGCAGAAACAGCATCCCGACTATCCCGATATCTA<br>TGGCGCGATTGATCAGAATGGGACCTATCAAAATGTGCGTACCGGCGAAGATGGTAAACTGA<br>CCTTTAAGAATCTGAGCGATGGCAAATATCGCCTGTTTGAAAATAGCGAACCCGCTGGCTAT<br>AAACCGGTGCAGAATAAGCCGATTGTGGCGTTTCAGATTGTGAATGGCGAAGTGCGTGATGT<br>GACCAGCATTGTGCCGCAGGATATTCCGGCTACATATGAATTTACCAACGGTAAACATTATAT<br>CACCAATGAACCGATACCGCCGAAA |
| SnoopTag (SnoopT) | AAACTGGGCGATATTGAATTTATTAAAGTGAACAAA |
| His-EutM | **ATG**CATCATCATCACCACCACGGTTCTGGTTCTGGTTCTGGTTCTGGTTCTGGTTCTGAAGC<br>ATTAGGAATGATTGAAACCCGGGGCCTGGTTGCGCTGATTGAGGCCTCCGATGCGATGGTA<br>AAAGCCGCGCGCGTGAAGCTGGTCGGCGTGAAGCAGATTGGCGGTGGCCTGTGTACTGCC<br>ATGGTGCGTGGCGATGTGGCGGCGTGCAAAGCCGCAACCGATGCTGGCGCCGCTGCGGCG<br>CAGCGCATTGGCGAGTTGGTCTCCGTACACGTGATTCCACGCCCGCACGGCGATCTGGAAG<br>AAGTGTTCCCGATCAGCTTCAAAGGCGACAGCAACATTTGA |
| His-SpyC-EutM-SnoopC | **ATG**CATCATCATCACCACCACGGTTCTGGTTCTGGCAGCGGATCTGGCAGCGGAGATAGTG<br>CTACCCATATTAAATTCTCAAAACGTGATGAGGACGGCAAAGAGTTAGCTGGTGCAACTATG<br>GAGTTGCGTGATTCATCTGGTAAAACTATTAGTACATGGATTTCAGATGGACAAGTGAAAGAT<br>TTCTACCTGTATCCAGGAAAATATACATTTGTCGAAACCGCAGCACCAGACGGTTATGAGGTA<br>GCAACTGCTATTACCTTTACAGTTAATGAGCAAGGTCAGGTTACTGTAAATGGCTCTGGTTCT<br>GGTTCTGGTTCTGAAGCATTAGGAATGATTGAAACCCGGGGCCTGGTTGCGCTGATTGAGG<br>CCTCCGATGCGATGGTAAAAGCCGCGCGCGTGAAGCTGGTCGGCGTGAAGCAGATTGGCG<br>GTGGCCTGTGTACTGCCATGGTGCGTGGCGATGTGGCGGCGTGCAAAGCCGCAACCGATG<br>CTGGCGCCGCTGCGGCGCAGCGCATTGGCGAGTTGGTCTCCGTACACGTGATTCCACGCC<br>CGCACGGCGATCTGGAAGAAGTGTTCCCGATCAGCTTCAAAGGCGACAGCAACATTGTCGA<br>CGGGAGTGGTGGCAGCGGAGGCAAGCCGCTGCGTGGTGCCGTGTTTAGCCTGCAGAAACA<br>GCATCCCGACTATCCCGATATCTATGGCGCGATTGATCAGAATGGGACCTATCAAAATGTGC |

| Name | Nucleotide sequence |
| --- | --- |
| | GTACCGGCGAAGATGGTAAACTGACCTTTAAGAATCTGAGCGATGGCAAATATCGCCTGTTT<br>GAAAATAGCGAACCCGCTGGCTATAAACCGGTGCAGAATAAGCCGATTGTGGCGTTTCAGAT<br>TGTGAATGGCGAAGTGCGTGATGTGACCAGCATTGTGCCGCAGGATATTCCGGCTACATATG<br>AATTTACCAACGGTAAACATTATATCACCAATGAACCGATACCGCCGAAATAA |
| SpyC-EutM-SnoopC | **ATG**GATAGTGCTACCCATATTAAATTCTCAAAACGTGATGAGGACGGCAAAGAGTTAGCTGG<br>TGCAACTATGGAGTTGCGTGATTCATCTGGTAAAACTATTAGTACATGGATTTCAGATGGACA<br>AGTGAAAGATTTCTACCTGTATCCAGGAAAATATACATTTGTCGAAACCGCAGCACCAGACG<br>GTTATGAGGTAGCAACTGCTATTACCTTTACAGTTAATGAGCAAGGTCAGGTTACTGTAAATG<br>GCTCTGGTTCTGGTTCTGGTTCTGAAGCATTAGGAATGATTGAAACCCGGGGCCTGGTTGCG<br>CTGATTGAGGCCTCCGATGCGATGGTAAAAGCCGCGCGCGTGAAGCTGGTCGGCGTGAAG<br>CAGATTGGCGGTGGCCTGTGTACTGCCATGGTGCGTGGCGATGTGGCGGCGTGCAAAGCC<br>GCAACCGATGCTGGCGCCGCTGCGGCGCAGCGCATTGGCGAGTTGGTCTCCGTACACGTG<br>ATTCCACGCCCGCACGGCGATCTGGAAGAAGTGTTCCCGATCAGCTTCAAAGGCGACAGCA<br>ACATTGTCGACGGGAGTGGTGGCAGCGGAGGCAAGCCGCTGCGTGGTGCCGTGTTTAGCC<br>TGCAGAAACAGCATCCCGACTATCCCGATATCTATGGCGCGATTGATCAGAATGGGACCTAT<br>CAAAATGTGCGTACCGGCGAAGATGGTAAACTGACCTTTAAGAATCTGAGCGATGGCAAATA<br>TCGCCTGTTTGAAAATAGCGAACCCGCTGGCTATAAACCGGTGCAGAATAAGCCGATTGTGG<br>CGTTTCAGATTGTGAATGGCGAAGTGCGTGATGTGACCAGCATTGTGCCGCAGGATATTCCG<br>GCTACATATGAATTTACCAACGGTAAACATTATATCACCAATGAACCGATACCGCCGAAATAA |
| His-SnoopT-EutM-SpyT | **ATG**CATCATCATCACCACCACGGGAGTGGTGGCAGCGGAGGCAAACTGGGCGATATTGAAT<br>TTATTAAAGTGAACAAAGGCAGCGGTAGCGGCAGCGGTAGCGGCAGCGGTAGCATGGAAGC<br>ATTAGGAATGATTGAAACCCGGGGCCTGGTTGCGCTGATTGAGGCCTCCGATGCGATGGTA<br>AAAGCCGCGCGCGTGAAGCTGGTCGGCGTGAAGCAGATTGGCGGTGGCCTGTGTACTGCC<br>ATGGTGCGTGGCGATGTGGCGGCGTGCAAAGCCGCAACCGATGCTGGCGCCGCTGCGGCG<br>CAGCGCATTGGCGAGTTGGTCTCCGTACACGTGATTCCACGCCCGCACGGCGATCTGGAAG<br>AAGTGTTCCCGATCAGCTTCAAAGGCGACAGCAACATTGGATCCGGTGGAGGGTCTGGTGG<br>CGGATCAGGAAGTGCCCACATCGTGATGGTGGACGCCTACAAGCCGACGAAGTAA |
| SnoopT-EutM-SpyT | **ATG**AAACTGGGCGATATTGAATTTATTAAAGTGAACAAAGGCAGCGGTAGCGGCAGCGGTAG<br>CGGCAGCGGTAGCATGGAAGCATTAGGAATGATTGAAACCCGGGGCCTGGTTGCGCTGATT<br>GAGGCCTCCGATGCGATGGTAAAAGCCGCGCGCGTGAAGCTGGTCGGCGTGAAGCAGATT<br>GGCGGTGGCCTGTGTACTGCCATGGTGCGTGGCGATGTGGCGGCGTGCAAAGCCGCAACC<br>GATGCTGGCGCCGCTGCGGCGCAGCGCATTGGCGAGTTGGTCTCCGTACACGTGATTCCAC<br>GCCCGCACGGCGATCTGGAAGAAGTGTTCCCGATCAGCTTCAAAGGCGACAGCAACATTGG<br>ATCCGGTGGAGGGTCTGGTGGCGGATCAGGAAGTGCCCACATCGTGATGGTGGACGCCTA<br>CAAGCCGACGAAGTAA |

| Name | Nucleotide sequence |
|---|---|
| His-Thrombin-GFP | **ATG**GGCAGCAGCCATCATCATCATCATCACAGCAGCGGCCTGGTGCCGCGCGGCAGCCATA<br>TGATGTCAAAAGGAGAAGAACTTTTTACAGGTGTAGTACCTATCTTGGTTGAATTGGATGGTG<br>ATGTTAACGGTCACAAATTTTCTGTACGTGGTGAAGGTGAAGGTGATGCAACTAACGGTAAAT<br>TGACACTTAAATTCATTTGTACAACTGGAAAACTTCCTGTTCCTTGGCCTACTCTTGTTACAAC<br>ATTGACATATGGAGTACAATGTTTTTCACGTTATCCTGATCATATGAAACGTCACGATTTTTTT<br>AAATCTGCTATGCCAGAAGGTTATGTACAAGAACGTACAATTTCATTTAAAGATGACGGAACA<br>TATAAAACACGTGCTGAAGTAAAATTCGAAGGTGACACTCTTGTTAATCGTATCGAATTGAAA<br>GGAATCGATTTCAAAGAAGATGGTAACATTTTGGGACACAAACTTGAATACAACTTCAACTCT<br>CATAATGTTTATATCACAGCTGACAAACAAAAAAACGGTATTAAAGCTAATTTTAAAATTCGTC<br>ACAATGTTGAAGATGGATCTGTTCAATTGGCTGATCATTATCAACAAAATACACCAATCGGAG<br>ACGGACCAGTATTGCTTCCAGATAACCACTACCTTTCTACTCAATCAGTTCTTTCAAAAGATC<br>CTAACGAAAAACGTGACCATATGGTACTTCTTGAATTTGTTACAGCAGCAGGTATCACTCACG<br>GTATGGACGAACTTTATAAATAA |
| His-Thrombin-SnoopC-GFP-SpyC | **ATG**GGCAGCAGCCATCATCATCATCATCACAGCAGCGGCCTGGTGCCGCGCGGCAGCCATA<br>TGGGGAGTGGTGGCAGCGGAGGCAAGCCGCTGCGTGGTGCCGTGTTTAGCCTGCAGAAAC<br>AGCATCCCGACTATCCCGATATCTATGGCGCGATTGATCAGAATGGGACCTATCAAAATGTG<br>CGTACCGGCGAAGATGGTAAACTGACCTTTAAGAATCTGAGCGATGGCAAATATCGCCTGTT<br>TGAAAATAGCGAACCCGCTGGCTATAAACCGGTGCAGAATAAGCCGATTGTGGCGTTTCAGA<br>TTGTGAATGGCGAAGTGCGTGATGTGACCAGCATTGTGCCGCAGGATATTCCGGCTACATAT<br>GAATTTACCAACGGTAAACATTATATCACCAATGAACCGATACCGCCGAAAGGAGGTTCAGG<br>CGGTTCAGGGATGTCAAAAGGAGAAGAACTTTTTACAGGTGTAGTACCTATCTTGGTTGAATT<br>GGATGGTGATGTTAACGGTCACAAATTTTCTGTACGTGGTGAAGGTGAAGGTGATGCAACTA<br>ACGGTAAATTGACACTTAAATTCATTTGTACAACTGGAAAACTTCCTGTTCCTTGGCCTACTCT<br>TGTTACAACATTGACATATGGAGTACAATGTTTTTCACGTTATCCTGATCATATGAAACGTCAC<br>GATTTTTTTAAATCTGCTATGCCAGAAGGTTATGTACAAGAACGTACAATTTCATTTAAAGATG<br>ACGGAACATATAAAACACGTGCTGAAGTAAAATTCGAAGGTGACACTCTTGTTAATCGTATCG<br>AATTGAAAGGAATCGATTTCAAAGAAGATGGTAACATTTTGGGACACAAACTTGAATACAACT<br>TCAACTCTCATAATGTTTATATCACAGCTGACAAACAAAAAAACGGTATTAAAGCTAATTTTAA<br>AATTCGTCACAATGTTGAAGATGGATCTGTTCAATTGGCTGATCATTATCAACAAAATACACC<br>AATCGGAGACGGACCAGTATTGCTTCCAGATAACCACTACCTTTCTACTCAATCAGTTCTTTC<br>AAAAGATCCTAACGAAAAACGTGACCATATGGTACTTCTTGAATTTGTTACAGCAGCAGGTAT<br>CACTCACGGTATGGACGAACTTTATAAATCCGGATCAGGATCCGGCGGCGATAGTGCTACCC<br>ATATTAAATTCTCAAAACGTGATGAGGACGGCAAAGAGTTAGCTGGTGCAACTATGGAGTTG<br>CGTGATTCATCTGGTAAAACTATTAGTACATGGATTTCAGATGGACAAGTGAAAGATTTCTAC<br>CTGTATCCAGGAAAATATACATTTGTCGAAACCGCAGCACCAGACGGTTATGAGGTAGCAAC<br>TGCTATTACCTTTACAGTTAATGAGCAAGGTCAGGTTACTGTAAATGGCTAA |

| **Name** | **Nucleotide sequence** |
| --- | --- |
| His-Thrombin-SpyT-GFP-SnoopT | **ATG**GGCAGCAGCCATCATCATCATCATCACAGCAGCGGCCTGGTGCCGCGCGGCAGCCATA<br>TGATGGGCAGCAGCGGCGCCCACATCGTGATGGTGGACGCCTACAAGCCGACGAAGGGTT<br>CAGGGGGATCCGGTATGTCAAAAGGAGAAGAACTTTTTACAGGTGTAGTACCTATCTTGGTT<br>GAATTGGATGGTGATGTTAACGGTCACAAATTTTCTGTACGTGGTGAAGGTGAAGGTGATGC<br>AACTAACGGTAAATTGACACTTAAATTCATTTGTACAACTGGAAAACTTCCTGTTCCTTGGCCT<br>ACTCTTGTTACAACATTGACATATGGAGTACAATGTTTTTCACGTTATCCTGATCATATGAAAC<br>GTCACGATTTTTTTAAATCTGCTATGCCAGAAGGTTATGTACAAGAACGTACAATTTCATTTAA<br>AGATGACGGAACATATAAAACACGTGCTGAAGTAAAATTCGAAGGTGACACTCTTGTTAATCG<br>TATCGAATTGAAAGGAATCGATTTCAAAGAAGATGGTAACATTTTGGGACACAAACTTGAATA<br>CAACTTCAACTCTCATAATGTTTATATCACAGCTGACAAACAAAAAAACGGTATTAAAGCTAAT<br>TTTAAAATTCGTCACAATGTTGAAGATGGATCTGTTCAATTGGCTGATCATTATCAACAAAATA<br>CACCAATCGGAGACGGACCAGTATTGCTTCCAGATAACCACTACCTTTCTACTCAATCAGTTC<br>TTTCAAAAGATCCTAACGAAAAACGTGACCATATGGTACTTCTTGAATTTGTTACAGCAGCAG<br>GTATCACTCACGGTATGGACGAACTTTATAAATCCGGATCAGGATCCGGCGGCAAACTGGGC<br>GATATTGAATTTATTAAAGTGAACAAATAA |
| His-Thrombin-PTDH | **ATG**GGCAGCAGCCATCATCATCATCATCACAGCAGCGGCCTGGTGCCGCGCGGCAGCCATA<br>TGATGCTGCCGAAACTGGTCATCACCCATCGTGTGCACGAAGAAATCCTGCAACTGCTGGCT<br>CCGCATTGCGAACTGATTACCAATCAGACCGACAGCACTCTGACCCGTGAAGAAATTCTGCG<br>TCGCTGCCGTGACGCACAGGCAATGATGGCGTTTATGCCTGATCGTGTGGATGCTGATTTCC<br>TGCAGGCTTGTCCGGAACTGCGCGTAATTGGCTGCGCGCTGAAAGGTTTCGACAATTTCGA<br>CGTTGATGCGTGTACTGCTCGTGGCGTGTGGCTGACCTTCGTTCCGGACCTGCTGACTGTG<br>CCGACTGCGGAACTGGCTATCGGCCTGGCGGTTGGTCTGGGCCGTCACCTGCGTGCAGCA<br>GACGCGTTTGTTCGTTCTGGTAAATTTCGCGGCTGGCAGCCTCGCTTTTACGGTACCGGTCT<br>GGACAACGCGACCGTGGGCTTCCTGGGTATGGGTGCAATCGGTCTGGCGATGGCTGATCGT<br>CTGCAGGGTTGGGGCGCTACCCTGCAATACCACGCGCGTAAAGCTCTGGATACGCAAACGG<br>AACAACGCCTGGGTCTGCGTCAAGTGGCTTGCAGCGAGCTGTTCGCTTCTAGCGACTTCATT<br>CTGCTGGCTCTGCCGCTGAACGCAGATACCCTGCACCTGGTCAACGCGGAACTGCTGGCGC<br>TGGTACGTCCGGGCGCACTGCTGGTTAACCCGTGCCGTGGCTCTGTGGTCGACGAAGCCG<br>CAGTTCTGGCGGCTCTGGAACGCGGTCAACTGGGCGGCTACGCGGCGGACGTGTTTGAGA<br>TGGAAGATTGGGCGCGTGCTGATCGTCCGCAGCAGATTGATCCAGCCCTGCTGGCGCACCC<br>GAACACGCTGTTTACTCCGCACATCGGTTCTGCAGTACGTGCGGTTCGTCTGGAGATCGAG<br>CGTTGTGCGGCTCAGAACATTCTGCAGGCGCTGGCGGGCGAGCGTCCGATCAACGCAGTG<br>AATCGCCTGCCGAAAGCCAACCCTGCGGCTGACTCGAGATCTGCAGCTGGTTAA |
| His-Thrombin-SnoopC-PTDH-SpyC | **ATG**GGCAGCAGCCATCATCATCATCATCACAGCAGCGGCCTGGTGCCGCGCGGCAGCCATA<br>TGGGGAGTGGTGGCAGCGGAGGCAAGCCGCTGCGTGGTGCCGTGTTTAGCCTGCAGAAAC<br>AGCATCCCGACTATCCCGATATCTATGGCGCGATTGATCAGAATGGGACCTATCAAAATGTG<br>CGTACCGGCGAAGATGGTAAACTGACCTTTAAGAATCTGAGCGATGGCAAATATCGCCTGTT<br>TGAAAATAGCGAACCCGCTGGCTATAAACCGGTGCAGAATAAGCCGATTGTGGCGTTTCAGA<br>TTGTGAATGGCGAAGTGCGTGATGTGACCAGCATTGTGCCGCAGGATATTCCGGCTACATAT |

| Name | Nucleotide sequence |
|---|---|
| | GAATTTACCAACGGTAAACATTATATCACCAATGAACCGATACCGCCGAAAGGAGGTTCAGG<br>CGGTTCAGGGATGCTGCCGAAACTGGTCATCACCCATCGTGTGCACGAAGAAATCCTGCAA<br>CTGCTGGCTCCGCATTGCGAACTGATTACCAATCAGACCGACAGCACTCTGACCCGTGAAGA<br>AATTCTGCGTCGCTGCCGTGACGCACAGGCAATGATGGCGTTTATGCCTGATCGTGTGGAT<br>GCTGATTTCCTGCAGGCTTGTCCGGAACTGCGCGTAATTGGCTGCGCGCTGAAAGGTTTCG<br>ACAATTTCGACGTTGATGCGTGTACTGCTCGTGGCGTGTGGCTGACCTTCGTTCCGGACCTG<br>CTGACTGTGCCGACTGCGGAACTGGCTATCGGCCTGGCGGTTGGTCTGGGCCGTCACCTG<br>CGTGCAGCAGACGCGTTTGTTCGTTCTGGTAAATTTCGCGGCTGGCAGCCTCGCTTTTACGG<br>TACCGGTCTGGACAACGCGACCGTGGGCTTCCTGGGTATGGGTGCAATCGGTCTGGCGATG<br>GCTGATCGTCTGCAGGGTTGGGGCGCTACCCTGCAATACCACGCGCGTAAAGCTCTGGATA<br>CGCAAACGGAACAACGCCTGGGTCTGCGTCAAGTGGCTTGCAGCGAGCTGTTCGCTTCTAG<br>CGACTTCATTCTGCTGGCTCTGCCGCTGAACGCAGATACCCTGCACCTGGTCAACGCGGAA<br>CTGCTGGCGCTGGTACGTCCGGGCGCACTGCTGGTTAACCCGTGCCGTGGCTCTGTGGTC<br>GACGAAGCCGCAGTTCTGGCGGCTCTGGAACGCGGTCAACTGGGCGGCTACGCGGCGGAC<br>GTGTTTGAGATGGAAGATTGGGCGCGTGCTGATCGTCCGCAGCAGATTGATCCAGCCCTGC<br>TGGCGCACCCGAACACGCTGTTTACTCCGCACATCGGTTCTGCAGTACGTGCGGTTCGTCT<br>GGAGATCGAGCGTTGTGCGGCTCAGAACATTCTGCAGGCGCTGGCGGGCGAGCGTCCGAT<br>CAACGCAGTGAATCGCCTGCCGAAAGCCAACCCTGCGGCTGACTCGAGATCTGCAGCTGGT<br>TCCGGATCAGGATCCGGCGGCGATAGTGCTACCCATATTAAATTCTCAAAACGTGATGAGGA<br>CGGCAAAGAGTTAGCTGGTGCAACTATGGAGTTGCGTGATTCATCTGGTAAAACTATTAGTA<br>CATGGATTTCAGATGGACAAGTGAAAGATTTCTACCTGTATCCAGGAAAATATACATTTGTCG<br>AAACCGCAGCACCAGACGGTTATGAGGTAGCAACTGCTATTACCTTTACAGTTAATGAGCAA<br>GGTCAGGTTACTGTAAATGGCTAA |
| His-Thrombin-SpyT-PTDH-SnoopT | **ATG**GGCAGCAGCCATCATCATCATCATCACAGCAGCGGCCTGGTGCCGCGCGGCAGCCATA<br>TGATGGGCAGCAGCGGCGCCCACATCGTGATGGTGGACGCCTACAAGCCGACGAAGGGTT<br>CAGGGGGATCCGGTATGCTGCCGAAACTGGTCATCACCCATCGTGTGCACGAAGAAATCCT<br>GCAACTGCTGGCTCCGCATTGCGAACTGATTACCAATCAGACCGACAGCACTCTGACCCGT<br>GAAGAAATTCTGCGTCGCTGCCGTGACGCACAGGCAATGATGGCGTTTATGCCTGATCGTGT<br>GGATGCTGATTTCCTGCAGGCTTGTCCGGAACTGCGCGTAATTGGCTGCGCGCTGAAAGGT<br>TTCGACAATTTCGACGTTGATGCGTGTACTGCTCGTGGCGTGTGGCTGACCTTCGTTCCGGA<br>CCTGCTGACTGTGCCGACTGCGGAACTGGCTATCGGCCTGGCGGTTGGTCTGGGCCGTCA<br>CCTGCGTGCAGCAGACGCGTTTGTTCGTTCTGGTAAATTTCGCGGCTGGCAGCCTCGCTTTT<br>ACGGTACCGGTCTGGACAACGCGACCGTGGGCTTCCTGGGTATGGGTGCAATCGGTCTGG<br>CGATGGCTGATCGTCTGCAGGGTTGGGGCGCTACCCTGCAATACCACGCGCGTAAAGCTCT<br>GGATACGCAAACGGAACAACGCCTGGGTCTGCGTCAAGTGGCTTGCAGCGAGCTGTTCGCT<br>TCTAGCGACTTCATTCTGCTGGCTCTGCCGCTGAACGCAGATACCCTGCACCTGGTCAACGC<br>GGAACTGCTGGCGCTGGTACGTCCGGGCGCACTGCTGGTTAACCCGTGCCGTGGCTCTGT<br>GGTCGACGAAGCCGCAGTTCTGGCGGCTCTGGAACGCGGTCAACTGGGCGGCTACGCGGC<br>GGACGTGTTTGAGATGGAAGATTGGGCGCGTGCTGATCGTCCGCAGCAGATTGATCCAGCC |

| Name | Nucleotide sequence |
| --- | --- |
| | CTGCTGGCGCACCCGAACACGCTGTTTACTCCGCACATCGGTTCTGCAGTACGTGCGGTTC<br>GTCTGGAGATCGAGCGTTGTGCGGCTCAGAACATTCTGCAGGCGCTGGCGGGCGAGCGTC<br>CGATCAACGCAGTGAATCGCCTGCCGAAAGCCAACCCTGCGGCTGACTCGAGATCTGCAGC<br>TGGTTCCGGATCAGGATCCGGCGGCAAACTGGGCGATATTGAATTTATTAAAGTGAACAAAT<br>AA |
| His-Thrombin-<br>P450BM3m | **ATG**GGCAGCAGCCATCATCATCATCATCACAGCAGCGGCCTGGTGCCGCGCGGCAGCCATA<br>TGATGACAATTAAAGAAATGCCTCAGCCAAAAACGTTTGGAGAGCTTAAAAATTTACCGTTAT<br>TAAACACAGATAAACCGGTTCAAGCTTTGATGAAAATTGCGGATGAATTAGGAGAAATCTTTA<br>AATTCGAGGCGCCTGGTCGTGTAACGCGCTACTTATCAAGTCAGCGTCTAATTAAAGAAGCA<br>TGCGATGAATCACGCTTTGATAAAAACTTAAGTCAAGGGCTTAAATTTGTACGTGATTTTGCA<br>GGAGACGGGTTAGTTACAAGCTGGACGCATGAAAAAAATTGGAAAAAAGCGCATAATATCTT<br>ACTTCCAAGCTTCAGTCAGCAGGCAATGAAAGGCTATCATGCGATGATGGTCGATATCGCCG<br>TGCAGCTTGTTCAAAAGTGGGAGCGTCTAAATGCAGATGAGCATATTGAAGTACCGGAAGAC<br>ATGACACGTTTAACGCTTGATACAATTGGTCTTTGCGGCTTTAACTATCGCTTTAACAGCTTTT<br>ACCGAGATCAGCCTCATCCATTTATTACAAGTATGGTCCGTGCACTGGATGAAGCAATGAAC<br>AAGCAGCAGCGAGCAAATCCAGACGACCCAGCTTATGATGAAAACAAGCGCCAGTTTCAAGA<br>AGATATCAAGGTGATGAACGACCTAGTAGATAAAATTATTGCAGATCGCAAAGCAAGCGGTG<br>AACAAAGCGATGATTTATTAACGCATATGCTAAACGGAAAAGATCCAGAAACGGGTGAGCCG<br>CTTGATGACGAGAACATTCGCTATCAAATTATTACATTCTTAATTGCGGGACACGAAACAACA<br>AGTGGTCTTTTATCATTTGCGCTGTATTTCTTAGTGAAAAATCCACATGTATTACAAAAAGCAG<br>CAGAAGAAGCAGCACGAGTTCTAGTAGATCCTGTTCCAAGCTACAAACAAGTCAAACAGCTT<br>AAATATGTCGGCATGGTCTTAAACGAAGCGCTGCGCTTATGGCCAACTGCTCCTGCGTTTTC<br>CCTATATGCAAAAGAAGATACGGTGCTTGGAGGAGAATATCCTTTAGAAAAAGGCGACGAAC<br>TAATGGTTCTGATTCCTCAGCTTCACCGTGATAAAACAATTTGGGGAGACGATGTGGAAGAG<br>TTCCGTCCAGAGCGTTTTGAAAATCCAAGTGCGATTCCGCAGCATGCGTTTAAACCGTTTGG<br>AAACGGTCAGCGTGCGTGTATCGGTCAGCAGTTCGCTCTTCATGAAGCAACGCTGGTACTTG<br>GTATGATGCTAAAACACTTTGACTTTGAAGATCATACAAACTACGAGCTGGATATTAAAGAAA<br>CTTTAACGTTAAAACCTGAAGGCTTTGTGGTAAAAGCAAAATCGAAAAAAATTCCGCTTGGCG<br>GAATTCCTTCACCTAGCACTGAACAGTCTGCTAAAAAAGTACGCAAAAAGGCAGAAAACGCT<br>CATAATACGCCGCTGCTTGTGCTATACGGTTCAAATATGGGAACAGCTGAAGGAACGGCGC<br>GTGATTTAGCAGATATTGCAATGAGCAAAGGATTTGCACCGCAGGTCGCAACGCTTGATTCA<br>CACGCCGGAAATCTTCCGCGCGAAGGAGCTGTATTAATTGTAACGGCGTCTTATAACGGTCA<br>TCCGCCTGATAACGCAAAGCAATTTGTCGACTGGTTAGACCAAGCGTCTGCTGATGAAGTAA<br>AAGGCGTTCGCTACTCCGTATTTGGATGCGGCGATAAAAACTGGGCTACTACGTATCAAAAA<br>GTGCCTGCTTTTATCGATGAAACGCTTGCCGCTAAAGGGGCAGAAAACATCGCTGACCGCG |

| Name | Nucleotide sequence |
| --- | --- |
| | GTGAAGCAGATGCAAGCGACGACTTTGAAGGCACATATGAAGAATGGCGTGAACATATGTG<br>GAGTGACGTAGCAGCCTACTTTAACCTCGACATTGAAAACAGTGAAGATAATAAATCTACTCT<br>TTCACTTCAATTTGTCGACAGCGCCGCGGATATGCCGCTTGCGAAAATGCACGGTGCGTTTT<br>CAACGAACGTCGTAGCAAGCAAAGAACTTCAACAGCCAGGCAGTGCACGAAGCACGCGACA<br>TCTTGAAATTGAACTTCCAAAAGAAGCTTCTTATCAAGAAGGAGATCATTTAGGTGTTATTCCT<br>CGCAACTATGAAGGAATAGTAAACCGTGTAACAGCAAGGTTCGGCCTAGATGCATCACAGCA<br>AATCCGTCTGGAAGCAGAAGAAGAAAAATTAGCTCATTTGCCACTCGCTAAAACAGTATCCGT<br>AGAAGAGCTTCTGCAATACGTGGAGCTTCAAGATCCTGTTACGCGCACGCAGCTTCGCGCAA<br>TGGCTGCTAAAACGGTCTGCCCGCCGCATAAAGTAGAGCTTGAAGCCTTGCTTGAAAAGCAA<br>GCCTACAAAGAACAAGTGCTGGCAAAACGTTTAACAATGCTTGAACTGCTTGAAAAATACCC<br>GGCGTGTGAAATGAAATTCAGCGAATTTATCGCCCTTCTGCCAAGCATACGCCCGCGCTATT<br>ACTCGATTTCTTCATCACCTCGTGTCGATGAAAAACAAGCAAGCATCACGGTCAGCGTTGTCT<br>CAGGAGAAGCGTGGAGCGGATATGGAGAATATAAAGGAATTGCGTCGAACTATCTTGCCGA<br>GCTGCAAGAAGGAGATACGATTACGTGCTTTATTTCCACACCGCAGTCAGAATTTACGCTGC<br>CAAAAGACCCTGAAACGCCGCTTATCATGGTCGGACCGGGAACAGGCGTCGCGCCGTTTAG<br>AGGCTTTGTGCAGGCGCGCAAACAGCTAAAAGAACAAGGACAGTCACTTGGAGAAGCACAT<br>TTATACTTCGGCTGCCGTTCACCTCATGAAGACTATCTGTATCAAGAAGAGCTTGAAAACGCC<br>CAAAGCGAAGGCATCATTACGCTTCATACCGCTTTTTCTCGCATGCCAAATCAGCCGAAAAC<br>ATACGTTCAGCACGTAATGGAACAAGACGGCAAGAAATTGATTGAACTTCTTGATCAAGGAG<br>CGCACTTCTATATTTGCGGAGACGGAAGCCAAATGGCACCTGCCGTTGAAGCAACGCTTATG<br>AAAAGCTATGCTGACGTTCACCAAGTGAGTGAAGCAGACGCTCGCTTATGGCTGCAGCAGCT<br>AGAAGAAAAAGGCCGATACGCAAAAGACGTGTGGGCTGGGTAA |
| His-Thrombin-SnoopC-P450BM3m-SpyC | **ATG**GGCAGCAGCCATCATCATCATCATCACAGCAGCGGCCTGGTGCCGCGCGGCAGCCATA<br>TGGGGAGTGGTGGCAGCGGAGGCAAGCCGCTGCGTGGTGCCGTGTTTAGCCTGCAGAAAC<br>AGCATCCCGACTATCCCGATATCTATGGCGCGATTGATCAGAATGGGACCTATCAAAATGTG<br>CGTACCGGCGAAGATGGTAAACTGACCTTTAAGAATCTGAGCGATGGCAAATATCGCCTGTT<br>TGAAAATAGCGAACCCGCTGGCTATAAACCGGTGCAGAATAAGCCGATTGTGGCGTTTCAGA<br>TTGTGAATGGCGAAGTGCGTGATGTGACCAGCATTGTGCCGCAGGATATTCCGGCTACATAT<br>GAATTTACCAACGGTAAACATTATATCACCAATGAACCGATACCGCCGAAAGGAGGTTCAGG<br>CGGTTCAGGGATGACAATTAAAGAAATGCCTCAGCCAAAAACGTTTGGAGAGCTTAAAAATTT<br>ACCGTTATTAAACACAGATAAACCGGTTCAAGCTTTGATGAAAATTGCGGATGAATTAGGAGA<br>AATCTTTAAATTCGAGGCGCCTGGTCGTGTAACGCGCTACTTATCAAGTCAGCGTCTAATTAA<br>AGAAGCATGCGATGAATCACGCTTTGATAAAAACTTAAGTCAAGGGCTTAAATTTGTACGTGA<br>TTTTGCAGGAGACGGGTTAGTTACAAGCTGGACGCATGAAAAAAATTGGAAAAAAGCGCATA<br>ATATCTTACTTCCAAGCTTCAGTCAGCAGGCAATGAAAGGCTATCATGCGATGATGGTCGATA<br>TCGCCGTGCAGCTTGTTCAAAAGTGGGAGCGTCTAAATGCAGATGAGCATATTGAAGTACCG<br>GAAGACATGACACGTTTAACGCTTGATACAATTGGTCTTTGCGGCTTTAACTATCGCTTTAAC<br>AGCTTTTACCGAGATCAGCCTCATCCATTTATTACAAGTATGGTCCGTGCACTGGATGAAGCA<br>ATGAACAAGCAGCAGCGAGCAAATCCAGACGACCCAGCTTATGATGAAAACAAGCGCCAGTT |

| Name | Nucleotide sequence |
| --- | --- |
| | TCAAGAAGATATCAAGGTGATGAACGACCTAGTAGATAAAATTATTGCAGATCGCAAAGCAAG<br>CGGTGAACAAAGCGATGATTTATTAACGCATATGCTAAACGGAAAAGATCCAGAAACGGGTG<br>AGCCGCTTGATGACGAGAACATTCGCTATCAAATTATTACATTCTTAATTGCGGGACACGAAA<br>CAACAAGTGGTCTTTTATCATTTGCGCTGTATTTCTTAGTGAAAAATCCACATGTATTACAAAA<br>AGCAGCAGAAGAAGCAGCACGAGTTCTAGTAGATCCTGTTCCAAGCTACAAACAAGTCAAAC<br>AGCTTAAATATGTCGGCATGGTCTTAAACGAAGCGCTGCGCTTATGGCCAACTGCTCCTGCG<br>TTTTCCCTATATGCAAAAGAAGATACGGTGCTTGGAGGAGAATATCCTTTAGAAAAAGGCGAC<br>GAACTAATGGTTCTGATTCCTCAGCTTCACCGTGATAAAACAATTTGGGGAGACGATGTGGA<br>AGAGTTCCGTCCAGAGCGTTTTGAAAATCCAAGTGCGATTCCGCAGCATGCGTTTAAACCGT<br>TTGGAAACGGTCAGCGTGCGTGTATCGGTCAGCAGTTCGCTCTTCATGAAGCAACGCTGGTA<br>CTTGGTATGATGCTAAAACACTTTGACTTTGAAGATCATACAAACTACGAGCTGGATATTAAA<br>GAAACTTTAACGTTAAAACCTGAAGGCTTTGTGGTAAAAGCAAAATCGAAAAAAATTCCGCTT<br>GGCGGAATTCCTTCACCTAGCACTGAACAGTCTGCTAAAAAAGTACGCAAAAAGGCAGAAAA<br>CGCTCATAATACGCCGCTGCTTGTGCTATACGGTTCAAATATGGGAACAGCTGAAGGAACGG<br>CGCGTGATTTAGCAGATATTGCAATGAGCAAAGGATTTGCACCGCAGGTCGCAACGCTTGAT<br>TCACACGCCGGAAATCTTCCGCGCGAAGGAGCTGTATTAATTGTAACGGCGTCTTATAACGG<br>TCATCCGCCTGATAACGCAAAGCAATTTGTCGACTGGTTAGACCAAGCGTCTGCTGATGAAG<br>TAAAAGGCGTTCGCTACTCCGTATTTGGATGCGGCGATAAAAACTGGGCTACTACGTATCAA<br>AAAGTGCCTGCTTTTATCGATGAAACGCTTGCCGCTAAAGGGGCAGAAAACATCGCTGACCG<br>CGGTGAAGCAGATGCAAGCGACGACTTTGAAGGCACATATGAAGAATGGCGTGAACATATGT<br>GGAGTGACGTAGCAGCCTACTTTAACCTCGACATTGAAAACAGTGAAGATAATAAATCTACTC<br>TTTCACTTCAATTTGTCGACAGCGCCGCGGATATGCCGCTTGCGAAAATGCACGGTGCGTTT<br>TCAACGAACGTCGTAGCAAGCAAAGAACTTCAACAGCCAGGCAGTGCACGAAGCACGCGAC<br>ATCTTGAAATTGAACTTCCAAAAGAAGCTTCTTATCAAGAAGGAGATCATTTAGGTGTTATTCC<br>TCGCAACTATGAAGGAATAGTAAACCGTGTAACAGCAAGGTTCGGCCTAGATGCATCACAGC<br>AAATCCGTCTGGAAGCAGAAGAAGAAAAATTAGCTCATTTGCCACTCGCTAAAACAGTATCC<br>GTAGAAGAGCTTCTGCAATACGTGGAGCTTCAAGATCCTGTTACGCGCACGCAGCTTCGCG<br>CAATGGCTGCTAAAACGGTCTGCCCGCCGCATAAAGTAGAGCTTGAAGCCTTGCTTGAAAAG<br>CAAGCCTACAAAGAACAAGTGCTGGCAAAACGTTTAACAATGCTTGAACTGCTTGAAAAATAC<br>CCGGCGTGTGAAATGAAATTCAGCGAATTTATCGCCCTTCTGCCAAGCATACGCCCGCGCTA<br>TTACTCGATTTCTTCATCACCTCGTGTCGATGAAAAACAAGCAAGCATCACGGTCAGCGTTGT<br>CTCAGGAGAAGCGTGGAGCGGATATGGAGAATATAAAGGAATTGCGTCGAACTATCTTGCC<br>GAGCTGCAAGAAGGAGATACGATTACGTGCTTTATTTCCACACCGCAGTCAGAATTTACGCT<br>GCCAAAAGACCCTGAAACGCCGCTTATCATGGTCGGACCGGGAACAGGCGTCGCGCCGTTT<br>AGAGGCTTTGTGCAGGCGCGCAAACAGCTAAAAGAACAAGGACAGTCACTTGGAGAAGCAC<br>ATTTATACTTCGGCTGCCGTTCACCTCATGAAGACTATCTGTATCAAGAAGAGCTTGAAAACG<br>CCCAAAGCGAAGGCATCATTACGCTTCATACCGCTTTTTCTCGCATGCCAAATCAGCCGAAA<br>ACATACGTTCAGCACGTAATGGAACAAGACGGCAAGAAATTGATTGAACTTCTTGATCAAGG<br>AGCGCACTTCTATATTTGCGGAGACGGAAGCCAAATGGCACCTGCCGTTGAAGCAACGCTTA<br>TGAAAAGCTATGCTGACGTTCACCAAGTGAGTGAAGCAGACGCTCGCTTATGGCTGCAGCA |

| Name | Nucleotide sequence |
|---|---|
| | GCTAGAAGAAAAAGGCCGATACGCAAAAGACGTGTGGGCTGGGTCCGGATCAGGATCCGG<br>CGGCGATAGTGCTACCCATATTAAATTCTCAAAACGTGATGAGGACGGCAAAGAGTTAGCTG<br>GTGCAACTATGGAGTTGCGTGATTCATCTGGTAAAACTATTAGTACATGGATTTCAGATGGAC<br>AAGTGAAAGATTTCTACCTGTATCCAGGAAAATATACATTTGTCGAAACCGCAGCACCAGACG<br>GTTATGAGGTAGCAACTGCTATTACCTTTACAGTTAATGAGCAAGGTCAGGTTACTGTAAATG<br>GCTAA |
| His-Thrombin-SpyT-P450BM3m-SnoopT | **ATG**GGCAGCAGCCATCATCATCATCATCACAGCAGCGGCCTGGTGCCGCGCGGCAGCCATA<br>TGATGGGCAGCAGCGGCGCCCACATCGTGATGGTGGACGCCTACAAGCCGACGAAGGGTT<br>CAGGGGGATCCGGTATGACAATTAAAGAAATGCCTCAGCCAAAAACGTTTGGAGAGCTTAAA<br>AATTTACCGTTATTAAACACAGATAAACCGGTTCAAGCTTTGATGAAAATTGCGGATGAATTA<br>GGAGAAATCTTTAAATTCGAGGCGCCTGGTCGTGTAACGCGCTACTTATCAAGTCAGCGTCT<br>AATTAAAGAAGCATGCGATGAATCACGCTTTGATAAAAACTTAAGTCAAGGGCTTAAATTTGT<br>ACGTGATTTTGCAGGAGACGGGTTAGTTACAAGCTGGACGCATGAAAAAAATTGGAAAAAAG<br>CGCATAATATCTTACTTCCAAGCTTCAGTCAGCAGGCAATGAAAGGCTATCATGCGATGATG<br>GTCGATATCGCCGTGCAGCTTGTTCAAAAGTGGGAGCGTCTAAATGCAGATGAGCATATTGA<br>AGTACCGGAAGACATGACACGTTTAACGCTTGATACAATTGGTCTTTGCGGCTTTAACTATCG<br>CTTTAACAGCTTTTACCGAGATCAGCCTCATCCATTTATTACAAGTATGGTCCGTGCACTGGA<br>TGAAGCAATGAACAAGCAGCAGCGAGCAAATCCAGACGACCCAGCTTATGATGAAAACAAGC<br>GCCAGTTTCAAGAAGATATCAAGGTGATGAACGACCTAGTAGATAAAATTATTGCAGATCGCA<br>AAGCAAGCGGTGAACAAAGCGATGATTTATTAACGCATATGCTAAACGGAAAAGATCCAGAA<br>ACGGGTGAGCCGCTTGATGACGAGAACATTCGCTATCAAATTATTACATTCTTAATTGCGGGA<br>CACGAAACAACAAGTGGTCTTTTATCATTTGCGCTGTATTTCTTAGTGAAAAATCCACATGTAT<br>TACAAAAAGCAGCAGAAGAAGCAGCACGAGTTCTAGTAGATCCTGTTCCAAGCTACAAACAA<br>GTCAAACAGCTTAAATATGTCGGCATGGTCTTAAACGAAGCGCTGCGCTTATGGCCAACTGC<br>TCCTGCGTTTTCCCTATATGCAAAAGAAGATACGGTGCTTGGAGGAGAATATCCTTTAGAAAA<br>AGGCGACGAACTAATGGTTCTGATTCCTCAGCTTCACCGTGATAAAACAATTTGGGGAGACG<br>ATGTGGAAGAGTTCCGTCCAGAGCGTTTTGAAAATCCAAGTGCGATTCCGCAGCATGCGTTT<br>AAACCGTTTGGAAACGGTCAGCGTGCGTGTATCGGTCAGCAGTTCGCTCTTCATGAAGCAAC<br>GCTGGTACTTGGTATGATGCTAAAACACTTTGACTTTGAAGATCATACAAACTACGAGCTGGA<br>TATTAAAGAAACTTTAACGTTAAAACCTGAAGGCTTTGTGGTAAAAGCAAAATCGAAAAAAATT<br>CCGCTTGGCGGAATTCCTTCACCTAGCACTGAACAGTCTGCTAAAAAAGTACGCAAAAAGGC<br>AGAAAACGCTCATAATACGCCGCTGCTTGTGCTATACGGTTCAAATATGGGAACAGCTGAAG<br>GAACGGCGCGTGATTTAGCAGATATTGCAATGAGCAAAGGATTTGCACCGCAGGTCGCAAC<br>GCTTGATTCACACGCCGGAAATCTTCCGCGCGAAGGAGCTGTATTAATTGTAACGGCGTCTT<br>ATAACGGTCATCCGCCTGATAACGCAAAGCAATTTGTCGACTGGTTAGACCAAGCGTCTGCT<br>GATGAAGTAAAAGGCGTTCGCTACTCCGTATTTGGATGCGGCGATAAAAACTGGGCTACTAC<br>GTATCAAAAAGTGCCTGCTTTTATCGATGAAACGCTTGCCGCTAAAGGGGCAGAAAACATCG<br>CTGACCGCGGTGAAGCAGATGCAAGCGACGACTTTGAAGGCACATATGAAGAATGGCGTGA<br>ACATATGTGGAGTGACGTAGCAGCCTACTTTAACCTCGACATTGAAAACAGTGAAGATAATAA |

| Name | Nucleotide sequence |
| --- | --- |
| | ATCTACTCTTTCACTTCAATTTGTCGACAGCGCCGCGGATATGCCGCTTGCGAAAATGCACG<br>GTGCGTTTTCAACGAACGTCGTAGCAAGCAAAGAACTTCAACAGCCAGGCAGTGCACGAAG<br>CACGCGACATCTTGAAATTGAACTTCCAAAAGAAGCTTCTTATCAAGAAGGAGATCATTTAGG<br>TGTTATTCCTCGCAACTATGAAGGAATAGTAAACCGTGTAACAGCAAGGTTCGGCCTAGATG<br>CATCACAGCAAATCCGTCTGGAAGCAGAAGAAGAAAAATTAGCTCATTTGCCACTCGCTAAA<br>ACAGTATCCGTAGAAGAGCTTCTGCAATACGTGGAGCTTCAAGATCCTGTTACGCGCACGCA<br>GCTTCGCGCAATGGCTGCTAAAACGGTCTGCCCGCCGCATAAAGTAGAGCTTGAAGCCTTG<br>CTTGAAAAGCAAGCCTACAAAGAACAAGTGCTGGCAAAACGTTTAACAATGCTTGAACTGCTT<br>GAAAAATACCCGGCGTGTGAAATGAAATTCAGCGAATTTATCGCCCTTCTGCCAAGCATACG<br>CCCGCGCTATTACTCGATTTCTTCATCACCTCGTGTCGATGAAAAACAAGCAAGCATCACGG<br>TCAGCGTTGTCTCAGGAGAAGCGTGGAGCGGATATGGAGAATATAAAGGAATTGCGTCGAA<br>CTATCTTGCCGAGCTGCAAGAAGGAGATACGATTACGTGCTTTATTTCCACACCGCAGTCAG<br>AATTTACGCTGCCAAAAGACCCTGAAACGCCGCTTATCATGGTCGGACCGGGAACAGGCGT<br>CGCGCCGTTTAGAGGCTTTGTGCAGGCGCGCAAACAGCTAAAAGAACAAGGACAGTCACTT<br>GGAGAAGCACATTTATACTTCGGCTGCCGTTCACCTCATGAAGACTATCTGTATCAAGAAGA<br>GCTTGAAAACGCCCAAAGCGAAGGCATCATTACGCTTCATACCGCTTTTTCTCGCATGCCAA<br>ATCAGCCGAAAACATACGTTCAGCACGTAATGGAACAAGACGGCAAGAAATTGATTGAACTT<br>CTTGATCAAGGAGCGCACTTCTATATTTGCGGAGACGGAAGCCAAATGGCACCTGCCGTTGA<br>AGCAACGCTTATGAAAAGCTATGCTGACGTTCACCAAGTGAGTGAAGCAGACGCTCGCTTAT<br>GGCTGCAGCAGCTAGAAGAAAAAGGCCGATACGCAAAAGACGTGTGGGCTGGGTCCGGAT<br>CAGGATCCGGCGGCAAACTGGGCGATATTGAATTTATTAAAGTGAACAAATAA |
| Representative artificial operon for hybrid scaffolds : Upstream-RBSA-His-EutM-RBSA-SnoopT-EutM-SpyT-Downstream | ATCATAAAAAATTTATTTGCTTTGTGAGCGGATAACAATTATAATAGATTCAACAAACAGACAA<br>TCTGGTCTGTTTGTATTATGAACGAAGGAGGGATCTGGATCC**ATG**CATCATCATCACCACCA<br>CGGTTCTGGTTCTGGTTCTGGTTCTGGTTCTGGTTCTGAAGCATTAGGAATGATTGAAACCC<br>GGGGCCTGGTTGCGCTGATTGAGGCCTCCGATGCGATGGTAAAAGCCGCGCGCGTGAAGC<br>TGGTCGGCGTGAAGCAGATTGGCGGTGGCCTGTGTACTGCCATGGTGCGTGGCGATGTGG<br>CGGCGTGCAAAGCCGCAACCGATGCTGGCGCCGCTGCGGCGCAGCGCATTGGCGAGTTGG<br>TCTCCGTACACGTGATTCCACGCCCGCACGGCGATCTGGAAGAAGTGTTCCCGATCAGCTT<br>CAAAGGCGACAGCAACATTTGAATGAACGAAGGAGGGATCTGGATCC**ATG**AAACTGGGCGA<br>TATTGAATTTATTAAAGTGAACAAAGGCAGCGGTAGCGGCAGCGGTAGCGGCAGCGGTAGC<br>ATGGAAGCATTAGGAATGATTGAAACCCGGGGCCTGGTTGCGCTGATTGAGGCCTCCGATG<br>CGATGGTAAAAGCCGCGCGCGTGAAGCTGGTCGGCGTGAAGCAGATTGGCGGTGGCCTGT<br>GTACTGCCATGGTGCGTGGCGATGTGGCGGCGTGCAAAGCCGCAACCGATGCTGGCGCCG<br>CTGCGGCGCAGCGCATTGGCGAGTTGGTCTCCGTACACGTGATTCCACGCCCGCACGGCG<br>ATCTGGAAGAAGTGTTCCCGATCAGCTTCAAAGGCGACAGCAACATTGGATCCGGTGGAGG<br>GTCTGGTGGCGGATCAGGAAGTGCCCACATCGTGATGGTGGACGCCTACAAGCCGACGAA<br>GTAAGCGGCCGCCTCGAGGCCCAAGGTTTAAAGCCTTGGGCTTTTCTGTTTACTAGTTAACT<br>TCAGGACCTGCAGATAAAAAAAAT |

| Name | Nucleotide sequence |
|---|---|
| Representative expression cassette for cargo proteins : Upstream-RBS-His-Thrombin-SpyT-GFP-SnoopT-Downstream | TAATACGACTCACTATAGGGGAATTGTGAGCGGATAACAATTCCCCTCTAGAAATAATTTTGTTTAACTTTAAGAAGGAGATATACC**ATG**GGCAGCAGCCATCATCATCATCATCACAGCAGCGGCCTGGTGCCGCGCGGCAGCCATATGATGGGCAGCAGCGGCGCCCACATCGTGATGGTGGACGCCTACAAGCCGACGAAGGGTTCAGGGGGATCCGGTATGTCAAAAGGAGAAGAACTTTTTACAGGTGTAGTACCTATCTTGGTTGAATTGGATGGTGATGTTAACGGTCACAAATTTTCTGTACGTGGTGAAGGTGAAGGTGATGCAACTAACGGTAAATTGACACTTAAATTCATTTGTACAACTGGAAAACTTCCTGTTCCTTGGCCTACTCTTGTTACAACATTGACATATGGAGTACAATGTTTTTCACGTTATCCTGATCATATGAAACGTCACGATTTTTTTAAATCTGCTATGCCAGAAGGTTATGTACAAGAACGTACAATTTCATTTAAAGATGACGGAACATATAAAACACGTGCTGAAGTAAAATTCGAAGGTGACACTCTTGTTAATCGTATCGAATTGAAAGGAATCGATTTCAAAGAAGATGGTAACATTTTGGGACACAAACTTGAATACAACTTCAACTCTCATAATGTTTATATCACAGCTGACAAACAAAAAAACGGTATTAAAGCTAATTTTAAAATTCGTCACAATGTTGAAGATGGATCTGTTCAATTGGCTGATCATTATCAACAAAATACACCAATCGGAGACGGACCAGTATTGCTTCCAGATAACCACTACCTTTCTACTCAATCAGTTCTTTCAAAAGATCCTAACGAAAAACGTGACCATATGGTACTTCTTGAATTTGTTACAGCAGCAGGTATCACTCACGGTATGGACGAACTTTATAAATCCGGATCAGGATCCGGCGGCAAACTGGGCGATATTGAATTTATTAAAGTGAACAAATAAGCGGCCGCACTCGAGCACCACCACCACCACCACTGAGATCCGGCTGCTAACAAAGCCCGAAAGGAAGCTGAGTTGGCTGCTGCCACCGCTGAGCAATAACTA |

### Supplementary References

1 Xia, Y. *et al.* T5 exonuclease-dependent assembly offers a low-cost method for efficient cloning and site-directed mutagenesis. *Nucleic Acids Research* **47**, e15-e15, doi:10.1093/nar/gky1169 (2018).

2 Schmidt-Dannert, S., Zhang, G., Johnston, T., Quin, M. B. & Schmidt-Dannert, C. Building a toolbox of protein scaffolds for future immobilization of biocatalysts. *Appl Microbiol Biotechnol* **102**, 8373-8388, doi:10.1007/s00253-018-9252-6 (2018).

3 Seo, S. O. & Schmidt-Dannert, C. Development of a synthetic cumate-inducible gene expression system for *Bacillus*. *Appl Microbiol Biotechnol* **103**, 303-313, doi:10.1007/s00253-018-9485-4 (2019).

4 Zhang, G., Quin, M. B. & Schmidt-Dannert, C. Self-assembling protein scaffold system for easy *in vitro* coimmobilization of biocatalytic cascade enzymes. *Acs Catal* **8**, 5611-5620, doi:10.1021/acscatal.8b00986 (2018).

5 Li, Q.-S., Schwaneberg, U., Fischer, P. & Schmid, R. D. Directed evolution of the fatty-acid hydroxylase P450 BM-3 into an indole-hydroxylating catalyst. *Chemistry – A European Journal* **6**, 1531-1536, doi:10.1002/(SICI)1521-3765(20000502)6:9<1531::AID-CHEM1531>3.0.CO;2-D (2000).

6 Zhang, G., Johnston, T., Quin, M. B. & Schmidt-Dannert, C. Developing a protein scaffolding system for rapid enzyme immobilization and optimization of enzyme functions for biocatalysis. *ACS Synthetic Biology* **8**, 1867-1876, doi:10.1021/acssynbio.9b00187 (2019).

7 Salis, H. M., Mirsky, E. A. & Voigt, C. A. Automated design of synthetic ribosome binding sites to control protein expression. *Nat Biotechnol* **27**, 946-950, doi:10.1038/nbt.1568 (2009).

8 Studier, F. W. Protein production by auto-induction in high density shaking cultures. *Protein expression and purification* **41**, 207-234, doi:10.1016/j.pep.2005.01.016 (2005).

9 Starr lab. *Quantification of protein bands using densitometry*, <https://sites.google.com/a/umn.edu/starrlab/protocols/quantitation/quantification-of-protein-bands-using-densitometry> .

10 Omura, T. & Sato, R. The carbon monoxide-binding pigment of liver microsomes. *Journal of Biological Chemistry* **239**, 2379-2385, doi:10.1016/s0021-9258(20)82245-5 (1964).

11 Beyer, N. *et al.* P450(BM3) fused to phosphite dehydrogenase allows phosphite-driven selective oxidations. *Appl Microbiol Biotechnol* **101**, 2319-2331, doi:10.1007/s00253-016-7993-7 (2017).

12 Ebert, M. C. C. J. C., Guzman Espinola, J., Lamoureux, G. & Pelletier, J. N. Substrate-specific screening for mutational hotspots using biased molecular dynamics simulations. *Acs Catal* **7**, 6786-6797, doi:10.1021/acscatal.7b02634 (2017).

13 Tooker, B. C., Kandel, S. E., Work, H. M. & Lampe, J. N. Pseudomonas aeruginosa cytochrome P450 CYP168A1 is a fatty acid hydroxylase that metabolizes arachidonic acid to the vasodilator 19-HETE. *Journal of Biological Chemistry* **298**, 101629, doi:10.1016/j.jbc.2022.101629 (2022).

14 Vick, J. E. *et al.* Optimized compatible set of BioBrick vectors for metabolic pathway engineering. *Appl Microbiol Biotechnol* **92**, 1275-1286, doi:10.1007/s00253-011-3633-4 (2011).
